# Leveraging chromatin packing domains to target chemoevasion *in vivo*

**DOI:** 10.1101/2024.11.14.623612

**Authors:** Jane Frederick, Ranya K. A. Virk, I Chae Ye, Luay M. Almassalha, Greta M. Wodarcyk, David VanDerway, Paola Carrillo Gonzalez, Rikkert J. Nap, Vasundhara Agrawal, Nicholas M. Anthony, John Carinato, Wing Shun Li, Cody L. Dunton, Karla I. Medina, Rivaan Kakkaramadam, Surbhi Jain, Shohreh Shahabi, Guillermo Ameer, Igal G. Szleifer, Vadim Backman

**Affiliations:** Department of Biomedical Engineering, Northwestern University, Evanston, IL 60208, USA; Center for Physical Genomics and Engineering, Northwestern University, Evanston, IL 60208, USA; Chemistry of Life Processes Institute, Northwestern University, Evanston, IL 60208, USA; Department of Pharmaceutical Chemistry, University of California, San Francisco, San Francisco, CA 94158, USA; Department of Gastroenterology and Hepatology, Feinberg School of Medicine, Northwestern University, Chicago, IL 60611, USA; Department of Obstetrics and Gynecology, Prentice Women’s Hospital, Feinberg School of Medicine, Northwestern University, Chicago, IL 60611, USA; Department of Chemistry, Northwestern University, Evanston, IL 60208, USA

**Keywords:** Chromatin, Cancer, Chemotherapy, Plasticity, Biophysics

## Abstract

Cancer cells exhibit a remarkable resilience to cytotoxic stress, often adapting through transcriptional changes linked to alterations in chromatin structure. In several types of cancer, these adaptations involve epigenetic modifications and restructuring of topologically associating domains (TADs). However, the underlying principles by which chromatin architecture facilitates such adaptability across different cancers remain poorly understood. To investigate the role of chromatin in this process, we developed a physics-based mechanistic model that connects chromatin organization to cell fate decisions, specifically survival following chemotherapy. Our model builds on the observation that chromatin forms packing domains, which influence transcriptional efficiency through macromolecular crowding. The model accurately predicts chemoevasion *in vitro*, suggesting that changes in packing domains affect the likelihood of survival. Consistent results across diverse cancer types indicate that the model captures fundamental principles of chromatin-mediated adaptation, independent of the specific cancer or chemotherapy mechanisms involved. Based on these insights, we hypothesized that compounds capable of modulating packing domains, termed Transcriptional Plasticity Regulators (TPRs), could prevent cellular adaptation to chemotherapy. Using live-cell chromatin imaging, we conducted a compound screen that identified several TPRs which synergistically enhanced chemotherapy-induced cell death. The most effective TPR significantly improved therapeutic outcomes in a patient-derived xenograft (PDX) model of ovarian cancer. These findings underscore the central role of chromatin in cellular adaptation to cytotoxic stress and present a novel framework for enhancing cancer therapies, with broad potential across multiple cancer types.

## Introduction

Chemotherapy resistance, driven by the ability of cancer cells to rapidly adapt to cytotoxic stress, remains a major obstacle in cancer treatment. While genetic mutations have long been recognized as a major factor in resistance development (1– 4), the timescale of these mutations far exceeds the rapid cell fate decisions that follow chemotherapy exposure. Typically, mutations accumulate over months or years (5), yet cancer cells must make critical survival decisions within hours of chemotherapy administration (6). This temporal gap underscores the importance of non-genetic mechanisms in shaping immediate therapeutic responses.

Among these, chromatin-mediated transcriptional plasticity has emerged as a pivotal driver of cellular adaptation (7, 8). Defined as the ability of cells to dynamically alter gene expression in response to environmental cues, transcriptional plasticity is intricately influenced by the hierarchical organization of chromatin. This structure spans multiple length scales, from nucleosomes to topologically associating domains (TADs) to larger chromatin compartments. Alterations in chromatin organization are known to play a role in oncogene activation and chemoresistance. For instance, disruptions at TAD boundaries can facilitate enhancer hijacking, while large-scale compartmental shifts drive transcriptional reprogramming (9–11). However, recent studies reveal that changes in TAD structure do not universally lead to gene expression changes (12, 13). Additionally, epigenetic reprogramming and shifts in chromatin accessibility are associated with dedifferentiation, heightened drug resistance, and poor treatment outcomes (14–17). Yet, therapies targeting epigenetic mechanisms and chromatin topology show positive responses in only a small subset of patients (18). These findings underscore the need for a deeper understanding of the regulatory mechanisms by which chromatin influences transcription. For example, how does chromatin remodeling facilitate rapid transcriptional activation in response to chemotherapy? Unraveling this complexity is essential for understanding the processes that enable rapid cellular adaptation and, ultimately, therapy resistance.

To address this gap, we propose that chromatin packing domains – nanoscale structures smaller than TADs that are characterized by mass fractal properties – are a key regulatory unit in transcriptional plasticity and chemotherapy resistance. Packing domains are densely packed regions of chromatin ranging from 60 to 90 nm in radius and containing 80-200 kbp of genomic material (19, 20). Given their high surface area-to-volume ratio, these domains enable efficient DNA packaging while maintaining accessibility for rapid transcriptional changes, a crucial feature for cells under chemotherapy-induced stress. The dense packing of nuclear components within packing domains alters the local environment, influencing the diffusion and binding kinetics of transcription factors, RNA polymerase, and other regulatory proteins (21, 22). This macromolecular crowding can facilitate or inhibit protein-DNA interactions depending on the specific chromatin context, providing a potential mechanism for modulating gene expression and enabling cancer cells to swiftly reprogram their transcriptomes in response to chemotherapeutic stress.

To investigate the role of packing domains in therapy resistance, we developed a Chromatin-Dependent Adaptability (CDA) model that integrates the effects of packing domains on transcription. This model effectively captures how the biophysical properties of packing domains modulate transcriptional plasticity in response to cytotoxic stress. Specifically, it utilizes the average scaling behavior of packing domains (*D*_*n*_) as a quantitative measure of the distribution of DNA density, which influences the transcriptional response to chemotherapy through chromatin-mediated crowding effects. Using Partial Wave Spectroscopic (PWS) microscopy, a label-free imaging technique (23), we validate our model by tracking real-time changes to overall nuclear chromatin organization in live cells. Based on our model, we hypothesized that compounds that modulate chromatin packing domains will influence the response to chemotherapy. Therefore, we explore the therapeutic potential of Transcriptional Plasticity Regulators (TPRs), a novel class of compounds that enhance chemotherapy efficacy by modulating chromatin conformation. TPRs target the fractal organization of packing domains, disrupting their spatial arrangement to alter gene accessibility. In combination with standard chemotherapy, TPRs significantly increased cell death *in vitro* and improved treatment outcomes in a patient-derived xenograft (PDX) model, showcasing their potential as a novel therapeutic strategy. By modulating the structure of packing domains, TPRs limit the ability of cancer cells to rewire transcriptional programs that promote survival, further highlighting their potential to inhibit chromatin-mediated adaptability. These findings establish a novel framework for understanding how targeting chromatin-dependent transcriptional plasticity can provide new strategies to combat therapy resistance across cancer types and treatment modalities.

## Results

### Developing a mechanistic model of chromatin-mediated cell survival

To investigate how chromatin structure influences cell survival, we applied a two-step computational modeling approach (Fig. 1A). Using our previously developed Chromatin Packing Macromolecular Crowding (CPMC) model to link chromatin structure to gene transcription (24, 25), we examine how variations in chromatin architecture (captured by the scaling exponent *D*) influence gene activation. Here, gene activation is measured by the transcript ratio *x*, which is the ratio of mRNA levels after versus before chemotherapy (*x* = *N*_2_*/N*_1_; Fig. 1A). Our new Chromatin-Dependent Adaptability (CDA) model further proposes that cells must reach a critical gene activation threshold, *x*_crit_, to survive cytotoxic stress. In this framework, cells with higher gene activity are more likely to withstand cell death (Fig. 1A). This concept aligns with findings by Paek et al., who observed that surpassing a threshold level of p53 accumulation triggers apoptosis (26). While the CPMC model initially addressed the role of chromatin in transcriptional response, the CDA model advances this by defining a gene activation threshold essential for survival under stress. Thus, the CDA model integrates established principles of the impact of chromatin on gene expression to provide a unified framework for understanding cellular adaptation to cytotoxic challenges.

**Fig. 1.**
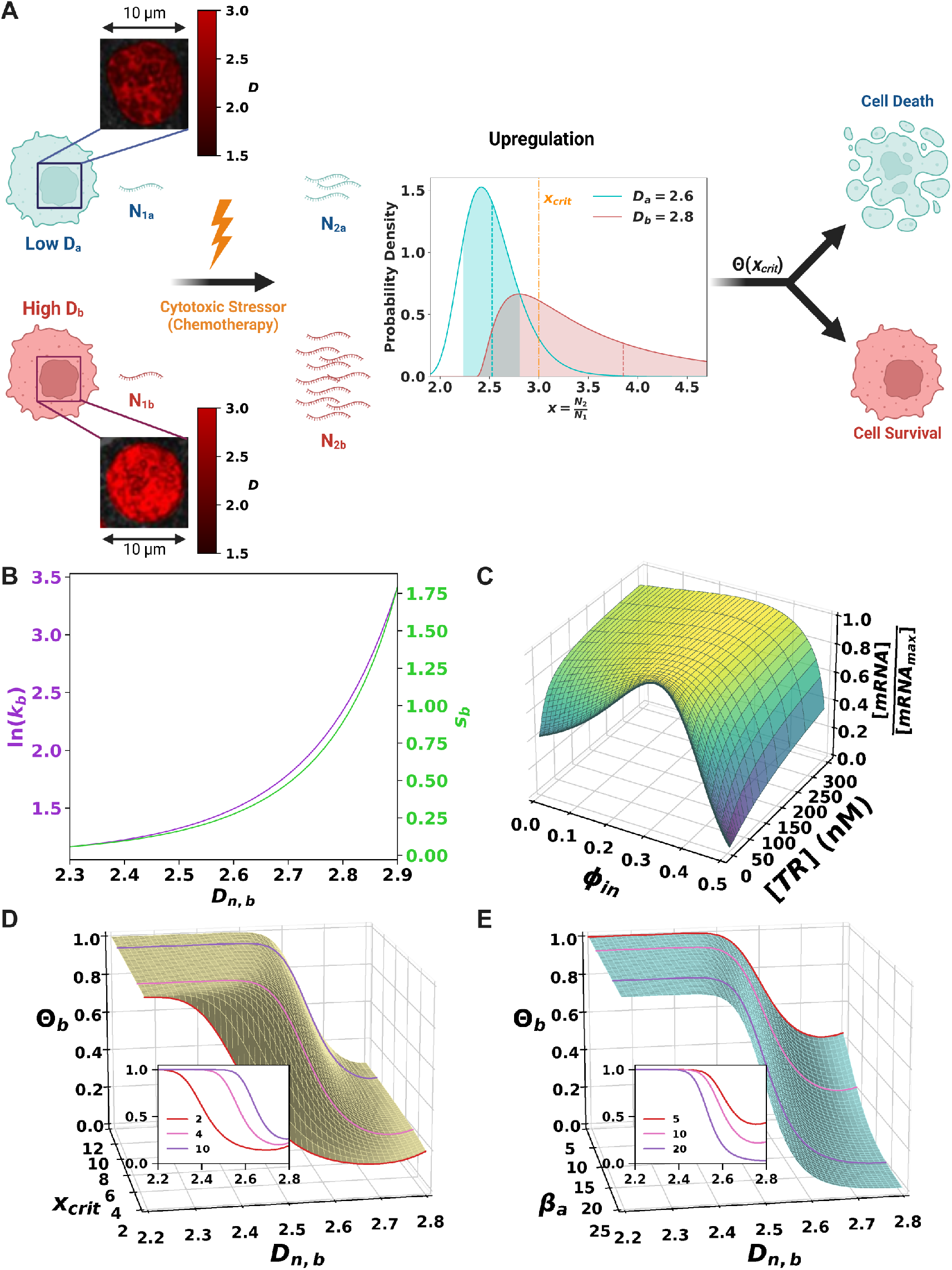
Linking chromatin to transcriptional plasticity and cell survival under cytotoxic stress using the Chromatin Dependent Adaptability (CDA) model. (A) Schematic illustrating the differential responses of cells with varying nuclear chromatin organization *D*_*n*_ to cytotoxic stress. PWS microscopy images show representative low and high *D*_*n*_ cells. The probability distribution function (PDF) of gene upregulation (*x* = *N*_2_ */N*_1_) is shown for both cell types, demonstrating increased mean and standard deviation of upregulated transcripts in high *D*_*n*_ cells, leading to a higher survival probability. (B) Quantitative relationship between *D*_*n*_, transcriptional malleability (*k*, purple), and transcriptional heterogeneity (*s*, green) for cell *b* with fixed parameters ln(*E*/Ē) = 0, *β*_*a*_ = 10, and *t* = 7 hours. (C) 3D plot showing the effect of transcriptional reactant (TR) concentration and local crowding (*ϕ*_in_) on the relative amount of mRNA produced. (D) Cell death probability (Θ_*b*_) as a function of *D*_*n*_ for varying upregulation thresholds (*x*_crit_) with fixed parameters ln(*E*/Ē) = −2, *β*_*a*_ = 10, and *t* = 7 hours. Inset shows individual curves for select *x*_crit_ values. (E) Cell death probability (Θ) as a function of *D*_*n*_ for different transcriptional amplification levels (*β*_*a*_) with fixed parameters ln(*E*/Ē) = −2, *x*_crit_ = 5, and *t* = 7 hours. Inset displays individual curves for select *β*_*a*_ values.

In the first part of our framework, we investigate how the transcriptional activity of a gene within a nanoscale chromatin packing domain (PD) is influenced by macromolecular crowding and the availability of transcriptional machinery. Packing domains are densely packed, mass-fractal structures that exhibit a power-law relationship between DNA quantity (*N*_PD_, in base pairs) and the occupied three-dimensional space (radius, *r*_PD_) following 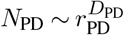, where *D*_PD_ is the scaling exponent describing DNA packing within the domain (19, 20). This compact organization affects biochemical kinetics by modulating macromolecular crowding, thus influencing the mobility and binding of transcription factors and RNA polymerase II (21, 22, 27). We introduce a local crowding metric, *ϕ*_in_, which quantifies the percentage of the transcriptional interaction volume occupied by crowders; previous studies show that *ϕ*_in_ has a non-monotonic relationship with mRNA synthesis (21, 22). This suggests that packing domains regulate transcription via their dense, heterochromatic cores and more open, euchromatic peripheries (28, 29), creating crowding gradients that affect protein diffusion and activity. Genes positioned at the periphery of a domain may thus reside in a more permissive transcriptional environment, while those in the core face repressive conditions. Cells with more high-*D*_PD_ domains, indicating greater transcriptional surface area, should therefore exhibit enhanced chromatin accessibility and a more efficient stress response. We quantify this using an average nuclear scaling parameter, *D*_*n*_, where *D*_*n*_≈ ⟨*D*_PD_⟩· VF. Here, ⟨*D*_PD_⟩ is the mean scaling exponent of all packing domains within a nucleus, and VF is the fraction of nuclear volume occupied by domains. As illustrated in Fig. 1A, cells with lower *D*_*n*_ are expected to have fewer high-*D*_PD_ domains and reduced transcriptional surface area, while cells with higher *D*_*n*_ access more gene surfaces, enhancing their adaptive response to stress.

Given that packing domains modulate gene expression by influencing macromolecular crowding and accessibility of genes to transcriptional reactants (e.g., transcription factors and RNA polymerase II), we examined how these elements jointly shape transcriptional outcomes. Specifically, we focused on the interplay between crowding and reactant availability, as these factors are central to mechanisms underlying non-genetic resistance. Two key modes of transcriptional adaptation include front-loading, characterized by high reactant concentrations that sustain pre-activated gene expression, and transcriptional plasticity, marked by rapid, stress-responsive activation at lower reactant levels. To investigate these mechanisms, we employed our established computational model of transcription which simulates key steps from transcription factor binding to mRNA synthesis using differential equations (21, 22). We systematically varied crowding (0 ≤ *ϕ* _in_ ≤ 0.55) and reactant concentrations (1× 10^−7^ to 1× 10^−2^ mM) to model gene expression patterns in both front-loading (high reactant) and plasticity (low reactant) conditions (see Table S1 and Supplementary Material). Our findings reveal that genes at low reactant concentrations (0.5 nM) are notably sensitive to crowding effects, while genes with higher concentrations (2 nM) show stable expression as crowding increases up to *ϕ*_in_ ≈ 0.3 (Fig. 1C). This buffering effect suggests that elevated reactant levels in front-loaded genes provide stability against crowding, whereas plastic genes exhibit greater responsiveness to chromatin structural changes. These results align with our previous studies showing that front-loaded, highly expressed genes exhibit minimal differential expression post-chemotherapy, while low-expression, plastic genes demonstrate marked transcriptional shifts (25). Overall, these insights highlight the pivotal role of chromatin structure in modulating transcriptional plasticity, which we focus on in this study.

To address the unknown spatial distribution and exact identity of stress-response genes critical to survival, our model generalizes gene expression patterns, enabling broader predictions of transcriptional responses. Using a probabilistic approach, we characterize chromatin packing across nuclei and predict global gene expression shifts. The probability distribution functions (PDFs) of these shifts are shown in Fig. 1A. We define the transcript ratio *x* = *N*_2_*/N*_1_ to quantify relative gene expression changes, which are assumed to follow a log-normal distribution:

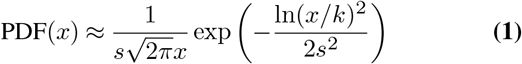

where *k* represents the typical change in gene expression and *s* reflects expression variability. For *s* ≪ 1, the mean expression change is roughly ln(*k*) and *s* can be approximated by the coefficient of variation (COV) of expression changes. Within this framework, *k* (termed transcriptional malleability) quantifies the capacity of a cell to adjust genetic programs in response to chemotherapy, while *s* (transcriptional heterogeneity) captures the range of expression responses within the cell population. Temporal changes in gene expression are modeled by relating transcript number *N* and expression rate *E* by 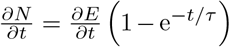, where *τ* is the mRNA elimination time constant. For stress-response dynamics, we approximate the gradual adaptation of gene expression through a first-order exponential growth model:

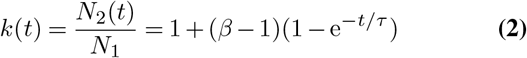

where *β* = *E*_2_*/E*_1_ denotes the post-chemotherapy change in expression rate. We further model changes in gene expression variability by incorporating initial variability and stress-induced adjustments over time:

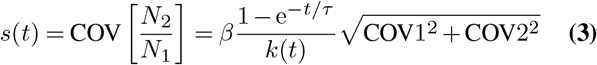

where COV is dependent on *ϕ*_in_ and *D*_*n*_ (see Supplementary Material).

Using the CPMC model, we explored how specific packing domain features shape transcriptional patterns, allowing us to predict both the mean (*k*) and variability (*s*) in gene expression changes (*x*). Specifically, we assessed the sensitivity of gene expression, denoted as *Se*(*E*), to various transcriptional regulators. Genes with similar molecular features (i.e., transcriptional reactant concentrations) were grouped to simplify the non-monotonic relationship between crowding and mRNA output (Fig. 1C), which we represented as a function linking transcriptional output to molecular inputs (Fig. S1 and Supplementary Material). We then examined three structural factors influencing gene expression within packing domains: the average nuclear scaling parameter *D*_*n*_, average genomic content per packing domain ⟨ *N*_PD_⟩, and average DNA fiber packing efficiency within domains ⟨*A*_*v*_⟩ (with *A*_*v*_ = 1 indicating a fully fractal domain). Our findings reveal that *D*_*n*_ exerts the strongest influence on gene expression, positioning *Se*(*E, D*_*n*_) = ∂ ln(*E*)/∂ ln(*D*_*n*_) (23, 25) as central our analysis (Fig. S2 and Supplementary Material). Using *Se*(*E, D*_*n*_) and known expression parameters from a reference cell *a*, we estimated the expression rate change *β* for cell *b* through:

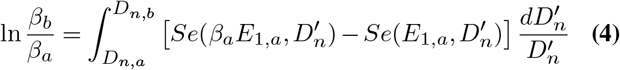

By solving for *β* and incorporating it into Eqs. 2 and 3, we derive the overall transcriptional response to chemotherapy as a function of *D*_*n*_. At higher *D*_*n*_ levels in cell *b*, both *k* and *s* increase (Fig. 1B), indicating that cells with more high-*D*_PD_ packing domains exhibit enhanced transcriptional plasticity and a wider range of gene expression responses.

We next developed an equation that correlates gene upregulation with cell death probability, noting that downregulation follows a similar formalism. The CDA model posits that cell survival relies on reaching critical expression thresholds, denoted as *x*_crit_. The probability of cell death, Θ, is represented by the cumulative distribution function (CDF) of *x*_crit_, yielding a sigmoidal form that can be approximated by a Hill equation (Supplementary Material). For cells *a* and *b*, characterized by distinct chromatin packing *D*_*n,a*_ and *D*_*n,b*_, the death probability Θ for cell *b* can be defined as:

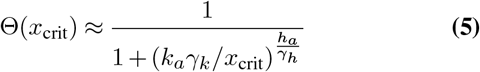

where 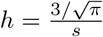 represents the inverse of COV or heterogeneity, *γ*_*k*_ = *k*_*b*_/*k*_*a*_ is the mean upregulation ratio between cells, and *γ*_*h*_ = *h*_*a*_/*h*_*b*_ = *s*_*b*_/*s*_*a*_ denotes differences in transcriptional variability. When *D*_*n,b*_ *> D*_*n,a*_, the CDA model predicts greater malleability and heterogeneity in cell *b*, shifting the log-normal distribution of gene upregulation and increasing survival probability (Figs. 1A-B). Among the parameters in Eq. 5, the relationship between *D*_*n*_ and cell death probability is primarily driven by expression change *β*_*a*_ and the critical threshold *x*_crit_. An elevated *x*_crit_ reduces survival prospects for low *D*_*n*_ cells (Fig. 1D), whereas increased upregulation (*β*_*a*_) enables high *D*_*n*_ cells to minimize death probability (Fig. 1E). Other factors, such as the initial expression rate before chemotherapy (ln(*E*_1_/*E*1), influenced by transcriptional regulators) and time post-exposure, have less impact on the *D*_*n*_–cell death probability relationship compared to *x*crit and *β*_*a*_ (Fig. S3). In summary, the CDA model establishes a physics-based framework linking chromatin structure to cellular adaptability. To validate these predictions, we next conducted experiments on diverse cancer cell lines, examining how temporal changes in *D*_*n*_ influence survival rates under chemotherapeutic stress.

### Pre-treatment organization of packing domains influences cancer cell survival across cell types and cytotoxic mechanisms

To thoroughly test the CDA model prediction that pre-stress nuclear chromatin structure impacts cell death, we employed a live-cell imaging modality that allowed us to visualize chromatin dynamics over time, linking initial chromatin states directly to subsequent survival outcomes. We utilized PWS microscopy, a label-free optical technique that captures wavelength-dependent variations in backscattered interference spectra (23). These variations reflect nanoscale differences in mass density, allowing for insights into chromatin structure (30). PWS microscopy is sensitive to length scales ranging from approximately 20 to 300 nm (31), enabling the detection of chromatin features from the nucleosome level (around 10 nm) to higher-order packing domains (approximately 160 nm in diameter). The spectral variations obtained are associated with the spatial autocorrelation function of chromatin mass density, with its shape corresponding to the nuclear chromatin organization parameter *D*_*n*_ (32). This capability allows for real-time monitoring of population-level *D*_*n*_ through time-course imaging, generating pixel-wise maps of packing domain organization, where each pixel value *D*_pixel_ represents the average across all packing domains within that pixel. To assess the impact of drug treatments on packing domains, we averaged *D*_pixel_ across individual nuclei to determine *D*_*n*_ and analyzed the resulting population-wide distribution of *D*_*n*_. Additionally, since PWS imaging is conducted at the cell-glass interface, only adherent, viable cells were imaged, as apoptotic cells tend to detach rapidly from the surface (33). Consequently, the observed changes in *D*_*n*_ population distributions reflect trends among cells that survive cytotoxic treatments.

Changes in chromatin organization within cancer cell populations during chemotherapy treatment were assessed by imaging at biologically relevant time points. Previous studies have shown that a two-hour pulse treatment with platinum drugs can induce chemoresistance within two days (34–36). Moreover, oxaliplatin can initiate the apoptotic cascade in HCT116 colon cancer cells within six hours, resulting in a significant increase in membrane-permeable dead cells after 24 hours (37). Accordingly, we imaged cell populations at six time points: immediately before chemotherapy administration (0 hours) and at 2, 6, 12, 24, and 48 hours post-treatment.

The CDA model predicts that cells with elevated *D*_*n*_ have an increased likelihood of surviving chemotherapy. To test this hypothesis, we examined changes in *D*_*n*_ at the population level, expecting that over time, only cells with higher *D*_*n*_ would persist. Following chemotherapy treatment, we observed a time-dependent increase in the average *D*_*n*_ of the population which became statistically significant at 24 hours post-treatment (*P <* 0.001; Fig. 2A). This shift in *D*_*n*_ coincided with the emergence of cell death markers and decreased cell viability (Fig. S4). Specifically, the population displayed a trend toward higher *D*_*n*_ starting at 12 hours, with average increases reaching approximately 4.5% after 24 hours (*P <* 0.001) and 10% after 48 hours (*P <* 0.001). In contrast, untreated control populations showed no significant changes in *D*_*n*_ across the same time frame. While these population-level observations support model predictions, they do not reveal whether the initial chromatin state influences individual treatment outcomes. To investigate this, we tracked small clusters of 2-5 HCT116 cells before and after 48 hours of ox-aliplatin treatment Analysis at this cluster level demonstrated a strong negative correlation (*R*^2^ = 0.75) between baseline chromatin state (*D*_*n*,0_, measured at 0 hours) and the percentage increase in *D*_*n*_ after chemotherapy (Fig. 2B). Clusters with lower initial *D*_*n*_ experienced a more substantial increase, whereas clusters with higher initial *D*_*n*_ showed only minor increases. Furthermore, clusters with initially higher *D*_*n*_ demonstrated greater resilience, with 9 clusters having *D*_*n*_ *>* 2.1 surviving treatment, compared to only 3 clusters with *D*_*n*_ *<* 2.1. These findings underscore the heterogeneity in response to chemotherapy and highlight that initial chromatin state influences survival outcomes.

**Fig. 2.**
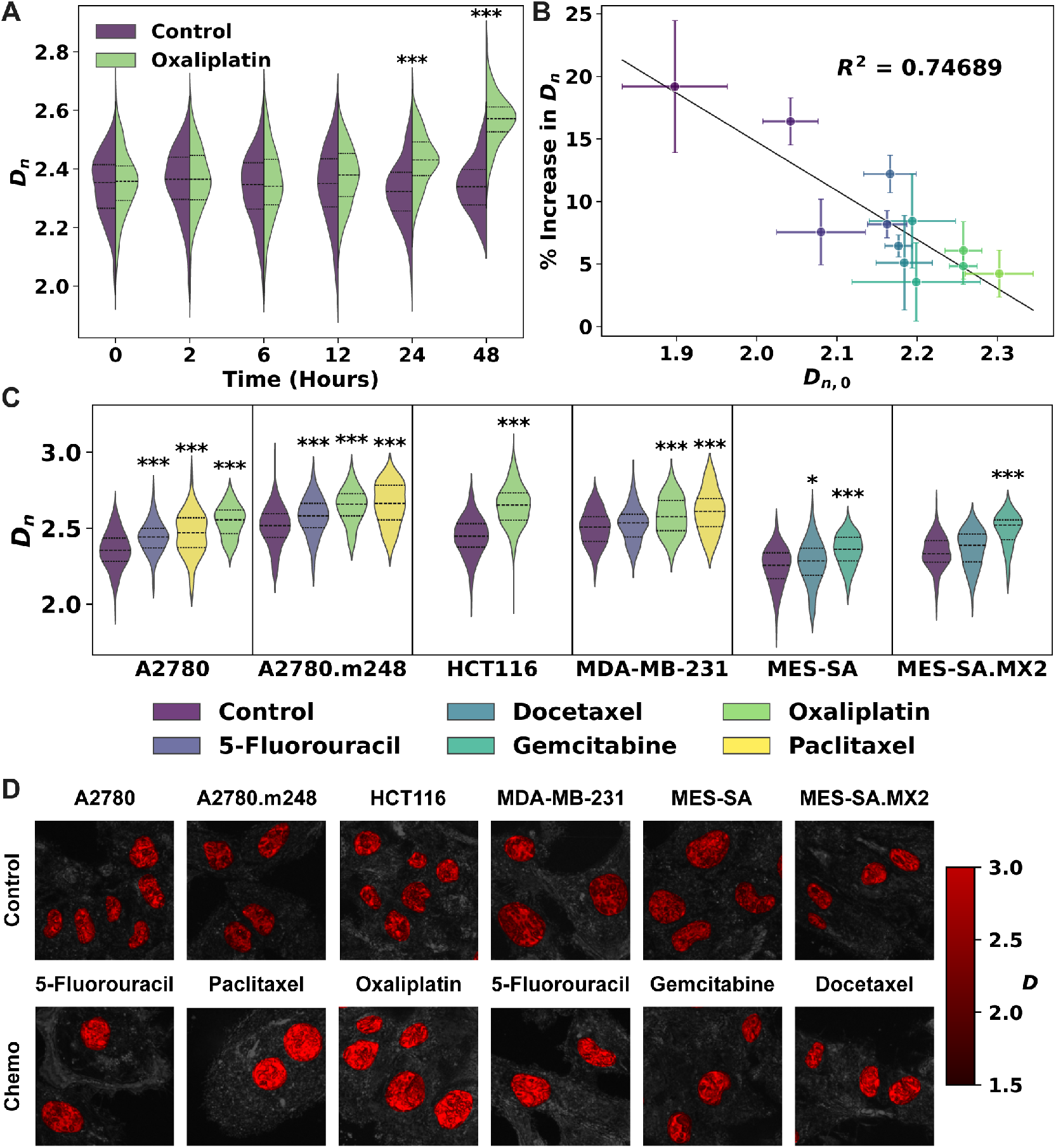
Chemotherapy induces alterations in chromatin across diverse cancer cell lines and treatment modalities. (A) Violin plots showing the distribution of *D*_*n*_ in HCT116 cells over a 48-hour treatment with 15 µM oxaliplatin. The control population remains stable, while treated cells show significantly higher *D*_*n*_ at 24 hours (*P* < 10^−15^) and 48 hours (*P* < 10^−32^). Sample sizes: *n* = 70 − 150 cells per condition. (B) Scatter plot of changes in *D*_*n*_ in individual HCT116 cell clusters after 48 hours of oxaliplatin treatment. Points represent average *D*_*n*_ change per cluster vs. initial *D*_*n*_ at 0 hours (*D*_*n*,0_). Initial cluster size ranged from 2 to 5 cells, while final size ranged from 1 to 12 cells. Error bars represent standard error of the mean. (C) Violin plots showing *D*_*n*_ distribution in surviving cells after 48-hour chemotherapy exposure across multiple cancer cell lines. Treatments include A2780, A2780.m248, MDA-MB-231 (vehicle, 5-fluorouracil, paclitaxel, oxaliplatin); HCT116 (vehicle, oxaliplatin); MES-SA, MES-SA/MX2 (vehicle, docetaxel, gemcitabine). Significance levels: ^∗^*P*<0.05, ^∗∗^*P*<0.01, ^∗∗∗^*P*<0.001 (t-test against control, unpaired, unequal variance). (D) Representative PWS microscopy images of control and treated cells after 48-hour treatments: A2780 (0.5 µM 5-fluorouracil), A2780.m248 (5 nM paclitaxel), HCT116 (15 µM oxaliplatin), MDA-MB-231 (0.5 µM 5-fluorouracil), MES-SA (50 nM gemcitabine), and MES-SA/MX2 (5 nM docetaxel). Pseudocolor indicates that brighter red corresponds to higher *D*_pixel_. Scale bars represent 15 µm.

Following the characterization of chromatin in HCT116 cells, we investigated whether the chemotherapy-dependent increase in population-level *D*_*n*_ is a widespread phenomenon across various cancer types and drug classes. Our study encompassed six cancer cell lines treated with three classes of chemotherapy: DNA intercalators (oxaliplatin), microtubule assembly inhibitors (paclitaxel and docetaxel), and nucleotide analogs (5-fluorouracil and gemcitabine). Cells were exposed to doses of these chemotherapies commonly employed as standard care for specific malignancies (Table S3). Specifically, ovarian cancer cells (A2780 and A2780.m248) and breast cancer cells (MDA-MB-231) received treatments with paclitaxel, oxaliplatin, or 5-fluorouracil (38–40), while colon cancer cells (HCT116) were treated with oxaliplatin, and leiomyosarcoma cells (MES-SA and MES-SA/MX2) were administered docetaxel or gemcitabine (41–43). The *D*_*n*_ distributions for control and chemotherapy-treated populations reveal consistent increases in *D*_*n*_ across diverse cancer types and drug classes (Fig. 2C). For the majority of drug-cell line combinations, these differences in *D*_*n*_ between control and treated populations were significant (*P* < 0.001; Fig. 2C). Additionally, representative PWS microscopy images in Fig. 2D visually depict the chromatin changes induced by various chemotherapy treatments. Notably, the extent of *D*_*n*_ increase correlates with chemotherapeutic efficacy; for instance, docetaxel exhibited lower efficacy compared to gemcitabine in soft tissue sarcoma (44). This relationship is further supported by substantial increases in *D*_*n*_ (*P* < 0.001) for more effective treatments like gemcitabine, contrasting with smaller or non-significant changes observed with less effective agents such as docetaxel (Fig. 2C). Similarly, 5-fluorouracil, which has shown limited efficacy as a single agent (45) yet is commonly used in adjuvant therapy (38, 40), induced the smallest change in *D*_*n*_ among ovarian and breast cancer cells.

Given the observed correlation between the increase in *D*_*n*_ and the efficacy of chemotherapy, we hypothesized that *D*_*n*_ may remain elevated in cells that develop stable resistance to chemotherapy. To test this hypothesis, we utilized two models of drug resistance: one involving a point mutation in a tumor suppressor gene and another with inherent drug resistance. For the first model, we examined A2780 cells harboring point mutations in the TP53 gene, a critical tumor suppressor, which demonstrated higher *D*_*n*_ levels in two subclones compared to the wild-type (Figs. S5A-B). Furthermore, we analyzed *D*_*n*_ data in relation to previously calculated survival outcomes from The Cancer Genome Atlas (TCGA) for patients with high-grade serous ovarian carcinoma exhibiting the same TP53 mutations (46), revealing an association between elevated *D*_*n*_ and low patient survival (Fig. S5C). In the second model, we focused on a drugresistant MES-SA subclone that also displayed higher *D*_*n*_ levels relative to its drug-sensitive counterpart (Figs. S5D-E). Collectively, these findings illustrate a consistent pattern of *D*_*n*_ increase following chemotherapy exposure across various cancer types and treatment strategies, thereby suggesting broad applicability of our CDA model.

### Chromatin dynamics predict chemotherapeutic response in cancer cells

To assess whether the CDA model quantitatively predicts changes in *D*_*n*_, we calculated the probability of cell death using our HCT116 cluster data from Fig. 2B. Unlike population-level data, cluster-level data allowed us to identify individual clusters that survived chemotherapy, providing data at 0, 24, and 48 hours on the same cell clusters. Our analysis focused on actively dividing cell clusters, where *D*_*n*_ is highly correlated due to inherited packing domain characteristics between parent and progeny cells (19). This approach excluded slow-cycling and quiescent cells, for which the impact of packing domains remains unexplored. To improve calculation accuracy, we corrected for cell division-induced drift within clusters (Supplementary Material). We then used the PDFs of *D*_*n*_ obtained from Fig. 2B to calculate experimental cluster death probabilities. By verifying that the *D*_*n*_ distribution shifted towards higher values across all clusters combined (Fig. 3A), we established a baseline *D*_*n*_ distribution at 0 hours that enabled us to predict the PDFs at 24 and 48 hours post-treatment. Starting with an initial estimate of Θ(*D*_*n*_), we refined the prediction by minimizing the mean squared error (MSE) between predicted and observed PDF(*D*_*n*_) values (see Supplementary Material). This process produced experimental values for Θ(*D*_*n*_), shown as points in Fig. 3C.

**Fig. 3.**
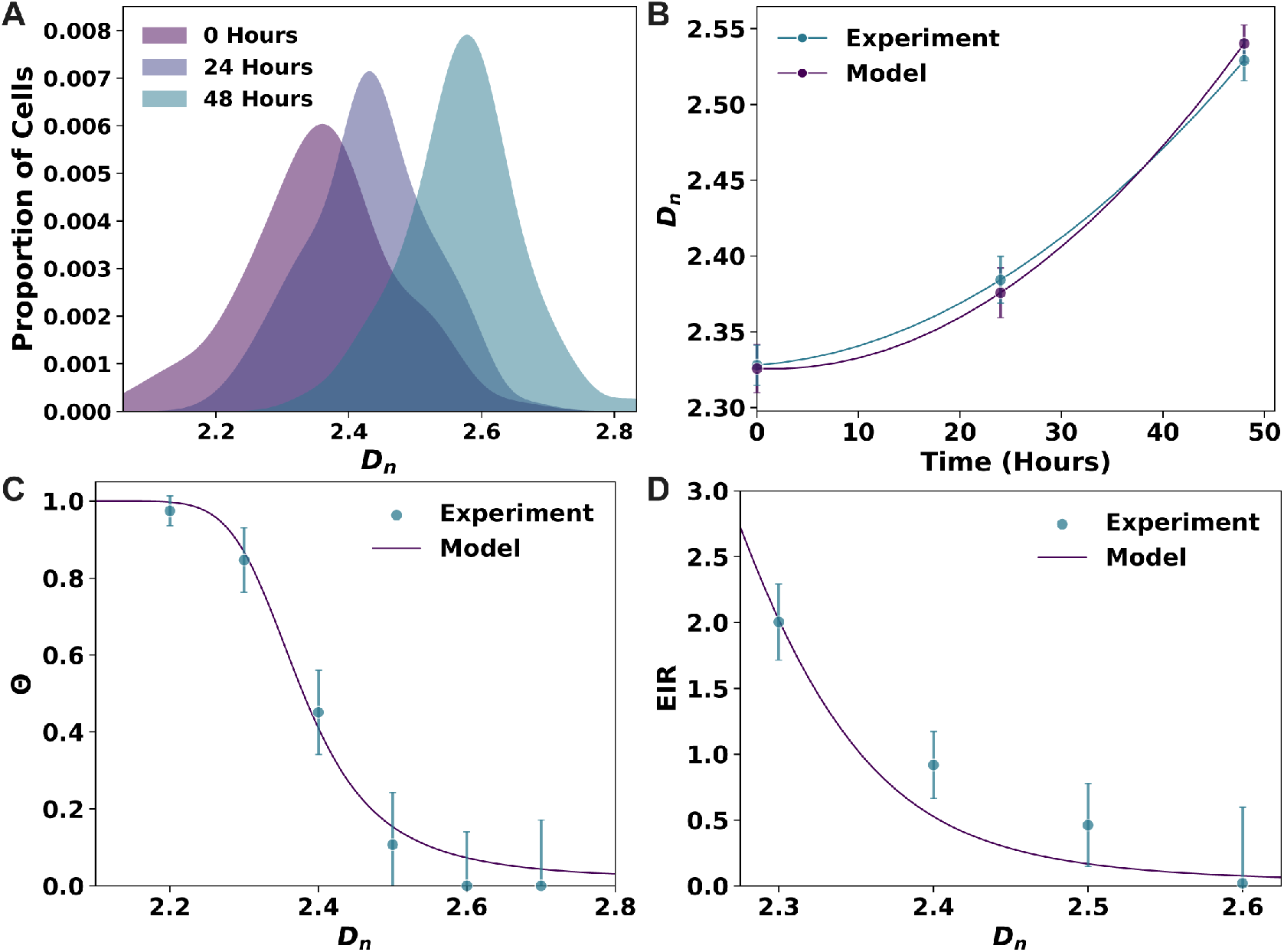
Model predictions of population-level chromatin dynamics, cell survival, and chemotherapy efficacy over time closely match experimental results. (A) Probability density functions (PDFs) of *D*_*n*_ in a population of HCT116 cell clusters at 0 (purple), 24 (blue), and 48 (teal) hours after treatment with 15 µM oxaliplatin, showing a progressive shift toward higher *D*_*n*_ values over time. (B) Comparison of experimental data (blue) and model predictions (purple) showing the increase in mean *D*_*n*_ of oxaliplatin-treated HCT116 clusters over 48 hours. Error bars represent standard error of the mean for experimental data and propagated error for model predictions. (C) Cell death probability (Θ) as a function of *D*_*n*_, with experimental data (blue points) derived from tracking HCT116 cell clusters over 48 hours of oxaliplatin treatment. The CDA model fit (purple line) to experimental data was optimized using three free parameters, resulting in a mean squared error (MSE) of 0.012. (D) Effective inhibition rate (EIR) per day, representing cumulative cell death from oxaliplatin treatment over 48 hours, as a function of mean cluster *D*_*n*_. Both experimental data (blue points) and model predictions (purple line) are shown, with error bars derived from error propagation.

We sought to identify the key parameters driving the relationship between *D*_*n*_ and Θ by fitting CDA model predictions to experimental data. While *D*_*n*_ could be measured directly, parameters critical to the model, such as *β*_*a*_, *x*_crit_, and ln(*E*/Ē), could not be derived from experiments alone. To address this, we implemented an optimization procedure to minimize the MSE between model predictions and experimentally observed Θ(*D*_*n*_) values. This approach assumes uniform gene expression across the cell population, which represents a limitation of our method. The optimized parameter values revealed that genes relevant for cell survival have an average baseline expression level (ln(*E*/Ē) = 0.05), yet require considerable upregulation (*β*_*a*_ ≈ 18) to meet a high threshold (*x*_crit_ ≈ 8). Despite the simplicity of the CDA model and its limited parameter set, it achieved a close fit to the experimental data (MSE=0.012; Fig. 3C). Notably, the MSE was primarily affected by the ratio *β*_*a*_/*x*_crit_, rather than by molecular factors such as the concentration of transcriptional reactants (Fig. S6). This result suggests that for HCT116 cells to survive chemotherapy, critical survival genes must undergo substantial upregulation to reach the activation threshold essential for cell viability. Additionally, the limited sensitivity to baseline expression levels implies that survival may depend less on initial expression and more on the ability of genes to achieve significant upregulation.

Since the CDA model accurately predicted cell death probabilities, we next evaluated whether it could also predict the experimentally observed changes in *D*_*n*_ over time as cells undergo chemotherapy. We hypothesized that the average *D*_*n*_ depends on the cumulative survival probability of individual cells across successive rounds of division. Letting *n*_*τ*_ represent the number of cell division intervals since treatment exposure, 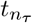 is defined as *n*_*τ*_ *τ*_2_, where *τ*_2_ is the cell doubling time, and cells either divide or die at each *τ*_2_ interval. Here, we found *τ*_2_ ≈ 18 hours, aligning with the average cell cycle duration for HCT116 cells. The survival probability of cells is the opposite of their probability of cell death, Θ, and is given by 1 ™Θ. For a given 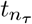,we calculated the population-level average *D*_*n*_ with the following equation, weighting each *D*_*n*_ by survival probability across multiple divisions:

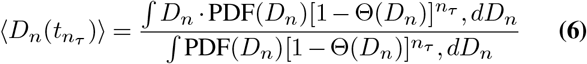

The CDA model predicted that chemotherapy would induce an overall increase in *D*_*n*_ at the population level, consistent with experimental observations (Fig. 3B). Furthermore, model predictions closely matched experimentally observed increases in *D*_*n*_, demonstrating the strong predictive accuracy of the model in capturing population-wide shifts in *D*_*n*_ over time.

To investigate how chromatin packing domains modulate the efficacy of chemotherapy, we calculated the effective inhibition rate (EIR) of oxaliplatin on HCT116 cells over the course of treatment. The EIR represents the cumulative effect of chemotherapy on cell death as influenced by the initial *D*_*n*_, with higher values indicating increased treatment efficacy. We defined the EIR by first modeling the survival probability of cells at time 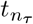,as follows:

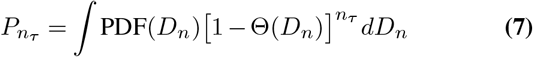

Using this expression, we related EIR to cell survival probability through 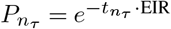,yielding:

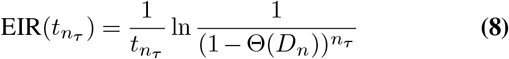

The EIR thus functions as a decay rate constant, quantifying the exponential reduction in cell survival as treatment progresses. Our results, corroborated by both experimental data and model predictions, showed an exponential decrease in EIR with increasing *D*_*n*_ (Fig. 3D). This trend indicates that even slight elevations in *D*_*n*_ substantially impair chemotherapy efficacy, underscoring the potential of targeting chromatin packing domains to improve treatment outcomes.

### Modulating chromatin structure sensitizes cancer cells to chemotherapy

Building on our finding that cells with high *D*_*n*_ demonstrate enhanced survival under chemotherapy, we explored whether chromatin-modifying drugs could increase chemotherapy-induced death in cancer cells. We introduce the term Transcriptional Plasticity Regulators (TPRs) to describe compounds that can decrease *D*_*n*_ to potentially disrupt cancer cell adaptability. Importantly, to maximize chemotherapeutic efficacy while limiting side effects, an essential criterion for TPRs is their selectivity in reducing *D*_*n*_ specifically in cancer cells, sparing non-cancerous cells.

According to the CDA model, isogenic cell populations with higher values of *D*_*n*_ exhibit greater survival under identical cytotoxic stress compared to populations with lower *D*_*n*_ valuess (Fig. 4A). Consequently, the survival of the cell population is strongly associated with the fraction of high *D*_*n*_ cells. To identify effective TPR compounds, we conducted a screen for those capable of rapidly reducing *D*_*n*_, specifically within one hour. Compounds that act within this short time frame are likely to impact nuclear structure directly via physical mechanisms, rather than indirectly through changes in levels of chromatin-modifying proteins. Additionally, this one-hour treatment window was selected to limit the cancer cell adaptability during the initial cellular response to chemotherapy. We identified compounds that might influence packing domains by any mechanism, enabling a broad test of CDA model predictions. For further evaluation, we used the CDA model to identify a threshold of *D*_*n*_ reduction that would maximize probability of cell death (to nearly 100%, with Θ ≈ 0.999) in HCT116 cells, as detailed in the Supplementary Material. This analysis established a critical threshold of *D*_*n*,crit_ = 2.12, which was then applied to identify promising TPR candidates.

**Fig. 4.**
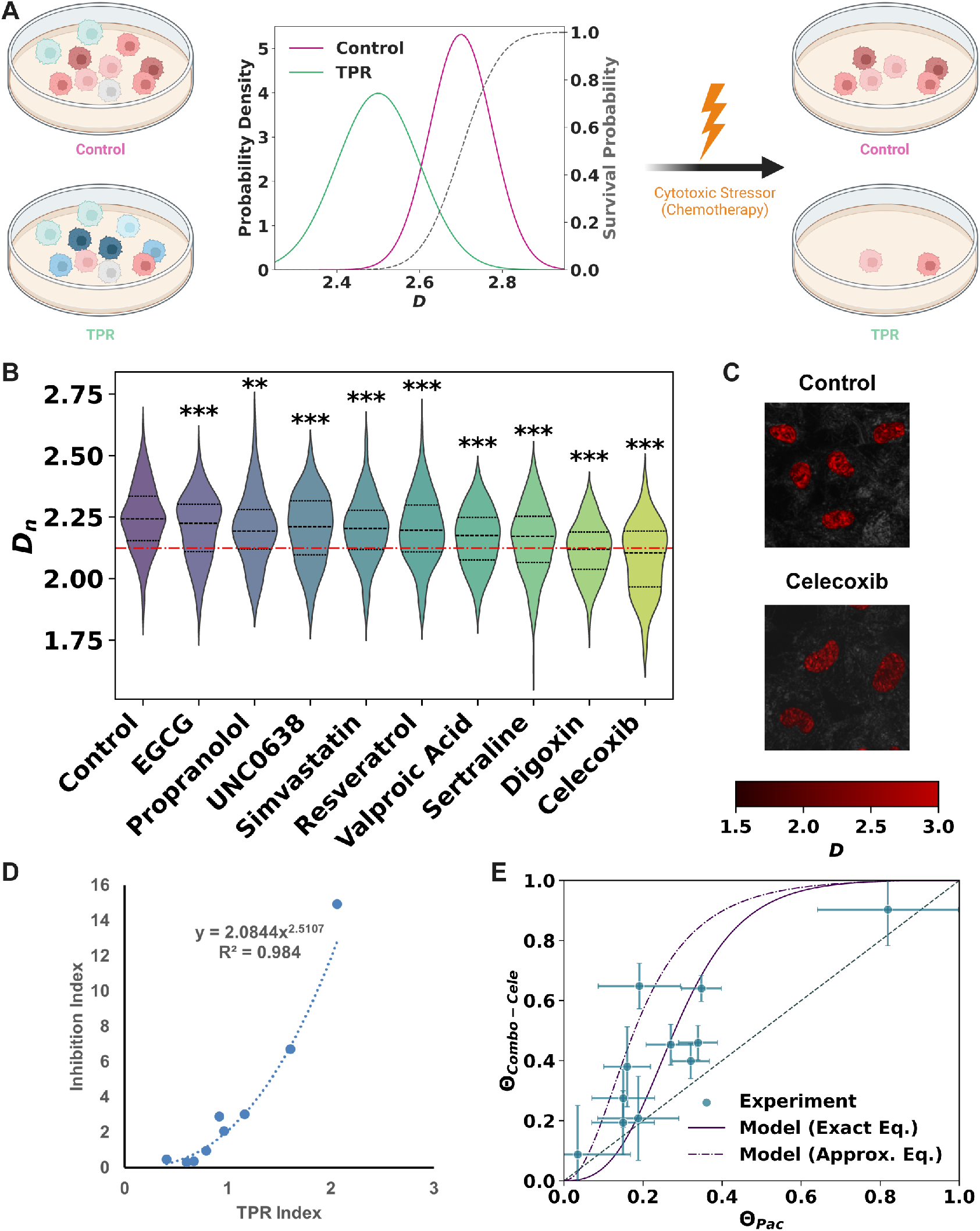
Transcriptional Plasticity Regulators (TPRs) modulate chromatin structure to enhance chemotherapeutic efficacy *in vitro*. (A) Schematic depicting differential survival in cancer cell populations with distinct *D*_*n*_ distributions under chemotherapy. Magenta and green curves represent control and TPR-treated populations, respectively. Dashed grey line indicates survival probability (1™ Θ). (B) TPR candidate drug screen in live A2780 cells, showing *D*_*n*_ distributions for control and potential TPR treatments. Dashed red line denotes *D*_*n*_ threshold below which Θ *>* 0.999. Significance levels: ^∗∗^*P*<0.01, ^∗∗∗^*P*<0.001 (t-test against control, unpaired, unequal variance). (C) Representative PWS microscopy images of control and celecoxib-treated A2780 cells. Pseudocolor indicates *D*_*n*_, with brighter red corresponding to higher values. Scale bars: 15 µm. (D) Correlation between TPR-induced chromatin change (TPR index) and increased cell death (inhibition index) upon combined chemotherapy and TPR treatment, showing that a reduction in *D*_*n*_ correlates with enhanced efficacy. The relationship follows *y* = 2.0844e^2.0107*x*^, with *R*^2^ = 0.984. (E) Comparison of cell death in A2780 cells treated with paclitaxel alone (Θ_Pac_) versus paclitaxel combined with celecoxib normalized by celecoxib alone (Θ_Combo-Cele_). Blue dots represent experimental data with error bars as standard error of the mean. Solid purple line indicates model prediction using exact equation (Eq. 11) with the parameters ln(*E/Ē*) ≈ −3, *β*_*a*_ ≈ 6, and mean squared error (MSE) = 0.031. Dashed purple line is the prediction using approximate equation (Eq. 12).

We screened two main categories of TPRs in live A2780 cells – epigenetic regulators and compounds that could alter the nuclear physicochemical environment – based on their potential to modulate packing domains. A list of these TPR candidates, along with their observed effects on chromatin structure, is presented in Table S4. Epigenetic regulators targeting histone tail modifications, such as acetylation and methylation, were expected to disrupt nucleosome interactions. While these compounds significantly decreased the average *D*_*n*_ values in cancer cells (Fig. 4, *P* < 0.001 for all compounds except propranolol), their overall effect on packing domains remained modest, and Dn did not fall below the critical threshold (*D*_*n*,crit_) associated with near-complete cell death.

Since chromatin is a negatively charged polymer, we hypothesized that altering the ionic environment might enhance chromatin interactions and thereby more widely affect *D*_*n*_. To test this, we investigated the effects of celecoxib, a nonsteroidal anti-inflammatory drug (NSAID) known to influence Na^+^, K^+^, and Ca^2+^ channels (47), and digoxin, an inhibitor of Na^+^/K^+^-ATPase. Both drugs significantly reduced *D*_*n*_ below the critical threshold, *D*_*n*,crit_ (*P* < 0.001; Figs. 4B-C). Furthermore, when combined with chemotherapy, celecoxib and digoxin markedly increased cancer cell apoptosis (*P* < 0.001); however, without chemotherapy, their effects were minimal (*P* > 0.05; Fig. S7), suggesting a synergistic interaction with chemotherapy. Finally, to evaluate cancer-cell specificity, we tested TPR candidates in noncancerous osteoblasts derived from hMSCs. Among these candidates, celecoxib was the only TPR that did not significantly affect *D*_*n*_ in non-cancerous cells (*P* > 0.05; Fig. S8). This cancer-selective modulation of chromatin, coupled with enhanced chemotherapy effectiveness, establishes celecoxib as our lead TPR candidate.

Next, we evaluated the effects of TPRs across multiple cancer types. From our initial screen, we selected compounds that significantly decreased *D*_*n*_ with varying potency, likely through distinct nuclear mechanisms, for further analysis. A 30-minute treatment with each compound revealed slight variations across cell lines (Fig. S9). For example, VPA had stronger effects in A2780.m248 cells than in A2780 cells, while digoxin—effective in A2780, A2780.m248, and MDA-MB-231 cells—was less potent in MES-SA and MES-SA/MX2 cells compared to celecoxib. Importantly, the effects of celecoxib on *D*_*n*_ are not due to COX-2 inhibition, as it was able to reduce *D*_*n*_ in HCT116 cells which are COX-2 deficient (48). This indicates that the effects of celecoxib may involve an alternate ion-modulating pathway. These results are further supported by the smaller impact of aspirin on *D*_*n*_ (Fig. S9), which is a COX-1/2 inhibitor but is not expected to modulate ion channels. Although the exact pathways remain outside the focus of this study, these findings confirm the impact of TPRs on packing domain structure, enabling tests of CDA model predictions.

We next assessed whether top TPRs increased cancer cell death when combined with chemotherapy, measuring percent inhibition as an indicator (0% indicating no effect and 100% indicating complete cell death). Across cancer types, TPRs with chemotherapy increased cell death, with strong TPRs (celecoxib and digoxin) showing the most significant effects (*P* < 0.01; Fig. S10). To quantify the relationship between chromatin modulation and chemotherapy efficacy, we developed two CDA model-based indices to measure the impact of *D*_*n*_ changes on cell death from chemotherapy (see Supplementary Material). The TPR Index assesses the relative impact of one TPR on chromatin packing domains compared to another TPR:

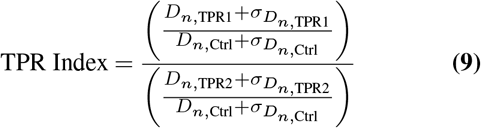

where 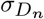 is the standard deviation of the PDF(*D*_*n*_), and measurements of two TPRs and a control required. The inhibition index quantifies added chemotherapy efficacy:

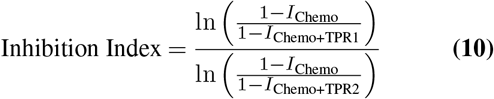

where *I* is percent inhibition (Fig. S10), comparing combination treatment of chemotherapy with one of two TPRs against chemotherapy alone. We observed an exponential relationship between the TPR and inhibition indices (*R*^2^ = 0.984), indicating that larger *D*_*n*_ reductions are associated with greater chemotherapy-induced cell death (Fig. 4D). This synergistic effect suggests that TPRs enhance chemotherapy efficacy beyond individual impacts.

To further examine the effects of TPRs in enhancing chemotherapy-induced cell death, we conducted a quantitative analysis of the CDA model prediction that reducing the fraction of high *D*_*n*_ cells in a population should increase susceptibility to cytotoxic treatments. For this purpose, we used celecoxib as the TPR in combination with paclitaxel and treated A2780 ovarian cancer cells under increasing chemotherapy doses to vary levels of cytotoxic stress. This experimental setup enabled us to investigate whether increased transcriptional plasticity aids cell survival under high-stress conditions. To quantify cell death, we calculated the inhibition rate (IR) based on the fraction of dead cells (percent inhibition, *I*), allowing us to quantify cell death probability as Θ = 1 − exp(− IR ·*t*) (data points in Fig. 4E; detailed methods in Supplementary Material). To generate CDA model predictions, we adapted Eq. 5 to determine the death probability of TPR co-treated cells (Θ_*b*_) using the probability of death in cells treated with chemotherapy alone (Θ_*a*_):

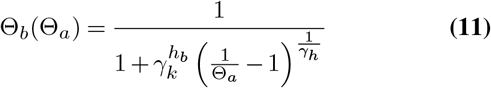

By varying Θ_*a*_ from 0 to 1 for cell population *a* (high *D*_*n*_, chemotherapy alone), we predicted the death probability for population *b* (low *D*_*n*_, chemotherapy combined with TPR treatment). Using the *D*_*n*_ distributions for untreated and celecoxib-treated A2780 cells (Fig. 4B), we optimized model parameters for best fit to experimental data, yielding an initial level of survival gene expression ln(*E*/Ē) ≈ − 3 and an upregulation *β*_*a*_ ≈ 6 (MSE = 0.031). Our analysis showed that gene upregulation *β*_*a*_ was the primary contributor to the model fit, while the initial expression level ln(*E/E*) had a minimal effect (Fig. S11). In agreement with model predictions, celecoxib-treated cells (lower *D*_*n*_) exhibited higher rates of chemotherapy-induced cell death compared to untreated cells (higher *D*_*n*_; Fig. 4E). The combination treatment resulted in a sigmoidal dose-response curve, highlighting the enhanced efficacy of TPR co-treatment over chemotherapy alone.

To identify which model parameters best predict cell death probability, we applied several simplifying assumptions. Specifically, we used steady-state conditions (*t* ≫ *τ*, where *τ* is the mRNA decay rate constant), high gene upregulation (*β*_*a*_ ≫1), a weak dependence of 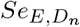 on *D*_*n*_, and small changes in *D*_*n*_ (Δ*D*_*n*_ = *D*_*n,b*_ *™ D*_*n,a*_ ≪*D*_*n,a*_). Under these assumptions, we derived an approximate expression for 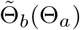

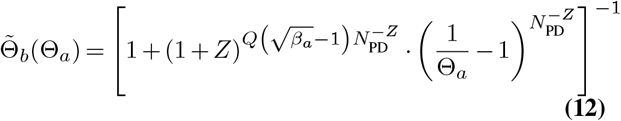

where 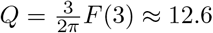 and *Z* = Δ*D*_*n*_/*D*_*n,a*_ (Supplementary Material). This simplified model aligns well with experimental data (dashed line; Fig. 4E), indicating that cell death probability is largely governed by *D*_*n,a*_, Δ*D*_*n*_, and *β*_*a*_. These findings support the CDA model predictions that TPRs synergize with chemotherapy by modulating *D*_*n*_, suggesting potential applications in optimizing therapeutic strategies.

### TPRs improve chemotherapeutic efficacy and mitigate tumor adaptation *in vivo*

To evaluate the potential of TPRs to hinder cancer cell adaptation to chemotherapy beyond our initial *in vitro* findings, we conducted an *in vivo* assessment. Using a patient-derived xenograft (PDX) model of ovarian cancer, we focused on celecoxib, the most potent TPR identified, to examine its effects on tumor adaptation. We hypothesized that mice treated with either vehicle or TPR alone would exhibit increased tumor growth over time (Fig. 5A, red), consistent with our previous observations of limited anti-cancer activity for TPRs alone (Fig. S7). Additionally, our CDA model and *in vitro* data suggested that a combination of chemotherapy and TPR would result in significantly reduced tumor growth compared to chemotherapy alone (Fig. 5A, purple). To minimize chemotherapeutic toxicity, we used a low dose of paclitaxel to ensure survival of mice while observing the effects of chemotherapy. We therefore expected continued tumor growth in the chemotherapy-treated mice. We administered celecoxib (25 mg/kg) in conjunction with a reduced dose of paclitaxel (1.7 mg/kg) to evaluate potential synergy in mitigating tumor growth. Results indicated that tumors in the three single-treatment groups (vehicle, celecoxib, or paclitaxel alone) continued to grow, with the vehicle group exhibiting the fastest growth, followed by paclitaxel and celecoxib (Fig. 5B). Tumors treated with celecoxib displayed slower growth than those treated with paclitaxel, as illustrated by the orange and yellow lines in Fig. 5B, likely due to the induction of cell cycle arrest. Over the course of 30 days, vehicle and paclitaxel-treated tumors doubled in volume. In stark contrast, the celecoxib-paclitaxel combination limited tumor growth to only a 20% increase in volume.

**Fig. 5.**
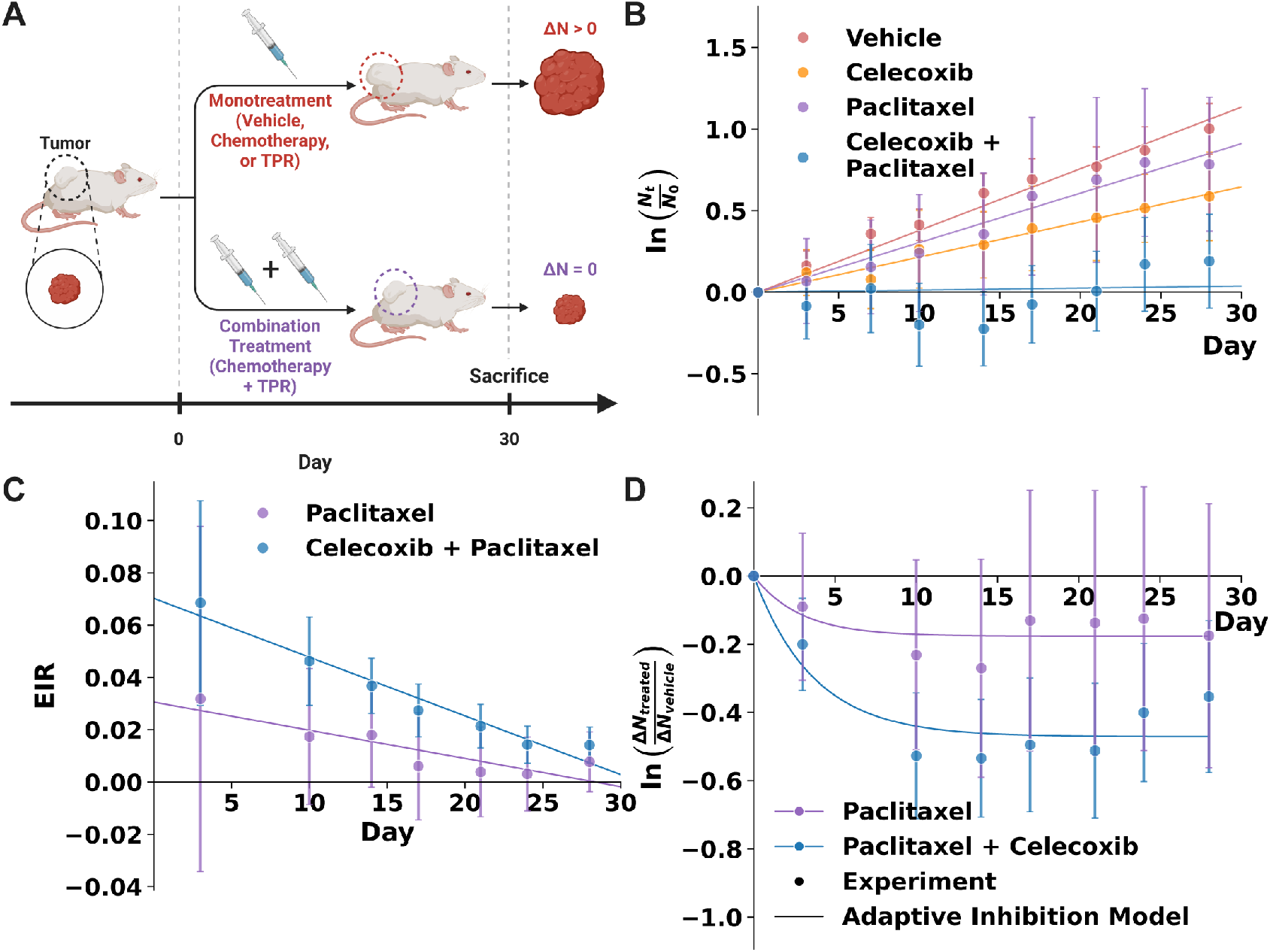
*In vivo* validation of TPR-enhanced chemotherapy efficacy using patient-derived xenograft (PDX) models of ovarian cancer. (A) Schematic illustrating the treatment regimen for ovarian PDX studies and predicted outcomes, highlighting expected tumor growth patterns under different treatment conditions over 30 days. (B) Growth curves of ovarian carcinoma PDX tumors under various treatments. Co-treatment with celecoxib (25 mg/kg) and paclitaxel (1.7 mg/kg) resulted in minimal growth over 30 days compared to monotherapy with paclitaxel (1.7 mg/kg), celecoxib (25 mg/kg), or vehicle (DMSO). Animals were treated orally with celecoxib and intraperitoneally with paclitaxel daily for one week. Points represent mean tumor volume normalized by the volume at day 0 ± standard error of the mean, while lines indicate linear regression fits. Effective Inhibition Rate (EIR) for paclitaxel alone and the paclitaxel + celecoxib combination over time, calculated using vehicle as a reference for paclitaxel and celecoxib as a reference for the combination. EIR was calculated using the equation EIR = [ln(*V*_*t*,control_ /*V*_0,control_) − ln(*V*_*t*,treated_ /*V*_0,treated_)]/*t*. Points show mean EIR ± standard error of the mean, with lines representing linear regression fits. (D) Normalized tumor growth rate over time for paclitaxel alone and the paclitaxel + celecoxib combination. Points indicate experimental data (mean ± standard error of the mean). Solid lines show best fits using the adaptive inhibition model (Eq. 13).

We further evaluated the impact of TPR compounds on chemotherapy efficacy by calculating the effective inhibition rate (EIR) to quantify cumulative cancer cell death at specific time points. By utilizing *in vitro* PWS data from A2780 ovarian cancer cells and extrapolating these results to relevant time scales for the PDX study, our CDA model predicted two key outcomes: first, the EIR would decline over time for both chemotherapy and combination treatments; second, the EIR for combination therapy would consistently surpass that of chemotherapy alone (Fig. S12). We validated these predictions by calculating the EIR from the PDX data using the formula EIR = [ln(*V*_*t*,control_/*V*_0,control_) ™ln(*V*_*t*,treated_/*V*_0,treated_)]/*t*. In accordance with our CDA model predictions, both treatment groups exhibited a decreasing EIR over time, indicating reduced efficacy at later stages; however, the combination treatment maintained a higher EIR throughout the experimental duration (Fig. 5C). The results indicate that TPR compounds enhance the overall effectiveness of chemotherapy by prolonging its inhibitory effects.

To explore how chromatin-mediated transcriptional plasticity influences adaptation to chemotherapy, we developed a mechanistic model that directly relates to our *in vivo* data, overcoming the challenge of quantifying the distribution of *D*_*n*_ within tumors. This model distinguishes between two subpopulations: an “unadaptable” group with lower *D*_*n*_ values and “adaptable” cells with higher *D*_*n*_ values. We hypothesize that the rate of tumor adaptation to chemotherapy depends on the initial *D*_*n*_ values, with cells possessing higher *D*_*n*_ values adapting more rapidly. This implies that a subset of cells will adapt within a critical time frame, determined by the distribution of *D*_*n*_ values across the tumor cell population. To model tumor growth in the absence of adaptation, we define the relative growth rate, *V* (*t*), as ln(*V* (*t*)/*V* (0)) = (*p* ‒ *c*)*t*, where *p* is the cell proliferation rate and *c* is the inhibition rate due to chemotherapy. We refine this expression to incorporate an adaptation term (further details in Supplementary Material):

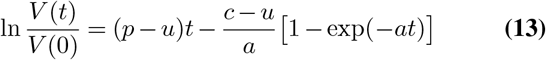

where *t* is the time since treatment initiation, *u* is the unadaptable inhibition rate, and *a* is the adaptation rate.

To validate that our simplified adaptation model aligns with our CDA model, which mechanistically links packing domains to cell fate, we utilized *in vitro* PWS microscopy data from A2780 cells. This approach allowed us to predict tumor volume changes and evaluate the effects of TPR treatment on chemotherapy inhibition rate (*c*) and adaptation rate (*a*). We estimated these rates by fitting the CDA model predictions for relative tumor volume (growth normalized by control; see Supplementary Material) using Equation 13, with *c* and *a* as free parameters. In this analysis, we assumed a constant unadaptable inhibition rate, *u*, across treatments and set *p* = 0, as model predictions were normalized by the control. We initially performed a fit with *u* as a free parameter, yielding an unadaptable inhibition rate of *u* = 0.15, which likely reflects the high chemotherapy dosage applied in the *in vitro* studies (Fig. S13). For chemotherapy treatment alone, we estimated values of *c* = 1.13 and *a* = 2.20 (RMSE = 0.009, *R*^2^ = 0.99).

With the combination of chemotherapy and TPR, we observed a 60% increase in *c* to 1.81 and a 31% decrease in *a* to 1.52 (RMSE = 0.017, *R*^2^ = 0.99). These findings suggest that TPR enhances chemotherapy efficacy over time by reducing *D*_*n*_, which in turn decreases the adaptation rate (Fig. S13).

To evaluate the relevance of our model predictions in an *in vivo* context, we applied the adaptive inhibition model to the PDX data. We fit normalized tumor growth values from both paclitaxel alone (normalized to vehicle control) and the combination treatment with celecoxib (normalized to celecoxib alone), estimating parameters *u, c*, and *a*. We set a fixed proliferation rate of *p* = 1× 10^−10^, given the normalization of tumor volumes against respective controls. Our estimates yielded near zero values for *u*, while the values for *c* were significantly lower than those predicted from *in vitro* data, suggesting that the paclitaxel dose was insufficient to prevent tumor growth and could be readily adapted to. Specifically, for paclitaxel alone, we found *c* = 0.062 and *a* = 0.35 (RMSE = 0.049, *R*^2^ = 0.60), whereas the combination treatment yielded *c* = 0.13 and *a* = 0.27 (RMSE = 0.069, *R*^2^ = 0.85). This indicates a 108% increase in the inhibition rate (*c*) and a 28% decrease in the adaptation rate (*a*) with celecoxib cotreatment compared to paclitaxel alone. In summary, the *in vivo* results corroborate our *in vitro* findings and model predictions, demonstrating that targeted chromatin modulation significantly enhances chemotherapeutic efficacy and limits tumor adaptation in a clinically relevant context.

## Discussion

Treatment resistance remains a major challenge in effective cancer therapy, primarily due to the limited understanding of non-genetic factors (i.e., cellular plasticity) that drive resistance. In this study, we introduce the CDA model as a novel biophysical framework that links chromatin organization to cellular adaptability and chemotherapy resistance. The model incorporates the effects of macromolecular crowding through the chromatin density distribution within packing domains, enabling a direct association between chromatin structure and cellular phenotype through transcriptional regulation. Our simplified model predicts that cell survival under treatment relies on the upregulation of key genes, a process dependent on the chromatin packing within these domains. Importantly, the model demonstrates that the likelihood of chemotherapy-induced cell death can be predicted by the nuclear chromatin scaling parameter, *D*_*n*_, at the onset of treatment (Fig. 1). In multiple cancer types and across different chemotherapy mechanisms, we consistently observed increases in *D*_*n*_ following treatment (Fig. 2), suggesting an extensible adaptive stress response that aids treatment evasion. As elevated *D*_*n*_ appears to facilitate chemotherapy resistance (Fig. 3), we investigated targeting of chromatin packing domains as a therapeutic strategy. Notably, TPR agents that reduced *D*_*n*_ substantially increased chemotherapy efficacy (Fig. 4), with the leading candidate showing enhanced *in vivo* effectiveness in a PDX model of ovarian cancer (Fig. 5). Specifically, our adaptive inhibition model revealed that combining paclitaxel with celecoxib reduced adaptation rates and improved tumor growth inhibition compared to paclitaxel alone.

Our findings align with recent studies underscoring the role of transient transcriptional states in promoting chemoresistance through mechanisms such as altered drug accumulation, enhanced drug export, and changes in drug targets and signaling pathways (49). Cancer cells adapt rapidly—often within timescales shorter than cell division—by leveraging diverse transcriptional networks, emphasizing the need for compounds that can interrupt these responses during therapy (23, 50). Househam et al. observed extensive transcriptional diversity both within and between colorectal tumors, linking cancer plasticity with transcriptional malleability (51). Shaffer et al. further demonstrated that such transcriptional heterogeneity allows cells to endure adverse conditions long enough for mutations to emerge (52). Additionally, Heide et al. found that genetic and non-genetic mechanisms evolve together within tumors, highlighting the necessity of targeting both for effective treatment (53). Beyond cancer, our findings reflect broader adaptability principles, such as the “plasticity-first” evolution theory, which suggests that environmental triggers can initiate phenotypic plasticity, facilitating evolutionary adaptation under stress (54, 55). Here, plasticity involves transcriptional changes that improve fitness in response to stressors and reveal genetic variation, often with minimal impact except under specific, challenging conditions. Such principles have been applied in natural systems, including coral resilience amid environmental shifts (55). In cancer cells, we find that elevated *D*_*n*_ correlates with increased transcriptional malleability and heterogeneity, influencing cell phenotypes similarly to plasticity observed in plants.

While our findings offer valuable insights, several limitations of our model and experimental approach warrant consideration. First, although we use *D*_*n*_ as an indicator of nuclear packing domain organization, this measure provides only an averaged view that potentially oversimplifies the inherently complex three-dimensional arrangement. As a result, we are unable to probe the gene-specific spatial localization within domains that could influence transcriptional activity. To gain finer insights into gene positioning within packing domains, approaches like DNA fluorescence *in situ* hybridization (FISH) or CRISPR-based tagging could label gene loci for co-registration with packing domains identified by PWS microscopy. Second, our cluster tracking data (Fig. 2) reveal that clusters with initially low *D*_*n*_ levels tend to increase *D*_*n*_ over time, in contrast to clusters with higher initial *D*_*n*_. This suggests that cells with low *D*_*n*_ may either undergo cell death or use adaptive strategies, potentially by increasing the number of packing domains to enhance survival. While our CDA model predicts a higher likelihood of cell death in low *D*_*n*_ cells, additional studies are needed to determine if these cells can indeed survive by upregulating packing domains as a form of adaptive response. Third, our model simplifies cellular decision-making to either survival or death at defined checkpoints, though cellular responses are often more nuanced. For instance, some cancer cells evade chemotherapy by exiting the cell cycle (56), while others become quiescent in response to irreversible damage (57, 58). Future model adaptations could incorporate cell cycle arrest pathways, potentially informed by regulatory factors such as p53 dynamics (59), to enhance the capacity to predict diverse cellular outcomes. Finally, we recognize that TPR compounds, like celecoxib, may influence cell cycle progression independently of their effects in combination with chemotherapy (60). Additionally, chemotherapy itself can impact cell cycle dynamics by prolonging mitotic intervals (61). Future iterations of our model and corresponding experimental validations should account for these complexities, providing a more holistic framework for understanding chromatin organization and cellular responses in therapeutic contexts.

In conclusion, this study presents a novel framework linking chromatin packing domains with chemoresistance, unveiling chromatin-regulated transcriptional plasticity as a key mechanism in cancer cell adaptation. By identifying TPR compounds that enhance chemotherapy efficacy, we demonstrate the potential of chromatin modulation as a strategy to counteract chemoresistance, with observed synergies offering promising avenues for high-efficacy combination therapies. Our findings reveal chromatin packing domains as pivotal in modulating cellular responses, highlighting the need for innovative imaging approaches and interventions that target both local and global regulators of chromatin structure. While our screen of TPR candidates was limited to epigenetic remodelers and compounds influencing nuclear ionic balance, future work should investigate additional chromatin regulators (i.e., remodeling complexes and pioneer transcription factors) as well as the underlying mechanisms of how ions influence chromatin conformation. Expanding these studies to a wider range of cancer types and treatments, including immunotherapies, could further elucidate the role of chromatin-mediated adaptation in treatment resistance. Ultimately, our insights into chromatin organization pave the way for new cancer therapies that target adaptive resistance mechanisms, offering a promising strategy for improving patient outcomes.

## Materials and Methods

### Cell Culture and Treatments

Leiomyosarcoma (MES-SA, MES-SA/MX2), breast (MDA-MB-231), colon (HCT-116, HT-29), and mouse embryonic fibroblast (MEF) cell lines were obtained from ATCC, while ovarian lines (A2780 variants, Ovcar8) were provided by Dr. Chia-Peng Huang Yang. All lines were cultured in ATCC-recommended media with 10% FBS and routinely tested for mycoplasma. Human mesenchymal stem cells (hMSCs, ATCC) were cultured in DMEM with 10% FBS and differentiated with osteogenic medium (Lonza) on day 2 post-seeding. Cells were plated on glass-bottom dishes and treated when 30% confluent. Chemotherapy treatments included paclitaxel (5 nM), oxaliplatin (15 µM), 5-fluorouracil (500 nM), docetaxel (5 nM), and gemcitabine (50 nM) for 48–72 hours. TPR compounds, including celecoxib (75 µM), valproic acid (100 µM), aspirin (1 mM), and others, were administered 30 minutes before imaging. Untreated controls were included for each cell type, and cells were imaged under physiological conditions (5% CO_2_, 37°C). For additional details, please refer to the supplementary methods.

### Partial Wave Spectroscopic (PWS) Microscopy

PWS microscopy was conducted on an inverted microscope (Leica DMIRB) equipped with a Hamamatsu CCD camera and a liquid crystal tunable filter, capturing spectrally resolved images from 500 to 700 nm (1 nm intervals). Pixel size at the sample plane was 267 nm, approaching the theoretical resolution limit of 261 nm with a 60x objective and 1.4 NA. Full details of the PWS setup are described in Almassalha et al. (23). Interference spectra were normalized to the incident light reflectance, and a Butterworth filter was applied to reduce noise. The nuclear chromatin organization parameter *D*_*n*_, derived from standard deviation (Σ) calculations of spectra, reflects chromatin density organization and was analyzed through custom MATLAB scripts (32). Pixel-wise *D* maps were generated, and *D*_pixel_ values were averaged per cell nucleus to find *D*_*n*_ across 100-200 cells per condition. Pseudo-colored PWS images were produced in Python, with *D*_*n*_ values (range 2-3) mapped to a red colormap. For additional methodological details, please refer to the supplementary methods.

### Cell Viability and Confluence Measurements

Cell viability assays were performed using fluorescence measurements with a BioTek Synergy Neo2 Reader at the Northwestern University HTAL Core facility. HCT116 cells, seeded at 1,500 cells per well in 96-well plates, were treated with 15 µM oxaliplatin and assessed at 0, 2, 6, 12, 24, and 48 hours using the ApoTox-Glo triplex assay. Fluorescence intensity was measured following a 30-minute incubation with the viability reagent. To calculate inhibition rates (*IR*), cell density was quantified via transmission microscopy and analyzed using ImageJ. Cell confluence data for treated versus control groups were used to calculate *IR* = [1/(*t*_*n*+1™_ *t*_*n*_)] •ln[*C*(*t*_*n*+1_)/*C*(*t*_*n*_)] and normalized inhibition *I*_norm_ = *C*_treatment_/*C*_control_. Further methodological details are provided in the supplementary methods.

### Chromatin-Dependent Adaptability (CDA) Model Implementation and Optimization

The CDA model was implemented in Python using NumPy and SciPy. Parameter scans and grid searches were conducted for key parameters: *D*_*n*,0_, ln(*E/Ē*), *β*_*a*_, *x*_crit_, and *T*_crit_. The `scipy.optimize.minimize` function with L-BFGS-B optimization was used to fit the model to experimental data, minimizing the MSE between model predictions and observed Θ vs. *D*_*n*_ curves. Key parameters optimized were *β*_*a*_, *x*_crit_, and ln(*E/Ē*), achieving an MSE of 0.031 for HCT116 cells with oxaliplatin. The *D*_*n,crit*_ threshold for TPR efficacy was iteratively refined based on control group *D*_*n*_ values, yielding a *D*_*n,crit*_ of approximately 2.123 with Θ = 0.999. Optimization for Θ_*b*_ vs. Θ_*a*_ used A2780 cell confluence data under TPR (celecoxib) and chemotherapy (paclitaxel), yielding *β*_*a*_ ≈ 6 and ln(*E*/Ē) ≈ − 3 with an MSE of 0.031. Complete methodological details are provided in the supplementary methods.

### Patient-Derived Xenograft (PDX) Tumor Models

Patient-derived xenograft (PDX) models were generated from High Grade Serous Ovarian Cancer (HGSOC) tissue obtained from chemotherapy-naive patients at the Northwestern University Prentice Women’s Hospital with IRB approval. Fourth-passage tumor fragments were subcutaneously implanted into NOD/SCID gamma (NSG) mice. Once tumors reached 150–200 mm^3^, mice were randomized into treatment groups: celecoxib control, paclitaxel control, 25 mg/kg celecoxib, 1.7 mg/kg paclitaxel, and a combination of celecoxib and paclitaxel. Celecoxib (25 mg/kg) was administered daily by oral gavage, and paclitaxel (1.7 mg/kg) twice weekly intraperitoneally. Tumor volume (*V* = *l •w*^2^/2) and body weight were monitored biweekly over 4 weeks, after which tumors were harvested post-euthanasia. Nonlinear least squares optimization was used to fit the adaptive inhibition model to the data. Further methodological details are available in the supplementary methods.

### Statistical Analysis

Statistical analyses were performed in Python, using SciPy and pandas for data processing. Welch’s t-test was employed for pairwise comparisons between a reference condition and other conditions across groups, including assessments of nuclear *D*_*n*_ values, cell viability, and PDX tumor volumes. Statistical significance was defined as *P* < 0.05 and denoted as ^∗^*P* < 0.05, ^∗∗^*P* < 0.01, and ^∗∗∗^*P* < 0.001. Violin plots were used for *D*_*n*_ distributions, showing quartiles and frequency distribution, with SEM as the standard for error bars. Additional details are provided in the supplementary methods.

## ACKNOWLEDGEMENTS

This work was supported by National Institutes of Health grants U54CA268084, U54CA193419, R01CA228272, R01CA225002, R01CA155284, R01CA165309, T32GM132604, T32GM008152, and T32HL076139; National Science Foundation grants EFMA-1830961, EFMA-1830968, EFMA-1830969, CBET-1249311, EFRI-1240416, DGE-0824162, and DGE-1842165; the Lefkofsky Innovation Award; and the Chicago Biomedical Consortium with support from the Searle Funds at The Chicago Community Trust. We thank Rob and Kristen Goldman, Mr. David Sachs, and the Christina Carinato Charitable Foundation for philanthropic support. Flow cytometry was performed at the Northwestern University Flow Cytometry Core Facility, supported by NCI Cancer Center Support Grant CA060553.

