## Supporting Information for "Leveraging chromatin packing domains to target chemoevasion *in vivo*"

**Characterization of Nuclear Chromatin Structure.** We examine how chromatin architecture influences gene expression by linking its spatial organization to transcriptional activity. Chromatin structure is characterized using several scaling exponents, each describing distinct aspects of its distribution across spatial scales. The arrangement of chromatin into mass fractal packing domains (Packing domains) (1, 2) forms the basis of our analysis, with scaling exponents that quantify how chromatin mass or density varies with spatial size. Key exponents include: (1) the polymeric scaling exponent (Flory exponent), which relates polymer size to the number of monomers; (2) the mass scaling exponent for individual packing domains, $D_{\text{PD}}$, reflecting how chromatin mass scales with its size within a domain; (3) the ensemble average of mass scaling across all packing domains in a nucleus, $\langle D_{\text{PD}} \rangle$, representing the global chromatin landscape; (4) the autocorrelation function (ACF) scaling exponent, $D_{\text{ACF}}$, which captures the decay of chromatin density correlations with distance; and (5) the experimentally measured scaling from Partial Wave Spectroscopic (PWS) microscopy, $D_n$. At smaller scales, the polymeric scaling exponent distinguishes general polymer behavior from the chromatin-specific scaling seen in packing domains. However, our primary focus is on the power-law scaling behavior within packing domains, which has been linked to transcriptional activity (1). Packing domains are discrete, densely packed regions within the nucleus, where $D_{\text{PD}}$ quantifies the relationship between chromatin mass and volume at the domain level (1, 2), while $\langle D_{\text{PD}} \rangle$ reflects the overall packing domain organization at the nuclear level. Given the limitations of high-resolution electron microscopy for large-scale studies, we use alternative methods to study packing domain organization across cell populations.

The nuclear chromatin structure is effectively characterized by the autocorrelation function (ACF) (2, 3), which measures chromatin density correlations as a function of spatial separation, $N$. The ACF reveals the spatial arrangement of dense chromatin regions and more open areas throughout the nucleus, and is defined as:

$$\text{ACF}(r) = r^{D_{\text{ACF}}-3} \tag{1}$$

where $D_{\text{ACF}}$ represents the average scaling exponent. A higher $D_{\text{ACF}}$ indicates slower decay of chromatin density correlations, suggesting more uniform chromatin distribution over larger regions, while a lower $D_{\text{ACF}}$ signifies rapid decay and more compact chromatin areas. Notably, $D_{\text{ACF}}$ is shaped by both the packing domains and the inter-domain spaces that separate them. These inter-domain regions, consisting of less densely packed chromatin or unoccupied areas, significantly influence the ACF's decay rate. The combined effect of these two regions contributes to the composite exponent $D_{\text{ACF}}$, which provides a broader view of chromatin organization across the nucleus. The ACF is particularly sensitive to the volume fraction of packing domains within a nucleus (VF), underscoring the need to consider large-scale chromatin organization when studying its transcriptional role.

The scaling exponent from PWS microscopy, $D_n$, is closely related to $D_{\text{ACF}}$ but displays distinct features. Their relationship is approximated as:

$$D_n \approx D_{\text{ACF}} + \frac{\ln(\text{VF})}{b} \tag{2}$$

where $b$ is a constant determined by optical and sample properties (3). This relationship shows that $D_n$ increases logarithmically with VF, with more noticeable effects at lower chromatin densities. The PWS signal, $\Sigma$, related to $D_n$, follows a sigmoidal trend in the linear regime ($\Sigma \sim A(D_b - D_0)$), where $D_b < 3$ is a parameter derived from the Whittle-Matern model of the ACF (3). PWS measurements, conducted within a coherence volume of approximately 1 $\mu\text{m}^3$, allow for interference-based nanoscale analysis. The signal $\Sigma^2$ is proportional to the Fourier transform of the ACF, integrated over a defined region in Fourier space. When packing domains fully permeate the nucleus ($\text{VF} = 1$), a singular ACF is observed. However, when chromatin occupies both packing domains and inter-domain regions, multiple ACFs emerge. Assuming uncorrelated densities between these regions allows their Fourier transforms to be summed, giving:

$$\Sigma^2 = \Sigma^2_{\text{PD}}\text{VF} + \Sigma^2_{\text{inter}}(1 - \text{VF}) \tag{3}$$

Since the lower density of inter-domain chromatin corresponds to higher frequencies in Fourier space, it often falls outside the detection range of PWS, making its contributions negligible. For the purposes of our model, we treat $D_n$ as a close approximation of $D_{\text{ACF}}$. Therefore, in subsequent sections, $D_n$ is used to represent the experimentally derived value for calculations, though this assumption may not always hold.

**Chromatin Packing Macromolecular Crowding (CPMC) Model.** We have previously described a Chromatin Packing Macromolecular Crowding (CPMC) model which analyzes how chromatin can affect gene expression through the mechanism of macromolecular crowding (4, 5). The CPMC model considers the total chromatin ACF to calculate $D_{\text{ACF}}$, which determines the variance in molecular crowding across transcriptional interaction volumes. The correlation between gene expression levels and chromatin density is particularly significant under varying average crowding conditions. Packing domains primarily affect gene expression through crowding, as they provide environments where some genes may be exposed to more favorable conditions. We propose that regions of chromatin located near the surfaces of packing domains exhibit heightened transcriptional activity due to optimal molecular crowding conditions. The fraction of the genome residing in these transcriptionally favorable regions depends on $D_{\text{ACF}}$. The gene expression rate $E$ is related to $D_{\text{ACF}}$ through both gene accessibility and the effects of molecular crowding on transcription kinetics.

Molecular crowding, a crucial factor in gene regulation, affects transcription by altering local biochemical environments. To analytically approximate the gene expression rate, denoted as $\varepsilon$, we leverage established relationships between macromolecular crowding and mRNA production rates. This approximation builds on the Macromolecular Crowding (MC) model described by Matsuda et al. (6) and Shim et al. (7), which forms the core of the CDA model. Here, $\varepsilon$ is considered a function of both the crowding parameter, $\phi_{\text{in}}$, and a vector of molecular factors, $\vec{m}$, that includes concentrations and affinities of transcriptional reactants (TRs) such as transcription factors (TFs) and RNA polymerase II (Pol II).

***Calculation of the Impact of Crowding on Transcription using the Macromolecular Crowding (MC) Model.*** The MC model simulates how crowding affects the diffusion and binding of TRs within the nucleus. These simulations, conducted through Brownian Dynamics and Monte Carlo methods, focus on the transcriptional interaction volume (denoted in variables as "in") surrounding a gene. It is essential to note that these simulations assume spherical crowders and transcriptional proteins of uniform radius and cylindrical genes, potentially oversimplifying the nuclear environment's complexity. We use $\phi_{\text{in,model}}$ to represent the fraction of space in the interaction volume occupied by spherical crowding agents used in the MC model. The MC model takes in many variables which are involved in the process of transcription, including protein concentrations, association and dissociation rates, and binding affinities. To denote all the inputs we use for the MC model, we denote the molecular factors as $\vec{m}$, account for variables which modulate mRNA synthesis. For the purposes of the CDA model, our molecular factors $\vec{m}$ use constant values for all input rates and varying concentrations of TRs. A higher input TR concentration represents genes with high expression (which we use to look at the effect on frontloaded genes prior to stress induction) and a lower TR concentration represents genes with low expression (namely, genes that are potentially involved in plasticity and are differentially expressed to a considerable extent after exposure to a stressor). For this reason, we evaluate the effect of both crowding ($\phi_{\text{in,model}}$) and initial expression ([TR]) on the amount of mRNA produced by transcription. A full description of the methodology used can be found in the methods section under "Macromolecular Crowding Model" and the input parameters are in Table 1, both of which utilize the published framework (6, 7) Within the interaction volume, mRNA synthesis exhibits a non-monotonic response to crowding due to competing effects of increased TR localization and reduced mobility (6, 7). Optimal transcription occurs at an intermediate crowding level, $\phi_{\text{in,model,max}}$, where transcription efficiency, controlled by both $\phi_{\text{in,model}}$ and $\vec{m}$, is maximized (Fig. 1B). The MC model provides crucial insights into the relationship between crowding and mRNA production. Specifically, it reveals a non-monotonic relationship with a clear peak at an optimal crowding level. We use this fundamental shape to inform our analytical approximation, focusing on capturing the peak and curvature of this relationship in a computationally efficient manner.

It is important to note that the output of the MC model is mRNA concentration ([mRNA]). We can convert this to the number of mRNA transcripts ($N$) using the volume used for the model to calculate the concentration of the input TRs, which is $V_{cell} = 500$ μm$^3$ (the volume of a typical HeLa cell) (6), as $N = [\text{mRNA}] \times N_A \times V_{cell}$, where $N_A$ is Avogadro's constant. The steady-state expression rate $\varepsilon$ can be found with $\varepsilon = N \times \nu$, where $\nu = 3 \times 10^{-4}$ s$^{-1}$ is the degradation rate (6). While we use transcript numbers and expression rates interchangeably in subsequent discussions, it is important to note that the underlying calculations account for the conversion between concentration and number of transcripts. However, in practice, we can determine the relevant properties of the relationship between crowding and transcription as a dimensionless function, which will be defined below.

***Taylor Series Approximation of the Gene Expression Rate and Integration Over Crowding.*** To approximate gene expression as a function of crowding and molecular factors, we consider the non-monotonic shape of the gene expression rate $\varepsilon(\vec{m}, \phi_{\text{in}})$. We perform a Taylor series expansion of $\varepsilon(\vec{m}, \phi_{\text{in}})$ around the point $\phi_{\text{in,max}}$, where transcription is maximized, yielding:

$$\varepsilon(\vec{m}, \phi_{\text{in}}) \approx \varepsilon(\vec{m}, \phi_{\text{in,max}}) + \frac{1}{2} \left. \frac{\partial^2 \varepsilon(\vec{m}, \phi_{\text{in}})}{\partial \phi_{\text{in}}^2} \right|_{\phi_{\text{in}} = \phi_{\text{in,max}}} (\phi_{\text{in}} - \phi_{\text{in,max}})^2 + \dots \quad \textbf{(4)}$$

Since the first derivative vanishes at the peak, the dominant term is the second derivative. Using the value at the peak and the second derivative is often sufficient to approximate $\varepsilon(\vec{m}, \phi_{\text{in}})$, so we truncate the Taylor series and use these two terms for further calculations.

To incorporate the effects of packing domains on transcriptional sensitivity, we consider the crowding parameter $\phi_{\text{in}}$, which varies locally due to chromatin packing heterogeneity. This variability can be modeled using a Gaussian distribution for $\phi_{\text{in}}$, $f(\phi_{\text{in}})$, which captures the natural fluctuations in macromolecular crowding densities. The variance of this distribution, $\sigma^2_{\phi_{\text{in}}}$, quantifies the degree of fluctuation and is derived from the power-law scaling properties of packing domains. Specifically, it depends on the interaction radius, $r_{\text{in}}$, and the dimension of the autocorrelation function $D_{\text{ACF}}$, representing the influence of chromatin structure on crowding density:

$$\sigma^2_{\phi_{\text{in}}} = \int \text{ACF}_{\text{in}}(r) \cdot \text{ACF}_{\phi}(r) dr \approx \left(\frac{3}{D_{\text{ACF}}}\right)^C \left(\frac{r_{\text{min}}}{r_{\text{in}}}\right)^{3-D_{\text{ACF}}} \sigma_o^2 \quad \textbf{(5)}$$

where $r_{\text{in}} = r_{\text{min,in}} + r_{\text{min}}(L/A_v)^{1/D_{\text{ACF}}}$, with $r_{\text{min}} = 1$ nm as the radius of the elementary chromatin unit (i.e., a DNA base pair), $r_{\text{min,in}} = 15$ nm representing the interaction volume for a single base pair, and $L = 6$ kbp as the gene length. Additionally, $\sigma_o^2 = \phi_c(1-\phi_c)(1-\phi_m)^2$, where $\phi_c$ is the crowding contribution due to chromatin (the CVC) and $\phi_m$ is the amount of crowding due to other mobile crowders. $D_{\text{ACF}}$, the dimension of the autocorrelation function, reflects how chromatin structure influences crowding density at different length scales. A higher $D_{\text{ACF}}$ indicates more uniform chromatin distribution, while a lower value suggests more heterogeneous packing, directly impacting local crowding variations.

The Gaussian distribution of crowding densities reflects how chromatin packing creates regions of varying crowding levels that influence transcriptional efficiency. These variations, represented by $\sigma^2_{\phi_{\text{in}}}$, are particularly important in capturing the crowding effects on transcription rates. We integrate $\varepsilon(\overrightarrow{m}, \phi_{\text{in}})$ over this Gaussian distribution using $\overline{\varepsilon} = \int \varepsilon(\overrightarrow{m}, \phi_{\text{in}}) f(\phi_{\text{in}}) d\phi_{\text{in}}$. We determine the average gene expression rate, $\overline{\varepsilon}$:

$$\overline{\varepsilon} \approx \varepsilon(\overrightarrow{m}, \phi_{\text{in,max}}) + \frac{1}{2}\sigma^2_{\phi_{\text{in}}} \left.\frac{\partial^2 \varepsilon(\overrightarrow{m}, \phi_{\text{in}})}{\partial \phi^2_{\text{in}}}\right|_{\phi\text{in}=\phi_{\text{in,max}}} \quad \textbf{(6)}$$

The second derivative approximates the sensitivity of gene expression to local crowding fluctuations and can be found using:

$$\left.\frac{\partial^2 \varepsilon(\overrightarrow{m}, \phi_{\text{in}})}{\partial \phi^2_{\text{in}}}\right|_{\phi_{\text{in}}=\phi_{\text{in,max}}} \approx -\sqrt{\varepsilon(\overrightarrow{m}, \phi_{\text{in,max}})\kappa(\overrightarrow{m})} \quad \textbf{(7)}$$

where $\kappa(\overrightarrow{m})$ is a fitting parameter that depends on molecular factors and quantifies the steepness of the gene expression curve near $\phi_{\text{in,max}}$. The parameter $\kappa(\overrightarrow{m})$ is estimated by analyzing the MC model results for various molecular factor combinations. It quantifies how sharply the gene expression rate changes around $\phi_{\text{in,max}}$. In practice, $\kappa(\overrightarrow{m})$ is determined by fitting the analytical approximation to the MC model output for each set of molecular factors.

Incorporating these terms, the average gene expression rate becomes:

$$\overline{\varepsilon} \approx \varepsilon(\overrightarrow{m}, \phi_{\text{in,max}}) - \frac{1}{2}\sigma^2_{\phi_{\text{in}}} \sqrt{\varepsilon(\overrightarrow{m}, \phi_{\text{in,max}})\kappa(\overrightarrow{m})} \quad \textbf{(8)}$$

This approximation captures two key biological aspects: (1) the base expression level at optimal crowding (first term), and (2) the reduction in expression due to local crowding fluctuations (second term). The interplay between these terms reflects how chromatin organization can modulate gene expression sensitivity.

To quantify the sensitivity of gene expression to variations in transcriptional reactant concentrations, we utilize linear regression to estimate the parameter $\kappa$. This parameter characterizes the slope of the relationship between the second derivative of the gene expression rate and the transcriptional reactant concentration, specifically in regions where the concentration exceeds a predetermined threshold of 0.002 nM. The analysis begins by defining the fitting region based on the transcriptional reactant concentration. Data points within this range are selected for regression, ensuring that only relevant data is included in the calculation of $\kappa$. The linear regression is performed using the least squares method, yielding the slope (i.e., $\kappa$; Fig. 1). The fitted linear relationship is expressed as:

$$\left.\frac{\partial^2 \varepsilon(\overrightarrow{m}, \phi_{\text{in}})}{\partial \phi^2_{\text{in}}}\right|_{\phi\text{in}=\phi_{\text{in,max}}} = \kappa \cdot \varepsilon(\overrightarrow{m}, \phi_{\text{in,max}}) + b, \quad \textbf{(9)}$$

where $\varepsilon''$ represents the second derivative of the gene expression rate, $[TR]$ denotes the transcriptional reactant concentration, and $b$ is the intercept.

***Derivation of Gene Expression Sensitivity to packing domain Characteristics.*** Building upon the approximation of $\overline{\varepsilon}$ developed in the previous section, we present a comparative analysis of gene expression sensitivity to various packing domain parameters by extending the CPMC model (5). In our model, we distinguish between the local expression rate ($\varepsilon$) and the ensemble expression rate ($E$). The local expression rate $\varepsilon$ represents the mRNA production rate within the interaction volume surrounding a specific gene, influenced by local chromatin packing characteristics. In contrast, the ensemble expression rate $E$ reflects the overall transcriptional output of a gene, considering both the local chromatin environment and larger-scale nuclear organization. The relationship between $\varepsilon$ and $E$ can be expressed as $E = \overline{\varepsilon} \cdot ER$, where $\overline{\varepsilon}$ is the average expression rate of an ensemble of genes with similar molecular regulators of transcription per unit of DNA, and $ER$ (Exposure Ratio) represents the accessible surface of a gene, given by $ER = A_{ER}(N_{\text{PD}}/A_v)^{-1/D_{\text{PD}}}$ (2). This formulation captures how both local crowding effects and larger-scale chromatin accessibility influence gene expression. It is essential to note that while $\varepsilon$ primarily reflects local chromatin effects, $E$ incorporates both local and global influences on gene expression. This distinction allows our model to capture the complex interplay between different scales of nuclear organization in determining transcriptional output. However, the relationship between $\varepsilon$ and $E$ is an approximation, as it assumes a degree of independence between local and global effects that may not always hold in the highly interconnected nuclear environment.

Transcriptional responsiveness, defined as the ratio of final to initial expression rates ($E_2/E_1$), is governed by three physical regulators of transcription: average nuclear packing domain scaling behavior ($D_n$), genomic size ($N_{\text{PD}}$), and crowding ($\phi_{\text{in}}$). We employ analytical relations to quantify the sensitivity of gene expression to each of these regulatory factors, denoting the dependence of sensitivity on average expression rate ($\overline{\varepsilon}$) and exposure ratio ($ER$) as $Se_{\overline{\varepsilon}}$ and $Se_{ER}$, respectively. The sensitivity of gene expression to $D_n$, denoted as $Se_{E,D_n}$, is defined as:

$$Se_{\overline{\varepsilon},D_n} = \frac{\partial \ln \overline{\varepsilon}}{\partial \ln D_n} \approx -G(\overline{\varepsilon}) \left[ D_n \ln\left(\frac{r_{\text{in}}}{r_{\text{min}}}\right) + \frac{3-D_n}{D_n}\frac{r_{\text{min}}}{r_{\text{in}}}\left(\frac{L}{A_v}\right)^{1/D_n} \ln\left(\frac{L}{A_v}\right)\right] + \frac{3}{D_n}\ln\left(\frac{N_{\text{PD}}}{A_v}\right)\left[\frac{1-2\phi_c}{1-\phi_c}\right] \quad \textbf{(10)}$$

$$Se_{ER,D_n} = \frac{\partial \ln ER}{\partial \ln D_n} \approx \frac{1}{D_n}\ln\left(\frac{N_{\text{PD}}}{A_v}\right) \quad \textbf{(11)}$$

$$Se_{E,D_n} = Se_{\overline{\varepsilon},D_n} + Se_{ER,D_n} \quad \textbf{(12)}$$

Here, $\phi_c = A_v\left(\frac{N_{\text{PD}}}{A_v}\right)^{1-3/D_n}$ and $G(\overline{\varepsilon})$ is a dimensionless function. The expression is given by:

$$G(\overline{\varepsilon}) = \frac{\kappa}{8\overline{\varepsilon}}(\sigma^2_{\phi_{\text{in}}})^2\left(1+\sqrt{1+\frac{16}{(\sigma^2_{\phi_{\text{in}}})^2}\frac{\overline{\varepsilon}}{\kappa}}\right) \quad \textbf{(13)}$$

For most physiological scenarios, where $\frac{16}{\sigma^2_{\phi_{\text{in}}}} \gg 1$, $G(\overline{\varepsilon})$ can be approximated as $G(\overline{\varepsilon}) \approx \frac{1}{2}\sigma^2_{\phi_{\text{in}}}\sqrt{\frac{\kappa}{\overline{\varepsilon}}}$.

For domain size $N_{\text{PD}}$, the sensitivity equations are as follows:

$$Se_{\overline{\varepsilon},N_{\text{PD}}} = \frac{\partial \ln \overline{\varepsilon}}{\partial \ln N_{\text{PD}}} \approx -G(\overline{\varepsilon})\left(1-\frac{3}{D_n}\right)\left[\frac{1-2\phi_c}{1-\phi_c}\right] \quad \textbf{(14)}$$

$$Se_{ER,N_{\text{PD}}} = \frac{\partial \ln ER}{\partial \ln N_{\text{PD}}} \approx -\frac{1}{D_n} \quad \textbf{(15)}$$

$$Se_{E,N_{\text{PD}}} = Se_{\overline{\varepsilon},N_{\text{PD}}} + Se_{ER,N_{\text{PD}}} \quad \textbf{(16)}$$

Finally, for the volume packing efficiency of a packing domain $A_v$:

$$Se_{\overline{\varepsilon},A_v} = \frac{\partial \ln \overline{\varepsilon}}{\partial \ln A_v} \approx -G(\overline{\varepsilon})\left(\frac{3}{D_n}-1\right)\left[\frac{1-2\phi_c}{1-\phi_c}+\frac{A_v}{L}\frac{r_{\text{min}}}{r_{\text{in}}}\right] \quad \textbf{(17)}$$

$$Se_{ER,A_v} = \frac{\partial \ln ER}{\partial \ln A_v} \approx \frac{1}{D_n} \quad \textbf{(18)}$$

$$Se_{E,A_v} = Se_{\overline{\varepsilon},A_v} + Se_{ER,A_v} \quad \textbf{(19)}$$

As the CPMC model predicts, an increase in $D_{\text{ACF}}$ (or $D_n$) can lead to differential gene expression dynamics, with higher $D_{\text{ACF}}$ promoting the upregulation of highly expressed genes while downregulating genes with lower expression. To explore the impact of the packing domain characteristics as physical regulators of transcription, we examined experimental data obtained from ChromSTEM analysis of HCT116 cells (2). This high-resolution imaging technique allowed us to directly measure the distribution of packing domain characteristics ($D_{\text{PD}}$, $N_{\text{PD}}$, and $A_v$) across a cell nucleus. Using the $25^{\text{th}}$, $50^{\text{th}}$, and $75^{\text{th}}$ percentiles of domain-specific $D_{\text{PD}}$, $N_{\text{PD}}$, and $A_v$ to represent the range of the nuclear average parameters (Fig. 2), our analysis demonstrates that gene expression exhibits significantly higher sensitivity to changes in packing domain scaling behavior ($Se_{E,D_n}$) compared to changes in domain size ($Se_{E,N_{\text{PD}}}$) or average packing efficiency ($Se_{E,A_v}$), with differences in responsiveness spanning 1 to 2 orders of magnitude. This finding underscores the critical role of packing domain scaling ($D_n$) in determining cellular adaptability and response to stressors, justifying our focus on $D_n$ in the CDA model and subsequent experiments.

**Chromatin-Dependent Adaptability (CDA) Model.** We introduce the Chromatin-Dependent Adaptability (CDA) model, which leverages the established relationships between gene expression and chromatin organization to predict cell survival probabilities in response to stressors. This model integrates chromatin packing states with transcriptional responsiveness, providing a quantitative framework for understanding cellular adaptability. The CDA model posits that a cell population's survival probability under cytotoxic stress is influenced by the average nuclear packing domain organization, which modulates transcriptional responses. Importantly, the model does not suggest that all cells will survive all stressors through specific gene regulation; rather, it evaluates survival in a statistical context, reflecting the average initial chromatin packing state that encapsulates the crowding conditions of all genes within the cell.

***Derivation of*** $\Theta$. The regulation of transcripts for the gene(s) of interest is given by $x = N_2/N_1$, where $N_1$ and $N_2$ are the number of mRNA transcripts before and after stress exposure, respectively. We postulate that the probability density function (PDF) of the transcript regulation follows a log-normal distribution:

$$\text{PDF}(x) \approx \frac{1}{s\sqrt{2\pi}x} \mathrm{e}^{-\frac{\ln(x/m)^2}{2s^2}} \tag{20}$$

where $m$ represents the median of the distribution and $s$ is the shape parameter, which can also be interpreted as the standard deviation of $\ln(x)$, capturing the spread of the transcript regulation. When $s \ll 1$, $\ln(m) \approx \mu$, where $\mu$ represents the mean and $s$ approximates the coefficient of variation (COV) of transcript upregulation.

We hypothesize that a cell's survival, when exposed to a cytotoxic stressor within a critical time period, depends on the up-regulation of specific stress response genes exceeding a threshold level $x_{\text{crit}}$. This leads us to assume that the probability of survival as a function of $x$ can be approximated by a step function: if $x > x_{\text{crit}}$ the cell survives, and if $x < x_{\text{crit}}$ the cell dies. The resulting probability of cell death, $\Theta$, is defined as the cumulative distribution function (CDF) of this threshold $x_{\text{crit}}$:

$$\Theta(x_{\text{crit}}) = \text{CDF}(x_{\text{crit}}) = \frac{1}{2}\text{erfc}\left(\frac{\ln(m/x_{\text{crit}})}{\sqrt{2}s}\right) \tag{21}$$

with the complementary error function defined as $\text{erfc}(u) = 1 - \text{erf}(u) = \frac{2}{\sqrt{\pi}}\int_u^\infty e^{-v^2} dv$. Here, the use of erfc arises naturally from the log-normal distribution, as it corresponds to the tail probability beyond $x_{\text{crit}}$.

To facilitate numerical and analytical computation, we approximate the log-normal CDF using a Hill equation, which provides a more tractable form:

$$\Theta(x_{\text{crit}}) \approx \frac{1}{1 + (k/x_{\text{crit}})^h} \tag{22}$$

where the parameter $k \approx m$ represents the malleability of the response, indicating the median level of transcript regulation required for survival, and $h \approx 3/(s\sqrt{\pi})$ represents the Hill coefficient, which is inversely related to the COV or heterogeneity of the response. This approximation captures the sigmoidal nature of the cell survival curve as a function of $x_{\text{crit}}$, allowing for easier computation while preserving the essential features of the original log-normal distribution.

We now generalize this formulation to compare the survival probabilities of two cells, labeled $a$ and $b$. As discussed in the main text (see Fig. 1), comparing the upregulation of cell $a$ with that of cell $b$, we can calculate the death probability of cell $b$ as follows:

$$\Theta_b(x_{\text{crit}}) \approx \frac{1}{1 + (k_a\gamma_k/x_{\text{crit}})^{\frac{h_a}{\gamma_h}}} \tag{23}$$

where $\gamma_k = k_b/k_a$ and $\gamma_h = h_a/h_b$. Here, $\gamma_k$ represents the ratio of the malleability of the upregulation response between cell $b$ and cell $a$ (i.e., the ratio of the median transcript regulation levels $N_{2a,b}/N_{1a,b}$), and $\gamma_h$ represents the ratio of the COVs of the upregulation of the two cells, indicating the relative heterogeneity in the transcript regulation responses. These ratios, $\gamma_k$ and $\gamma_h$, account for differences in the transcript dynamics between the two cells, enabling a comparative survival analysis.

However, determining the exact threshold $x_{\text{crit}}$ experimentally for all relevant pro-survival genes in cell $b$ can be challenging. An alternative method of calculating the death probability in cell $b$ is to use the known death probability for cell $a$ to solve for $x_{\text{crit}}$. Given Eq. 22, we can rewrite Eq. 23 as:

$$\Theta_b(\Theta_a) = \frac{1}{1 + \gamma_k^{h_b}(\frac{1}{\Theta_a} - 1)^{\frac{1}{\gamma_h}}} \tag{24}$$

In this formulation, $\Theta_b$ is directly related to $\Theta_a$, allowing us to compute the death probability of cell $b$ using the known survival characteristics of cell $a$. This equation simplifies comparative analyses between different cell populations.

***Incorporating Temporal Dynamics.*** The CPMC model's versatility allows application to both steady-state and non-steady-state conditions. For steady-state, the number of mRNA transcripts ($N$) is not time-dependent, i.e., $\frac{\partial N}{\partial t} = 0$. In this scenario, CPMC predicts $N$ by determining the expression rate $E$ and using the relation $E = N\nu$ for the mRNA degradation rate $\nu$ (6). However, many biological processes, including responses to chemotherapeutic agents, occur under non-steady-state conditions.
To address these conditions, we developed an extended formalism. Let a cell population be exposed to a stressor at time $t = 0$. The number of transcripts at time $t$ is determined by:

$$\frac{\partial N}{\partial t} = \frac{\partial E}{\partial t}(1 - \mathrm{e}^{-t/\tau}) \tag{25}$$

where $\tau$ is the mRNA elimination time constant, calculated from the half-life of mRNA, $\tau_{1/2}$, as $\tau = \frac{1}{\ln 2}\tau_{1/2}$. Based on existing literature, we adopt $\tau_{1/2} = 10$ hours (8). This differential equation captures the delay in transcript accumulation as the system transitions from initial exposure to a new steady state.
We approximate $\frac{\partial N}{\partial t} \approx \frac{N_2(t) - N_1}{N_1}$ as the relative change in transcripts and $\frac{\partial E}{\partial t} \approx \frac{E_2(t) - E_1}{E_1}$ as the relative change in mRNA production rate. Both $N_2(t)$ and $E_2(t)$ are time-dependent after stress exposure, reflecting the dynamic nature of the cellular response. Defining $\beta = \frac{E_2(t)}{E_1}$ and $k = \frac{N_2(t)}{N_1}$, we describe transcriptional malleability with:

$$k(t) = 1 + (\beta - 1)(1 - \mathrm{e}^{-t/\tau}) \tag{26}$$

such that $k(t)$ evolves over time, approaching $\beta$ as $t \to \infty$, indicating eventual transcript upregulation.
If we know $\beta_a$, the average upregulation of the expression rate for cell $a$, we can predict $\beta_b$ using:

$$\delta = \frac{\beta_b}{\beta_a} = \exp\left(\int_{D_{n,a}}^{D_{n,b}} Se(\beta_a E_1, D'_n) - Se(E_1, D'_n)\frac{dD'_n}{D'_n}\right) \tag{27}$$

to link differences in gene expression to chromatin structure via $D_n$. We compute $k_b = k_a\gamma_k$ by substituting $\beta_b = \delta\beta_a$ into Eq. 26, yielding the transcriptional malleability for cell $b$.
To account for evolving heterogeneity in gene expression over time, we developed an equation integrating both initial variability and stress-induced changes:

$$s(t) = COV\left[x = \frac{N_2(t)}{N_1}\right] = \beta\frac{1 - \mathrm{e}^{-t/\tau}}{k(t)}\sqrt{COV\left[E_1\right]^2 + COV\left[E_2\right]^2} \tag{28}$$

where $COV[E_1] = \sqrt{2}G(\bar{\varepsilon}_i)$ and $\bar{\varepsilon}_i$ is the expression rate of a given gene per unit of DNA pre- ($i = 1$) and post- ($i = 2$) stimulus. Assuming transcriptional heterogeneity is negligible prior to treatment ($COV[E_2] \gg COV[E_1]$), we express the time-dependent heterogeneity for cells $a$ and $b$ as:

$$s_a(t) = \beta_a\frac{1 - \mathrm{e}^{-t/\tau}}{k_a(t)}\sqrt{2}G\left(\bar{\varepsilon}_{2,a}\right) \tag{29}$$

$$s_b(t) = \beta_b\frac{1 - \mathrm{e}^{-t/\tau}}{k_b(t)}\sqrt{2}G\left(\bar{\varepsilon}_{2,b}\right) \tag{30}$$

where $\bar{\varepsilon}_{2,a} = \beta_a\bar{\varepsilon}_{1,a}$ and $\bar{\varepsilon}_{2,b} = \beta_b\bar{\varepsilon}_{1,b}$. These expressions indicate how heterogeneity evolves in each cell over time, modulated by changes in gene expression rates and transcript numbers.
If $\bar{\varepsilon}_{1,a}$ is known, the expression rate $\bar{\varepsilon}_{1,b}$ can be found using the relation $\gamma_{\bar{\varepsilon}_i} = \bar{\varepsilon}_{i,b}/\bar{\varepsilon}_{i,a}$. Here, $\gamma\bar{\varepsilon}_i$ represents the average change in expression rate for cell $a$ compared to cell $b$ before ($i = 1$) or after ($i = 2$) treatment with a cytotoxic stressor, calculated using $Se_{\bar{\varepsilon},D_n}$:

$$\gamma_{\bar{\varepsilon}} = \frac{\bar{\varepsilon}_b}{\bar{\varepsilon}_a} = \frac{E_b}{E_a}\frac{ER(D_{n,a})}{ER\left(D_{n,b}\right)} = N_{\mathrm{PD}}^{\frac{1}{D_{n,b}} - \frac{1}{D_{n,a}}}\exp\left(\int_{D_{n,a}}^{D_{n,b}} Se(\bar{\varepsilon}_a, D'_n)\frac{dD'_n}{D'_n}\right) \tag{31}$$

Thus, we can determine the expression rate in cell $b$ after treatment, $\bar{\varepsilon}_{2,b}$, using:

$$\bar{\varepsilon}_{2,b} = \bar{\varepsilon}_{1,a}\beta_a\gamma_{\bar{\varepsilon}_2} = \bar{\varepsilon}_{1,a}\delta\beta_a\frac{p_{g,a}}{p_{g,b}} = \bar{\varepsilon}_{1,a}\delta\beta_a N_{\mathrm{PD}}^{\frac{1}{D_{n,b}} - \frac{1}{D_{n,a}}} \tag{32}$$

where $p_{g,a}$ and $p_{g,b}$ represent gene accessibility in cells $a$ and $b$, respectively, inversely related to chromatin packing density. The gene accessibility ratio $\frac{p_{g,a}}{p_{g,b}}$ is expressed as a function of $N_{\mathrm{PD}}$ and $D_n$. Substituting Eqs. 29, 30, 31, and 32 into Eq. 28, we calculate the heterogeneity ratio, $h_b = 3/(s_b\sqrt{\pi}) = h_a/\gamma_h$, necessary for computing the death probability $\Theta(x_{\mathrm{crit}})$ in Eq. 23.

***Evaluating Model Parameters.*** To predict the cell death probability $\Theta$, the CDA model requires estimation of several key parameters: (1) the gene upregulation factor $\beta$; (2) the survival threshold $x_{\text{crit}}$; (3) the relative initial expression of the upregulated gene(s) $\ln(E_1/\overline{E}_1)$; (4) the critical decision time point $T_{\text{crit}}$. As a function of $D_n$, $\Theta$ follows a sigmoidal curve where cells with low $D_n$ have a close to 100% probability of death when they encounter a stressor and cells with high $D_n$ having close to 0% probability (Figs. 1D-E and 3) The two main parameters that are key to the trends for $\Theta$ are $\beta$ and $x_{\text{crit}}$, which shift the probability of death for high $D_n$ cells and the inflection point of the sigmoidal curve, respectively (Figs. 1D-E). Two other parameters that have less of an impact on model predictions are $\ln(E_1/\overline{E}_1)$ and $T_{\text{crit}}$.
The critical timepoint for the cell death decision, $T_{\text{crit}}$, is an important parameter in the CDA model. Previous studies have shown that cells typically exhibit signs of apoptosis 5-10 hours post-chemotherapy treatment, with apoptosis induction varying significantly depending on the dosage (9–11). Here, we use $T_{\text{crit}} = 7$ hours. Increasing $T_{\text{crit}}$ changes the sigmoidal curve so that cells with lower $D_n$ have a lower probability of death, as illustrated in Fig. 3, which shows the relationship between chromatin packing scaling ($D_n$) and cell death probability for different $T_{\text{crit}}$ values.

***Incorporating Population Distribution of $D_n$.*** Our experimental results reveal that cell populations exhibit heterogeneous responses to chemotherapy, characterized by a distribution of $D_n$. Cells with extreme values of $D_n$ are particularly sensitive to chemotherapy, necessitating a model that incorporates this variability. To represent this distribution, we define a probability distribution function (PDF) that spans a range of $D_n$ values, allowing us to capture the full spectrum of cell responses.
We introduce $n_\tau$ as the number of cell doubling intervals since exposure to the stressor, with $\tau_2$ representing the characteristic cell doubling time. Cells decide whether to undergo apoptosis or division based on a comparison between the critical time for apoptosis, $T_{\text{crit}}$, and the doubling time, $\tau_2$. Specifically, when $T_{\text{crit}} < \tau_2$, the decision time after $n_\tau$ doublings is given by $t_{n_\tau} = n_\tau \cdot \tau_2$.
The probability of cell survival, $P_s$, is inversely related to the probability of cell death, $\Theta$, through the relationship $P_s = 1 - \Theta$. For a population with a distribution of $D_n$ values, the overall survival probability is determined by integrating over the population distribution:

$$P_s(n_\tau) = \int \text{PDF}(D_n)\left[1 - \Theta(D_n)\right]^{n_\tau} dD_n \qquad \textbf{(33)}$$

This formulation highlights how the survival probability declines with successive cell divisions, with the rate of decline modulated by each cell's specific $D_n$ value.
To determine the population-averaged $D_n$, $\langle D_n \rangle$, we compute a weighted average of $D_n$, where the weighting is determined by the survival probability:

$$\langle D_n(t_{n_\tau}) \rangle = \frac{\int D_n \cdot \text{PDF}(D_n)[1 - \Theta(D_n)]^{n_\tau} dD_n}{\int \text{PDF}(D_n)[1 - \Theta(D_n)]^{n_\tau} dD_n} \qquad \textbf{(34)}$$

This expression provides a time-dependent measure of $\langle D_n(t_{n_\tau}) \rangle$, accounting for how cell survival evolves through repeated divisions under stress.
Next, we introduce the effective inhibition rate (EIR), which quantifies the cumulative effect of chemotherapy on the cell population. The EIR describes the rate at which the survival probability decays over time, modeled as an exponential decay with $P_s = \exp(-t_{n_\tau} \cdot EIR)$. For a homogeneous population where all cells have the same $D_n$ value (i.e., $\text{PDF}(D_n) = \delta(D_n - D_{n,a})$), we derive the following expression for the EIR:

$$EIR(t_{n_\tau}) = \frac{1}{t_{n_\tau}} \ln\left(\frac{1}{\left[1 - \Theta(D_{n,a})\right]^{n_\tau}}\right) \qquad \textbf{(35)}$$

To assess the rate of inhibition per doubling interval, we define the incremental inhibition rate (IR):

$$IR(n_\tau) = \frac{1}{t_1} \ln\left(\frac{P_s(n_\tau)}{P_s(n_\tau + 1)}\right) \qquad \textbf{(36)}$$

where $t_1 = \tau_2$. In the case of a homogeneous population, where $\text{PDF}(D_n) = \delta(D_n - D_{n,a})$, we find that $IR(n_\tau) = EIR(n_\tau)$.
Assuming that cell proliferation rates are independent of $D_n$, the total number of cells at time $t$ without chemotherapy follows an exponential growth law such that $N(t) = N(0)\exp(t\ln(2)/\tau_2)$. When chemotherapy is applied, the growth rate is modified by the inhibition rate, yielding:

$$\frac{N(t)}{N(0)} = \exp\left(\frac{t\ln(2)}{\tau_2} - t \cdot IR(t)\right) \qquad \textbf{(37)}$$

Experimental measurements of $P_s$ must therefore be normalized by the expected number of cells at time $t$ in the absence of stress, $\exp(t\ln(2)/\tau_2)$, to account for natural cell proliferation.

Finally, we express the full inhibition rate as a function of time:

$$IR(t) = \frac{1}{P_s(t)}\frac{\partial P_s(t)}{\partial t} = \frac{1}{t_1}\int \mathrm{PDF}(D_n)\left[1-\Theta(D_n)\right]^{t/t_1}\ln\left[1-\Theta(D_n)\right]dD_n \tag{38}$$

This equation captures the cumulative effect of stress on the population, incorporating the heterogeneity in $D_n$. By integrating the survival probabilities across the distribution of $D_n$ values, this model provides a more comprehensive understanding of how chemotherapy influences the population dynamics over time.

**Estimation of Cell Division Induced Drift in Population $D_n$.** To accurately predict cell death probability as a function of $D_n$, we developed a comprehensive approach that models the effects of both time and cell division on chromatin packing domains. In theory, if $D_n$ remained the same in daughter cells after cell division, $\Theta$ could be directly estimated simply by looking at the difference in the PDFs of $D_n$ at two time points, as long as the growth rate is known. However, we have previously seen that there is some drift in $D_n$ when daughter cells are compared to the parent cell (1), indicating the need to take cell division into account when estimating experimental $\Theta$. Therefore, we can define the change in the number of cells using the equation:

$$\frac{dN(D_n,t)}{dt} = \frac{\ln 2}{\tau_2}\int N(D_n',t)\cdot f(D_n - D_n')\,dD_n' - N(D_n,t)\cdot IR(D_n,t) \tag{39}$$

where $N$ is the cell number as a function of $D_n$ and time $t$, $\tau_2$ is the doubling time, $f(D_n - D_n')$ is the probability of a daughter cell having $D_n$ if the mother cell had $D_n'$, and $IR$ is the inhibition rate for a cell with $D_n$ at time $t$. While $N$, $D_n$, and $IR$ are relatively simple to determine experimentally, $f(D_n - D_n')$ requires more complex estimation.

***Definition of Drift in $D_n$.*** To isolate the effect of cell division on $D_n$, we first determined all of the separate factors that impact population drift, including (1) intrinsic variation within a cluster, (2) normal temporal changes, and (3) the impact of cell division. We start by representing the overall drift as a normally distributed PDF $f(\Delta D_n)$ which is the change in $D_n$ between two time points (before and after cell division) with a mean of 0 and a variance of $\mathrm{Var}[\Delta D_n]$. To find $\mathrm{Var}[\Delta D_n]$, we first start by defining the $D_n$ of a cluster of cells before ($D_{n,1}$) and after ($D_{n,2}$) cell division as:

$$D_{n,1} = \frac{1}{N_{\text{non}} + N_{\text{div}}}\left(\sum_{i=1}^{N_{\text{non}}} D_{n,1i} + \sum_{i=N_{\text{non}}+1}^{N_{\text{non}}+N_{\text{div}}} D_{n,1i}\right) \tag{40}$$

$$D_{n,2} = \frac{1}{N_{\text{non}} + 2N_{\text{div}}}\left(\sum_{i=1}^{N_{\text{non}}} D_{n,2i} + \sum_{i=N_{\text{non}}+1}^{N_{\text{non}}+N_{\text{div}}} D_{n,2i} + \sum_{i=N_{\text{non}}+N_{\text{div}}+1}^{N_{\text{non}}+2N_{\text{div}}} D_{n,2i}\right) \tag{41}$$

where $N_{\text{non}}$ is the number of non-dividing cells, $N_{\text{div}}$ is the number of dividing cells, $D_{n,1i}$ is the $D_n$ of cell $i$ in a cluster before division, $D_{n,2i}$ is the $D_n$ of cell $i$ in the same cluster after division.
The $D_n$ after cell division ($D_{n,2i}$) for dividing cells within a cluster ($i > N_{\text{non}}$) is:

$$D_{n,2i} = \overline{D}_{n,1i} + \delta D_{n,i} + \delta D_{n,2i}^{t} \tag{42}$$

where $\overline{D}_{n,1i}$ represents the time-averaged $D_n$ of the parent cell, $\delta D_{n,i}$ is the spread in $D_n$ due to division, and $\delta D_{n,2i}^{t}$ denotes the temporal fluctuation of $D_n$. Additionally, $\forall ij$, $D_{n,ji} = \overline{D}_{n,ji} + \delta D_{n,ji}^{t}$, as we can find the $D_n$ of any cell within the cluster before or after cell division given the parent cell $D_n$ and the temporal drift.
Combining Eqs. 40, 41, and 42, we can calculate $\Delta D_n = D_{n,2} - D_{n,1}$ as:

$$\Delta D_n = \sum_{i=1}^{N_{\text{non}}} \overline{D}_{n,1i}\left(\frac{1}{N_2} - \frac{1}{N_1}\right) + \sum_{i=1}^{N_{\text{non}}} \delta D_{n,2i}^{t}\frac{1}{N_2} - \sum_{i=1}^{N_{\text{non}}} \delta D_{n,1i}^{t}\frac{1}{N_1} + 2\sum_{i=N_{\text{non}}+1}^{N_{\text{non}}+N_{\text{div}}} \overline{D}_{n,1i}\left(\frac{1}{N_2} - \frac{1}{2N_1}\right) \tag{43}$$

$$+ \sum_{i=N_{\text{non}}+1}^{N_2} \delta D_{n,i}\frac{1}{N_2} + \sum_{i=N_{\text{non}}+1}^{N_2} \delta D_{n,i}^{t}\frac{1}{N_2} - \sum_{i=N_{\text{non}}+1}^{N_1} \delta D_{n,i}^{t}\frac{1}{N_1} \tag{44}$$

where $N_1 = N_{\text{non}} + N_{\text{div}}$ is the number of cells before cell division occurs and $N_2 = N_{\text{non}} + 2N_{\text{div}}$ is the number after.
From Eq. 43, we can now determine $\mathrm{Var}[\Delta D_n]$:

$$\mathrm{Var}[\Delta D_n] = \mathrm{Var}[\overline{D}_n]\frac{N_{\text{non}}N_{\text{div}}}{N_1 N_2^2} + \mathrm{Var}[\delta D_n^{t}]\left(\frac{1}{N_2} + \frac{1}{N_1}\right) + \mathrm{Var}[\delta D_n]\frac{2N_{\text{div}}}{N_2^2} \tag{45}$$

where $\text{Var}[\overline{D}_n]$ is the variance within a cluster, $\text{Var}[\delta D_n^t]$ is the spread from the same cell over several time points, and $\text{Var}[\delta D_n]$ is the variance induced by cell division. Each of these variances can be determined from untreated cell clusters that are tracked through cell division. Specfically, the variance in $D_n$ across all cells within a cluster is $\text{Var}[D_{n,i}] = \text{Var}[\overline{D}_n] + \text{Var}[\delta D_n^t]$. Therefore, if we isolate clusters of cells that do not divide in our experiment, we can extract the values and use them to determine the remaining variance in $D_n$ that is induced by cell division $\text{Var}[\delta D_n]$.

***Identifying Temporal Variance in Non-Dividing Clusters.*** Clusters were classified as dividing or non-dividing based on their growth ratios between two consecutive time points. Clusters that exhibited less than a 25% increase in cell count between the two time points were labeled as non-dividing. We performed a linear regression analysis to find the amount of variance in $D_{n,2}$ (the value after cell division) that is explained by $D_{n,1}$, which can be determined through the correlation coefficient. For each cluster, we calculated the mean and standard deviation of $D_n$ values, normalizing these values by subtracting the mean $D_n$ for all clusters at each time point $D_{n,\text{norm}} = D_n - \langle D_n \rangle$. The coefficient of determination ($N_{\text{non}}^2$) for this regression was calculated as 0.4081, indicating that 40.81% of the variance in the second time point's $D_n$ values could be explained by the values from the first time point. The rest of the variance in the second time point can therefore be estimated to be due to intrinsic cluster variations.

***Determining Variance from Cell Division Using Dividing Clusters.*** As we determined the amount of variance in a cluster that occurs due to natural temporal effects ($N_{\text{non}}^2$), we used this to find the temporal variance $\text{Var}[\delta D_n^t]$ with the variance of the population of dividing cells before cell division:

$$\text{Var}[\delta D_n^t] = \left(N_{\text{non}}^2\right) \text{Var}[D_{n,1,\text{div}}] \tag{46}$$

Additionally, given that the remaining variance is intrinsic cluster variations ($\text{Var}[D_{n,i}] = \text{Var}[\overline{D}_n] + \text{Var}[\delta D_n^t]$), we can determine the variance within a cluster using:

$$\text{Var}[\overline{D}_n] = \left(1 - N_{\text{non}}^2\right) \text{Var}[D_{n,1,\text{div}}]. \tag{47}$$

We can now use Eq. 45 to determine the drift that occurs due to cell division if we find $\text{Var}[\Delta D_n]$, the overall variance that occurs at after cell division due to all three variances. Given that $\Delta D_n = D_{n,2} - D_{n,1}$, we can find the population variance with:

$$\text{Var}[\Delta D_n] = \frac{1}{N_{\text{div}}} \sum_{i=1}^{N_{\text{div}}} (\Delta D_{n,i} - \overline{\Delta D}_{n,i})^2 \tag{48}$$

We then used the values from Eqs. 46, 47, and 48 in Eq. 45 to determine $\text{Var}[\delta D_n]$, given that $N_{\text{non}} = 2.27$, $N_{\text{div}} = 4.45$, $N_1 = 6.09$, and $N_2 = 11.18$. We found that the PDF describing the drift in $D_n$ due to cell division $f(D_n - D_n')$ can be modeled as being centered around 0 with a standard deviation of $0.0685 \pm 0.250$. For the CDA model, we perform a convolution of the input $\text{PDF}(D_n)$ with $f(D_n - D_n')$ to arrive at more accurate predictions of $D_n$ after exposure to a stressor.

**Closed-Form Approximations of Cell Death Probability.** The non-steady-state CPMC model assumes an exponential decay in mRNA transcript numbers over time (Eq. 25). As a result, transcriptional malleability $k$ and heterogeneity $s$ asymptotically approach a stable plateau when $t \gg \tau$. This plateau simplifies the CPMC-derived inputs used to calculate $\Theta$, leading to an approximate steady-state expression for the probability of cell death.

***Derivation of Steady-State Malleability and Heterogeneity.*** At steady state, the malleability ratio $\gamma_k$ can be calculated via Eq. 27. If the dependence of $Se_{E,D_n}$ on $D_n$ is negligible - i.e., $Se(\overline{\varepsilon}_{i,a}, D_{n,b}) - Se(\overline{\varepsilon}_{i,a}, D_{n,a}) \ll Se(\overline{\varepsilon}_{2,a}, D_n) - Se(\overline{\varepsilon}_{1,a}, D_n)$ for all $D_{n,a} < D_n < D_{n,b}$ and $i = 1,2$ - the steady-state equation simplifies to:

$$\gamma_k \approx \frac{\beta_b}{\beta_a} \approx \left(\frac{D_{n,b}}{D_{n,a}}\right)^{Se(\overline{\varepsilon}_{1,a}\beta_a, D_{n,a}) - Se(\overline{\varepsilon}_{1,a}, D_{n,a})} \tag{49}$$

By combining Eqs. 10, 11, and 12, the sensitivity equation is approximated as $Se(\overline{\varepsilon}_{1,a}, D_{n,a}) \approx \frac{1}{D_{n,a}} \ln N_{\text{PD}} - G(\overline{\varepsilon}_{1,a}) F(D_{n,a})$, where $G(\overline{\varepsilon}) \approx \frac{1}{2}\sigma_{\phi_{\text{in}}}^2 \sqrt{\frac{\overline{\kappa}}{\overline{\varepsilon}}}$ and $F(D_n) = D_n \ln \frac{r_{\text{in}}}{r_{\text{min}}} + \frac{3-D_n}{D_n} \frac{r_{\text{min}}}{r_{\text{in}}} L^{1/D_n} \ln L$. Substituting this into Eq. 49 and assuming $\overline{\kappa}$ is the same for both $\overline{\varepsilon}_{1,a}$ and $\overline{\varepsilon}_{1,b}$, we derive:

$$\gamma_k \approx \left(\frac{D_{n,b}}{D_{n,a}}\right)^{\frac{1}{2}\sigma_{\phi_{\text{in},a}}^2 F(D_{n,a}) \sqrt{\frac{\overline{\kappa}}{\overline{\varepsilon}_{1,a}}} \left(1 - \frac{1}{\sqrt{\beta_a}}\right)} \tag{50}$$

Given that gene upregulation is typically high ($\beta_a \gg 1$), we can further simplify Eq. 50, as $\frac{1}{\sqrt{\beta_a}}$ becomes negligible compared to 1:

$$\gamma_k \approx \left(\frac{D_{n,b}}{D_{n,a}}\right)^{\frac{1}{2}\sigma^2_{\phi_{\text{in},a}} F(D_{n,a}) \sqrt{\frac{\overline{\kappa}}{\overline{\varepsilon}_{1,a}}}} \tag{51}$$

This approximation shows that under conditions of high upregulation, the malleability ratio is primarily driven by the geometric ratio of $D_n$ values. Biologically, this implies that for highly responsive genes, the variation in expression is governed more by chromatin structure differences than by initial expression levels.

At steady state, heterogeneity is determined by solving Eq. 28, yielding $s = COV[E_2] = \sqrt{2}G(\overline{\varepsilon}_2)$:

$$s_a \approx \frac{1}{\sqrt{2}}\sigma^2_{\phi_{\text{in},a}} \sqrt{\frac{\overline{\kappa}}{\beta_a \overline{\varepsilon}_{1,a}}} \tag{52}$$

$$s_b \approx \frac{1}{\sqrt{2}}\sigma^2_{\phi_{\text{in},b}} \sqrt{\frac{\overline{\kappa}}{\beta_a \overline{\varepsilon}_{1,a} \gamma_{\overline{\varepsilon}_2}}} \tag{53}$$

If $Se_{D_n}$ exhibits weak dependence on $D_n$, specifically when $Se(\overline{\varepsilon}_a, D_{n,b}) - Se(\overline{\varepsilon}_a, D_{n,a}) \ll D_{n,b} - D_{n,a}$, we approximate $\gamma_{\overline{\varepsilon}}$ as:

$$\gamma_{\overline{\varepsilon}_2} = \frac{\overline{\varepsilon}_{2,b}}{\overline{\varepsilon}_{2,a}} \approx N_{\text{PD}}^{\frac{1}{D_{n,b}} - \frac{1}{D_{n,a}}} \left(\frac{D_{n,b}}{D_{n,a}}\right)^{Se(\overline{\varepsilon}_{2,a}, D_{n,a})} \tag{54}$$

For genes with high upregulation ($\beta_a \gg 1$), the post-stress expression rate $\overline{\varepsilon}_{2,a}$ far exceeds the critical rate $\overline{\kappa}$ further simplifying $Se(\overline{\varepsilon}_{2,a}, D_{n,a})$. This leads to:

$$\gamma_{\overline{\varepsilon}_2} \approx N_{\text{PD}}^{\frac{1}{D_{n,b}} - \frac{1}{D_{n,a}}} \left(\frac{D_{n,b}}{D_{n,a}}\right)^{\frac{1}{D_{n,a}} \ln N_{\text{PD}}} \tag{55}$$

$$\gamma_h \approx N_{\text{PD}}^{\frac{1}{2}\left(\frac{1}{D_{n,a}} - \frac{1}{D_{n,b}}\right)} \left(\frac{D_{n,a}}{D_{n,b}}\right)^{\frac{1}{2D_{n,a}} \ln N_{\text{PD}}} \left(\frac{r_{\text{min}}}{r_{\text{in}}}\right)^{D_{n,a} - D_{n,b}} \tag{56}$$

$$h_b \approx \frac{3}{s_b\sqrt{\pi}} \approx 3\sqrt{\frac{2}{\pi}} \frac{1}{\sigma^2_{\phi_{\text{in},b}}} \sqrt{\frac{\overline{\varepsilon}_{1,a}\beta_a}{\overline{\kappa}}} N_{\text{PD}}^{\frac{1}{2}\left(\frac{1}{D_{n,b}} - \frac{1}{D_{n,a}}\right)} \left(\frac{D_{n,b}}{D_{n,a}}\right)^{\frac{1}{2D_{n,a}} \ln N_{\text{PD}}} \tag{57}$$

***Linear Approximations Using $D_n$.*** We can simplify the complex exponential terms by expressing them as linear functions of $\Delta D_n = D_{n,b} - D_{n,a}$. When $\Delta D_n \ll D_{n,a}$, the following approximations hold:

$$N_{\text{PD}}^{\frac{1}{2}\left(\frac{1}{D_{n,b}} - \frac{1}{D_{n,a}}\right)} \approx 1 + \frac{1}{2}\left(\frac{1}{D_{n,b}} - \frac{1}{D_{n,a}}\right) \ln N_{\text{PD}} \tag{58}$$

$$\approx 1 - \frac{\Delta D_n}{2D^2_{n,a}} \ln N_{\text{PD}} \tag{59}$$

$$\left(\frac{D_{n,b}}{D_{n,a}}\right)^{\frac{1}{2D_{n,a}} \ln N_{\text{PD}}} \approx 1 + \frac{\Delta D_n}{2D^2_{n,a}} \ln N_{\text{PD}} \tag{60}$$

where $\frac{1}{(2D_{n,a})} \ln N_{\text{PD}} \sim 2$. Using these approximations, we simplify Eqs. 56 and 57:

$$\gamma_h \approx \left(\frac{r_{\text{min}}}{r_{\text{in}}}\right)^{D_{n,a} - D_{n,b}} \left(1 + \frac{\Delta D_n}{D^2_{n,a}} \ln N_{\text{PD}}\right) \tag{61}$$

$$h_b \approx 3\sqrt{\frac{2}{\pi}} \frac{1}{\sigma^2_{\phi_{\text{in},b}}} \sqrt{\frac{\overline{\varepsilon}_{1,a}\beta_a}{\overline{\kappa}}} \tag{62}$$

Substituting 62 into 51 gives:

$$\gamma_k^{h_b} \approx \left(\frac{D_{n,b}}{D_{n,a}}\right)^{\frac{3}{2\pi} F(D_{n,a}) \left(\frac{r_{\text{min}}}{r_{\text{in}}}\right)^{D_{n,b} - D_{n,a}} \left(\sqrt{\beta_a} - 1\right)} \tag{63}$$

Here, $\frac{3}{2\pi}F(D_{n,a}) \approx 12.6$. We introduce the constants $Q = \frac{3}{2\pi}F(3) \approx \frac{9}{\sqrt{2\pi}}\log\left(\frac{r_{\text{in}}}{r_{\text{min}}}\right) \approx 12.6$ and $N_{\text{PD}} = \left(\frac{r_{\text{in}}}{r_{\text{min}}}\right)^{D_n}$, where $N_{\text{PD}} \sim 15$ kbp, representing the number of base pairs in a gene's interaction volume for $D_n = 2.5$ and $r_{\text{min}} = 1$ nm. From this, we derive the final approximations:

$$\tilde{\gamma}_h = N_{\text{PD}}^{\Delta D_n/D_{n,a}} \tag{64}$$

$$\tilde{\gamma}_k^{h_b} = \left(1+\frac{\Delta D_n}{D_{n,a}}\right)^{Q(\sqrt{\beta_a}-1)/N_{\text{PD}}^{\Delta D_n/D_{n,a}}} \tag{65}$$

Substituting these into Eq. 24, we obtain a closed-form expression for $\Theta$:

$$\tilde{\Theta}_b(\Theta_a) = \left(1+\left(1+\frac{\Delta D_n}{D_{n,a}}\right)^{Q(\sqrt{\beta_a}-1)/N_{\text{PD}}^{\Delta D_n/D_{n,a}}}\left(\frac{1}{\Theta_a}-1\right)^{\frac{1}{N_{\text{PD}}^{\Delta D_n/D_{n,a}}}}\right)^{-1} \tag{66}$$

This equation shows that $\Theta_b(\Theta_a)$ primarily depends on $D_{n,a}$, $\Delta D_n$, and $\beta_a$, linking chromatin structure changes, gene upregulation, and cell survival probability.

***Simplified Linear Expressions for Malleability and Heterogeneity.*** To further simplify the expressions for malleability and heterogeneity, we start with the interaction volume defined as $N_{\text{PD,a}} = \left(\frac{r_{\text{in}}}{r_{\text{min}}}\right)^{D_{n,a}}$. Using this definition, we derive:

$$\tilde{\gamma}_h \approx \frac{N_{\text{PD,a}}}{N_{\text{PD,b}}} \tag{67}$$

$$\tilde{\gamma}_k^{h_b} \approx \left(\frac{D_{n,b}}{D_{n,a}}\right)^{Q(\sqrt{\beta_a}-1)\gamma_h} \tag{68}$$

In a first-order approximation, we can express these terms in a more linearized form. When $\Delta D_n$ is small:

$$\tilde{\gamma}_h \approx 1-\Delta D_n \ln\left(\frac{r_{\text{in}}}{r_{\text{min}}}\right) = 1-\frac{\Delta D_n}{D_{n,a}}\ln N_{\text{PD}} \tag{69}$$

$$\tilde{\gamma}_k^{h_b} \approx 1+\left(\frac{\Delta D_n}{D_{n,a}}\right)Q(\sqrt{\beta_a}-1) \tag{70}$$

where $N_{\text{PD}} \equiv N_{\text{PD,a}}$. By assuming realistic values for $r_{\text{min}}$, we can further simplify the logarithmic terms. This leads to the following approximations:

$$\tilde{\gamma}_h \approx 1-3\Delta D_n \tag{71}$$

$$\tilde{\gamma}_k^{h_b} \approx 1+3\Delta D_n(\beta_a-1) \tag{72}$$

These approximations indicate that increasing the DNA base pairs within the transcriptional interaction volume enhances both the malleability and heterogeneity of gene expression. Specifically, the results show that transcriptional sensitivity to changes in chromatin packing is influenced by both gene upregulation ($\beta_a$) and structural alterations ($\Delta D_n$). The findings suggest that for highly responsive genes, variations in expression are driven more by differences in chromatin structure than by initial expression levels.

**Adaptive Inhibition Model for Tumor Growth.** While the equations derived in the previous sections for predicting the number of cells after a treatment are feasible to study experimentally using *in vitro* cultures, it is difficult to produce the same measure for *in vivo* studies. Therefore, we derived an equation to test the CDA model predictions directly in PDX experiments using the volume of a tumor as a proxy for cell number. We define the relative tumor volume ($RTV$) as $RTV(t) = V_{\text{treatment}}(t)/V_{\text{control}}(t)$ which is roughly equivalent to the probability of cell survival $P_s$. The CDA model predictions demonstrate that the efficacy of chemotherapy should decrease over time, as indicated by the decrease in $EIR$ (Fig. 12). This similarly results in a rapid decrease of the $RTV$ until it plateaus due to reduced efficacy. This indicates that the speed of acclimation of cancer cells to chemotherapy is in part due to the PDF($D_n$). Immediately after treatment (small $t$), the $EIR$ should decrease linearly due to the shift of the population average $D_n$. However, at longer time points (larger $t$), only the tail of the PDF($D_n$) is affected by the chemotherapy.

Building on this observation, we posit that tumor growth is controlled by the cell death induced by chemotherapy ($u$) and the rate of adaptation to the treatment ($a$). Additionally, within a tumor, there exists an population of low $D_n$ cells that cannot adapt to therapy and one with high $D_n$ cells that can. We define the relative growth rate of tumors, $V(t)$, through $\ln(V(t)/V(0)) = (p-c)t$, where $p$ represents the tumor growth rate in the absence of treatment and $c$ denotes the growth inhibition rate from chemotherapy. To account for adaptation, we introduce a cumulative adaptation term, $P_a(t)$, resulting in the revised equation $\ln(V(t)/V(0)) = (p-c)t + c\int_0^t P_a(t)dt$. The integral term $c\int_0^t P_a(t)dt$ captures the accumulation of the adaptable cell population over time; however, this term does not imply active proliferation among adaptable cells. Instead, it reflects the ability of this population to mitigate the overall inhibitory effects of chemotherapy, thus enhancing tumor survival through time-dependent adaptation. We anticipate that the adaptation term $P_a(t)$ will increase until it plateaus at a maximum value, $P_a^{\max}$, influenced by treatment strength. By assuming $P_a(t) = P_a^{\max}(1-\exp(-at))$ and defining the unadaptable inhibition rate as $u = c(1-P_a^{\max})$, we derive the final adaptive inhibition model:

$$\ln\frac{V(t)}{V(0)} = (p-u)t - \frac{c-u}{a}(1-\exp(-at)) \quad \mathbf{(73)}$$

where $V(t)$ is the tumor volume at time $t$, $p$ is the proliferation rate, $u$ is the unadaptable inhibition rate, $c$ is the initial inhibition rate, and $a$ is the adaptation rate. The parameters $p$ and $u$ are specific to the cell line and chemotherapy, while $c$ and $a$ depend on the treatment modality and the evolution of PDF($D_n$) over time. The parameters $u$, $c$, and $a$ were estimated from the data, while a fixed proliferation rate of $p = 1\times10^{-10}$ was used in all analyses. This dependence on PDF($D_n$) encapsulates both the adaptive capacity of cells undergoing treatment and the baseline inhibition imposed on unadaptable cells.

***TPR and Inhibition Index Derivation.*** As it is difficult to simultaneously evaluate many potential drug candidates to determine which one has the most significant impact on chromatin and, subsequently, cell death, we sought to derive a quantifiable value to characterize the magnitude of change. To find this, we use Eq. 73, which tells us the change in cancer cell number as a function of multiple inhibition and adaptation rates. Simply plotting the change in $D_n$ against the the cell number $N(t)$ would depict a general trend but not a clear relationship. Therefore, we wanted to determine a value that is a function of $N$ and $D_n$ so that we can better visualize the relationship between the two. However, in order to do that, we need to know all of the rates that are input into Eq. 73. For simplification, we derived an equation to eliminate several terms such that we could use values from the CDA model to predict the trend. Since rates $p$ and $u$ are specific to a particular cell line and chemotherapy, we can compare the effects of two potential TPRs to eliminate the need to know these rates. The rates $c$ and $a$ depend on the PDF($D_n$) and the TPR. To eliminate $p$, we consider three groups of cells (in the same cell line): one treated with a chemotherapy (high $D_n$), and the others treated with two different candidate TPRs (low $D_n$).
Here, we consider two groups of cells: one treated with TPR $A$ and another treated with TPR $B$, both on the same cell line. For $a_At \gg 1$ and $a_Bt \gg 1$, Eq. 73 becomes:

$$\ln\frac{N_A(t)/N_A(0)}{N_B(t)/N_B(0)} \approx \frac{c_B-u}{a_B} - \frac{c_A-u}{a_A} \quad \mathbf{(74)}$$

The rate $c$ can be approximated as $c \approx \Theta/\tau$.
We can use the derived approximate equations for $\tilde{\gamma}_h$ and $\tilde{\gamma}_k^{h_b}$ to simplify Eq. 24. First, from Eq. 64, $\tilde{\gamma}_h$ can be approximated using $\tilde{\gamma}_h \approx 1-(\Delta D_n/D_n)\ln(N_{\mathrm{PD}})$. Then, from Eq. 65, $\tilde{\gamma}_k^{h_b}$ can be simplified to $\tilde{\gamma}_k^{h_b} \approx 1+(\Delta D_n/D_n)Q(\sqrt{\beta_a}-1)$. By plugging these approximations into Eq. 24 and defining $\gamma = \ln(N_{\mathrm{PD}}) + Q(\sqrt{\beta_a}-1)$ we find:

$$\Theta_B = \Theta_A\left(1-\gamma\frac{D_{n,B}-D_{n,A}}{D_{n,A}}\right) \quad \mathbf{(75)}$$

Here, $\gamma \approx 10$ for physiologically relevant $N_{\mathrm{PD}}$ and $\beta_a$. Therefore, a $\Delta D_n \approx 0.1$ can lead to large changes in $\Theta$.
From $c \approx \Theta/\tau$, we find that $c_B \approx c_A\left(1-\gamma\frac{D_{n,B}-D_{n,A}}{D_{n,A}}\right)$ and:

$$c_B - c_A \approx -c_A\gamma\frac{D_{n,B}-D_{n,A}}{D_{n,A}} \quad \mathbf{(76)}$$

$$c_B - u \approx -u\gamma\frac{D_{n,B}-D_{n,u}}{D_{n,u}} \quad \mathbf{(77)}$$

where $D_{n,u}$ is chosen to correspond to $u$. Similarly, rate $a$ can be expanded as $a_A \approx a_B\left(1-\gamma\frac{D_{n,B}-D_{n,A}}{D_{n,B}}\right)$. This simplifies Eq. 74 to $\ln(N_A/N_B) \approx (\gamma/a_B)u(D_{n,A}-D_{n,B})$.

### Supplementary Materials and Methods

**Cell Culture and Treatments.** We obtained leiomyosarcoma (MES-SA, #CRL-1976; MES-SA.MX2, #CRL-2274), breast (MDA-MB-231, #HTB-26), colon (HCT-116, #CCL-247; HT-29, #HTB-38), and mouse embryonic fibroblast (MEF) cell lines from ATCC. Cells were maintained in their respective media as per ATCC protocols, supplemented with 10% FBS (#10-082-147, ThermoFisher Scientific). Ovarian cell lines (A2780, A2780.M248, A2780.M273, A2780.M175, and Ovcar8) were provided by Dr. Chia-Peng Huang Yang, originally sourced from Dr. Elizabeth de Vries's lab at Albert Einstein College of Medicine (12). All cell lines were tested for mycoplasma contamination using Hoechst 33342 (#H3570, ThermoFisher Scientific) within the past year. Experiments utilized cells from passages 5 to 20.

Human mesenchymal stem cells (hMSCs, #PCS-500-012, ATCC) were cultured in Dulbecco's Modified Eagle Medium (DMEM) with 4.5 g/L glucose (#11965092, ThermoFisher Scientific), supplemented with 10% FBS and 5 mL of 10$\times$ penicillin-streptomycin (#151400-122, ThermoFisher Scientific). For differentiation studies, we seeded hMSCs at $1.5 \times 10^4$ cells/mL in 24-well glass-bottom plates (#P24-1.5H-N, Cellvis). After 2 days, we transitioned to hMSC Osteogenic Differentiation Medium containing $\beta$-glycerophosphate, ascorbate, and dexamethasone (#PT-3002, Lonza). Media were changed every other day, and cells were imaged on day 4 post-induction.

Cells were cultured in 35 mm glass-bottom Petri dishes (#D35-14-1.5-N, Cellvis), allowing at least 24 hours for re-adherence after trypsinization prior to pharmacological treatment. All chemotherapeutic and potential TPR agents were purchased from Sigma Aldrich, unless otherwise noted. We treated cells with chemotherapy agents when approximately 30% confluent for at least 48 hours prior to imaging: paclitaxel (48 hours, 5 nM), oxaliplatin (48 hours, 15 µM), 5-fluorouracil (72 hours, 500 nM), docetaxel (48 hours, 5 nM), or gemcitabine (48 hours, 50 nM). For putative TPR treatments, cells were treated for 30 minutes prior to imaging with the following compounds at specified concentrations: celecoxib (75 µM, #SML3031), valproic acid (100 µM), aspirin (1 mM), digoxin (100 nM, #D6003), UNC0638 (1 µM), UNC1999 (1 µM), EGCG (25 nM), ginsenoside RB2 (1 µM), curcumin (25 µM), 4-phenylbutyrate (100 mM), simvastatin (1 µM), mevastatin (1 µM), resveratrol (1 µM), valinomycin (1 µM, #V3639).

Each treated cell population measured by PWS microscopy was compared to an untreated control population of the same cell type, plated on the same day with identical seeding density. All cells were maintained and imaged under physiological conditions (5% $CO_2$ and 37°C) throughout the experiment.

**Partial Wave Spectroscopic (PWS) Microscopy.** We performed Partial Wave Spectroscopic (PWS) microscopy using a commercial inverted microscope (Leica DMIRB) equipped with a Hamamatsu Image-EM CCD camera (C9100-13) and a liquid crystal tunable filter (LCTF; CRi). Monochromatic spectrally resolved images were acquired from 500 to 700 nm at 1 nm intervals, with illumination provided by an Xcite-120 LED Lamp (Excelitas). The PWS microscope setup used in this study is described in detail by Almassalha et al. (13).

The Hamamatsu Image-EM CCD camera has a pixel size of 16 µm. Using a 60x objective lens, the pixel size at the sample plane was approximately 267 nm, closely matching the theoretical resolution limit of the microscope, calculated as approximately 261 nm using the Rayleigh criterion with a numerical aperture (NA) of 1.4 and an average illumination wavelength of 600 nm. Spectrally resolved images were normalized by the incident light reflectance from the glass-media interface, using an independent reference from a cell-free field of view. A low-pass Butterworth filter was applied to reduce spectral noise prior to calculating the standard deviation of interference spectra ($\Sigma$) at each pixel (14). The resulting signal, which is related to chromatin packing at the nanoscale, was analyzed to extract the chromatin scaling exponent ($D_n$), which is a measure of chromatin organization derived from the chromatin density autocorrelation function.

The chromatin packing scaling exponent $D_n$ is calculated from the PWS signal and reflects nanoscale variations in chromatin density. The $D_n$ values were calculated from PWS spectral data using custom MATLAB scripts, as previously described (3). This measurement correlates with the average scaling of the autocorrelation function ($D_{\text{ACF}}$), which describes how chromatin density correlations decay with increasing separation. Specifically, $D_n$ is related to $D_{\text{ACF}}$ through the equation:

$$D_n \approx D_{\text{ACF}} + \frac{\ln(\text{VF})}{b},$$

where VF is the chromatin volume fraction, and $b$ is a constant dependent on optical and sample properties. This relationship illustrates how $D_n$ increases logarithmically with VF, capturing chromatin density variations and their impact on gene regulation. Higher $D_{\text{ACF}}$ values suggest a slower decay of density correlations, implying greater chromatin similarity over larger distances. These scripts quantify the standard deviation of interference spectra ($\Sigma$) at each pixel and relate these variations to the mass-density spatial autocorrelation function. By fitting the measured $\Sigma^2$ spectra to a theoretical model, we extracted $D_n$ values for each pixel.

This approach produces a pixel-wise chromatin packing scaling ($D_{\text{pixel}}$) map for each cell. The $D_{\text{pixel}}$ map allows us to assess chromatin packing domain variations across the nucleus, where higher $D_n$ values correspond to denser chromatin regions. To determine $D_n$ for individual cells, we averaged $D_{\text{pixel}}$ across cell nuclei. Typically, 100 to 200 cells were analyzed per condition to ensure robust statistical comparisons.

Pseudo-colored live cell PWS images were generated using Python, mapping $D_n$ values to a red colormap. $D_n$ values ranging from 2 to 3 were visually represented, with higher values indicated by warmer colors, providing a spatial map of chromatin packing domain variations across the nucleus.

**Cell Viability and Confluence Measurements.** We performed cell viability assays using fluorescence measurements with a BioTek Synergy Neo2 Reader at the Northwestern University HTAL Core. HCT116 cells were plated on 96-well flat, clear-bottom plates at a seeding density of 1,500 cells per well. After 48 hours for adherence, we treated the cells with oxaliplatin (15 µM) and evaluated them at 0, 2, 6, 12, 24, and 48-hour time points using the ApoTox-Glo triplex assay (#G6320, Promega). We added 20 µL of the assay viability reagent to each well, incubated at 37°Cfor 30 minutes, and measured fluorescence intensity (EX 400/20, EM 505/20, gain 87).

To calculate drug inhibition rates, we measured cell confluence by assessing cell density on dishes using transmission microscopy, quantifying the relative inhibition of cells in response to chemotherapeutic agents alone or in combination with potential TPR agents. We measured cell plate density for an area spanning 600,000 to 2,500,000 µm$^2$ using either a 40× or 20× air objective prior to PWS microscopy measurements. These measurements were obtained for three independent dishes for each condition group. We quantified cell density from transmission microscopy images using ImageJ (version 1.53c), which employs automatic thresholding and particle analysis to determine the area occupied by cells.

We calculated the inhibition rate ($IR$) for each treatment condition using the following equation:

$$IR = \frac{1}{t_{n+1} - t_n} \cdot \ln\left(\frac{C(t_{n+1})}{C(t_n)}\right) \tag{78}$$

where $t_n$ represents the time in days.

For measurements looking at the amount of cell death, we calculated normalized inhibition ($I_{\text{norm}}$) as the ratio of confluence in treated groups to that in control groups:

$$I_{\text{norm}} = \frac{C_{\text{treatment}}}{C_{\text{control}}} \tag{79}$$

**Calculation of $\Theta$ from Predicted $D_n$ Distributions.** Given that $1 - \Theta = \mathrm{e}^{-IR(\Delta t)\cdot\Delta t}$, we can use data from cell cluster tracking experiments to determine the probability of cell death as long as we have the initial distribution of $D_n$ prior to the addition of chemotherapy. To find experimental $\Theta$ values from PWS microscopy data on PDFs of $D_n$ at various timepoints, we employed an iterative solution to solve Eq. 39, as the integral-differential equation is difficult to solve analytically:

$$\begin{cases} N(D_n, \Delta t) & \approx 2^{\Delta t/\tau_2} \int N(D_n', 0) \cdot f(D_n - D_n')\, dD_n'\ \mathrm{e}^{-IR(D_n,\Delta t)\cdot\Delta t} \ldots \\ N(D_n, n_t\Delta t) & \approx 2^{n_t\Delta t/\tau_2} \int N(D_n', (n_t-1)\Delta t) \cdot f(D_n - D_n')\, dD_n'\ \mathrm{e}^{-IR(n_t\Delta t)\cdot n_t\Delta t} \end{cases} \tag{80}$$

where $\Delta t$ is any small interval (i.e., one day) and $n_t$ is the number of time intervals. We analyzed confluence data for HCT116 cells to calculate the mean and standard deviation of confluence across experimental groups. The overall inhibition rate for all $D_n$ at a single time point, $IR(n_t\Delta t)$, is calculated using the ratio of confluence in treated groups to that in control groups at time $t$. The drift due to cell division $f(D_n - D_n')$ was determined to be normally distributed around a mean value of 0 with a standard deviation of 0.685 using untreated control cluster tracking experiments as described in the section "Estimation of Cell Division Induced Drift in Population $D_n$". When $\Delta t \to 0$ and $n_t \to \inf$, the iterative solution should converge to the exact solution.

The number of cells as a function of $D_n$ and time, $N(D_n, \Delta t)$, is essentially the PDF($D_n$) at a specific time point. Therefore, to solve Eq. 80, we used the PDF at $t = 0$ from the experiment to produce an estimate of what the PDF would be at 48 hours using an initial guess of the $\Theta(D_n)$. PDFs of $D_n$ values were generated from experimental data using histogram methods with a bin width of 0.1. For each doubling time ($\tau_2$), we convolved the current PDF with the PDF incorporating drift in $D_n$ due to one division ($f(D_n - D_n')$):

$$\text{PDF}(D_{n,2}) = \text{PDF}(D_{n,1}) * (\sqrt{n_{\text{div}}} \cdot \sigma_{\delta D_n}) \tag{81}$$

where PDF($D_{n,1}$) is the PDF at the first time point, $n_{\text{div}}$ is the number of divisions, and $\sigma_{\delta D_n} = 0.0685$ is the standard deviation of the division-induced drift. The number of divisions over 48 hours was calculated as $n_{\text{div}} = 48/\tau_2 = 2.71$ cell divisions. We estimated $\tau_2$ using the total number of cells across all clusters at 0 and 24 hours with $\tau_2 = 24/\log_2[N(24h)/N(0h)] = 17.71$ hours. The PDF that resulted from the convolution was then multiplied by 2 to account for cell population doubling and normalized to depict a PDF distribution. Subsequently, we multiplied the PDF by the probability of cell survival (1-$\Theta$) for each $D_n$ value. To account for incomplete cell cycles, we calculated the residual doubling time (remainder of $48/\tau_2$) using an appropriate correction factor and expected cell population increase for the remaining time.

To find the experimental $\Theta$ in the cluster tracking PWS data of HCT116 cells treated with oxaliplatin, we used a two-step MSE minimization process. We first calculated an initial prediction for $\Theta$ for each $D_n$ value at the second time point as:

$$\Theta(D_n) = 1 - \left(\frac{\text{PDF}(D_{n,1})}{\text{PDF}(D_{n,2})} \cdot IR(t)\right)^{\tau_2/t} \tag{82}$$

The vector of $\Theta$ values was then used as the argument of the objective function for MSE minimization. We minimized the PDF shape-related error between the actual and predicted $D_n$ distributions at 48 hours, defined as:

$$\text{MSE}_{\text{shape}} = \sum_i \left(\text{PDF}_{\text{actual}}(D_i) - \text{PDF}_{\text{predicted}}(D_i)\right)^2 \tag{83}$$

We then refined the fit by minimizing a combined error that includes both shape and area under the curve, given by:

$$\text{MSE}_{\text{total}} = \text{MSE}_{\text{shape}} \cdot \left(\sum \text{PDF}_{\text{predicted}} - 1\right)^2 \tag{84}$$

where $\sum \text{PDF}_{\text{predicted}}$ is the sum of all values in the predicted PDF multiplied by the bin width used in the PDF calculation.

**Inhibition and TPR Index Calculation.** The relative inhibition between the control and drug-treated conditions was determined using the following formula:

$$\text{Inhibition Index} = \ln\left(\frac{1 - I_{\text{Chemo}}}{1 - I_{\text{TPR1}}}\right) / \ln\left(\frac{1 - I_{\text{Chemo}}}{1 - I_{\text{TPR2}}}\right) \tag{85}$$

where $I$ represents the mean inhibition due to the indicated treatment. This measure quantifies the effect of the drug treatment relative to the control, taking into account the inhibition levels in both conditions.
The TPR index, reflecting relative chromatin packing domain modulation between two drug treatments, was calculated across cell lines by comparing their normalized responses. To compare two drugs (denoted as *Drug1* and *Drug2*), the following formula was used to compute the $\Delta D_n$ Index:

$$\text{TPR Index} = \frac{D_{n,\text{TPR1}} + \sigma_{D_{n,\text{TPR1}}}}{D_{n,\text{Ctrl}} + \sigma_{D_{n,\text{Ctrl}}}} / \frac{D_{n,\text{TPR2}} + \sigma_{D_{n,\text{TPR2}}}}{D_{n,\text{Ctrl}} + \sigma_{D_{n,\text{Ctrl}}}} \tag{86}$$

where $D_n$ is the mean normalized value for each drug and $\sigma_{D_n}$ is the standard deviation of the PDF$(D_n)$. This index accounts for differences in chromatin packing domain modulation and treatment efficacy between two drugs, adjusted for population heterogeneity in response.

**Calculation of $\Theta$ from Population-Level Cell Confluence Data.** To estimate the cell death probability ($\Theta$) for chemo-treated cells to look at the influence of TPRs, we developed an approach based on cell confluence data. This method compares expected growth rates (without drug inhibition) to actual growth rates (with drug inhibition) following the logic of Eq. 37, ultimately providing the inhibition rate ($IR$) for each treatment. The confluence at some time interval $\Delta t$ is:

$$C_{n_t+1} \equiv C(\Delta t + n_t \Delta t) = C(n_t \Delta t) \exp\left[g(n_t \Delta t) - IR(n_t \Delta t)\right] \tag{87}$$

For our calculations, we use $\Delta t = 1$ day.
Growth rate was calculated by solving Eq. 87 for $g$ in an untreated control population such that $g(n_t) = \ln\left[C(n_t+1)/C(n_t)\right]$ for $\Delta t = 1$ day. We removed negative growth rates, implying cell death or loss, from subsequent analyses as such observations were not expected in the control conditions. We then used a linear regression to model the relationship between confluence and growth rate with $g = m \cdot C + b$, where $m$ is the slope and $b$ is the intercept. The regression yielded an $R^2 = 0.78$, indicating that it accounted for a significant portion of the variability in the data.
The IR during the early treatment phase (days 1 to 3) was determined for all conditions using two complementary methods. The primary method utilized confluence ratios (Eq. 78), quantified changes in confluence over time for all treatments. For comparative analysis, particularly when evaluating drug treatments against the control, the difference between expected and actual growth rates was used:

$$IR_i = g_{\text{expected},i} - g_{\text{actual},i} \tag{88}$$

where $g_{\text{expected},i}$ represents the expected growth rate (from the control model) and $g_{\text{actual},i}$ is the growth rate under drug treatment $i$. This was especially useful in dose-response analyses and for calculating the average IR for treatments like celecoxib.
We additionally calculated the overall growth rate using $g_2 = \ln(C_2/C_0)/2$ for $\Delta t = \tau_2 = 2$ days, where $C_2$ is the confluence on day 3 and $C_0$ is the confluence on day 1. An initial cell confluence $C_0$ of 33% was chosen as the baseline confluence for all conditions, as this was the average value determined by the cell confluence algorithm for day 0. The cell doubling time $\tau_2$ was set to 2 days. These constants were applied across the control, paclitaxel, and celecoxib treatment groups.
The mean growth suppression by celecoxib was calculated using data from treatment-specific time points. For celecoxib-treated samples, we calculated normalized growth inhibition using Eq. 79. Averaged values across treatment dates provided an estimate of celecoxib's efficacy in inhibiting cell proliferation. Growth rates for celecoxib-treated cells were adjusted based on the mean

growth suppression. The average IR for celecoxib was used to modify the growth rates, and a two-parameter exponential model was fitted to the data:

$$IR_{\text{Celecoxib}} = a \times e^{b \times \text{Confluence}} \tag{89}$$

where $a$ and $b$ are fitting parameters, capturing the non-linear relationship between celecoxib concentration and its inhibitory effect.

For cells treated with paclitaxel alone, $\Theta$ was calculated by fitting an exponential growth model to the observed data:

$$\Theta = 1 - e^{-IR \times t} \tag{90}$$

where $IR$ is the inhibition rate and $t$ is the treatment duration in days. $\Theta$ values on day 1 were calculated directly from the IR data. For combination treatments (TPR + chemotherapy), the effective cell death probability was calculated by subtracting the $IR$ values of the combination-treated group from the celecoxib-treated group:

$$\Theta_{\text{Combination-Celecoxib}} = 1 - e^{-(IR_{\text{Combination}} - IR_{\text{Celecoxib}})t} \tag{91}$$

where $IR_{\text{Combination}}$ is the inhibition rate of the combination treatment and $IR_{\text{Celecoxib}}$ is the inhibition rate of celecoxib alone. On day 2, for samples treated with both celecoxib and paclitaxel, inhibition was assessed by matching confluence values from the combination treatment group to the closest paclitaxel-only group. Adjustments were made for initial confluence and the duration cells had been plated prior to treatment.

Final $\Theta$ values were calculated using:

$$\Theta = 1 - (1 - \Theta)^{(\ln(2)/(b + m \times \text{Confluence}))} \tag{92}$$

where $b$ and $m$ are the intercept and slope from the control growth rate regression. This equation adjusts the cell death probability based on the growth rate's dependence on confluence. To ensure data quality, we applied a filter to exclude samples with standard deviation values greater than 0.2, ensuring only reliable data were used in the final analysis.

**Macromolecular Crowding Model.** We simulated mRNA synthesis dynamics under macromolecular crowding using the kinetic models described by Matsuda et al. (6) and Shim et al. (7). This approach quantifies how variations in transcription factor and RNA polymerase concentrations affect mRNA production in crowded environments. The simulations, conducted in Python, compute mRNA expression profiles across different crowding conditions, denoted as $\phi_{\text{in,model}}$, which represent the degree of macromolecular crowding. Matrix operations were handled using NumPy, while symbolic solutions were obtained with SymPy. Parameter values were initialized according to Matsuda et al. (6), unless otherwise specified. The model employed a fixed mRNA synthesis rate, $k_m$, and dissociation constants for transcription factors ($K_D^{\text{TF}}$) and RNA polymerase ($K_D^{\text{Pol II}}$). The optimal $k_m$ was determined iteratively by aligning the model's maximum mRNA production at $\phi_{\text{in,model}}$ with the experimentally observed $\phi_{\text{in,experiment}}$ (within a margin of error of 0.005).

We simulated mRNA synthesis across a range of transcription factor and RNA polymerase concentrations (denoted $TR$), spanning from $1 \times 10^{-7}$ to $1 \times 10^{-2}$ mM. mRNA production was modeled using differential equations that describe the binding and dissociation interactions between transcription factors, RNA polymerase, and DNA, while accounting for crowding effects. The dissociation rates for RNA polymerase and transcription factors were calculated based on total DNA concentration and their respective dissociation constants, adjusted according to the crowding factor $\phi_{\text{in,model}}$, which ranged from 0 to 0.505, reflecting physiological crowding conditions. Crowding effects on transcription kinetics were incorporated by modifying rate constants based on empirical coefficients derived from previous Brownian Dynamics simulations (6). The system of nonlinear equations governing transcription was solved symbolically to obtain mRNA synthesis rates across different crowding levels.

For each simulation condition, we computed the steady-state mRNA concentration, defined as the maximum of the mRNA expression curve, and plotted the relationship between $\phi_{\text{in,model}}$, transcriptional reactant ($TR$) concentrations, and mRNA output. To evaluate the dynamics of mRNA synthesis, we calculated the peak mRNA concentration and its corresponding $\phi_{\text{in,model}}$, as well as the second derivatives of the mRNA expression curves. The outputs that were then used to approximate $\bar{\varepsilon}$ included the maximum steady-state mRNA concentrations, the $\phi_{\text{in,model}}$ values at which they occurred, and the curvature of the mRNA response, assessed via second derivative analysis.

**Chromatin-Dependent Adaptability (CDA) Model Implementation and Optimization.** We implemented the CDA model using Python with NumPy and SciPy libraries. First, to investigate the impact of the key parameters on model predictions, we performed parameter scans. The key parameters scanned were $D_{n,0}$ (initial chromatin packing domain behavior), $\ln(E/\overline{E})$ (initial gene expression), $\beta_a = E_2/E_1$ (transcriptional amplification factor), $x_{\text{crit}}$ (critical upregulation threshold), and $T_{\text{crit}}$ (critical decision time). We employed a grid search approach, systematically varying these parameters within physiologically relevant ranges.

For comparisons between model predictions and experimental data, we employed the `scipy.optimize.minimize` function to minimize the MSE. We used the L-BFGS-B algorithm for minimization, allowing for bounded optimization of parameters within physiologically relevant ranges.

For the optimization for $\Theta$ vs. $D_n$, the objective function for minimization was:

$$f(\vec{p}) = \sum_{i=1}^{n} \left[\Theta_{\text{model}}(D_{n,i}, \vec{p}) - \Theta_{\text{exp}}(D_{n,i})\right]^2 \quad \textbf{(93)}$$

where $\vec{p}$ is the vector of model parameters, $D_{n,i}$ are the experimental $D_n$ values, and $\Theta_{\text{exp},i}$ are the experimental $\Theta$ values. The optimization was constrained with bounds of $[0,1]$ for all $\Theta$ values to ensure biological plausibility. To determine the optimal values for $\beta_a$, $x_{\text{crit}}$, and $\ln(E/\overline{E})$, we performed a grid search over physiologically relevant ranges: $\beta_a$ from 5 to 25, $x_{\text{crit}}$ from 2 to 12, and $\ln(E/\overline{E})$ from -3 to 3. For each parameter combination, we calculated the MSE between the model-predicted and experimental $\Theta$ vs. $D_n$ curves, selecting the parameter set yielding the lowest MSE as optimal. Optimization yielded parameters consistent with key survival genes exhibiting low initial expression ($\ln(E/\overline{E}) \approx -3$) and significant upregulation ($\beta_a \approx 19$) above a critical threshold ($x_{\text{crit}} \approx 8$). The $\beta_a/x_{\text{crit}}$ ratio primarily determined fit quality, with the final model achieving an MSE of 0.031 for HCT116 cells treated with oxaliplatin.

To determine the threshold value for assessing TPR candidate efficacy, we first calculated the mean $D_n$ value for the control group. We then employed an iterative adjustment approach to identify the threshold value using the $\Theta$ function. The initial parameters for the model were set as follows: $x_{\text{crit}} \approx 8$, $\beta_a \approx 18$, and $\ln(E/\overline{E}) \approx 0.05$. Initially, the $D_n$ threshold value ($D_{n,crit}$) was set to the mean $D_n$ value of the control group. Using the $\Theta$ function, we calculated $D_{n,crit}$ incorporating these model parameters. $D_{n,crit}$ was then decreased incrementally by 0.001 until $\Theta$ approached a value just below 0.999. The final $D_{n,crit}$ threshold value obtained was approximately 2.123, with a corresponding $\Theta$ value of 0.999.

For the optimization for $\Theta_b$ vs. $\Theta_a$, we used cell confluence data of A2780 cells treated with the combination of TPR (celecoxib) and chemotherapy (paclitaxel) rather than PWS microscopy data. We utilized the experimental data of cell death probability for paclitaxel alone ($\Theta_{\text{pac}}$) and the combination treatment ($\Theta_{\text{combo}}$) to fit the model parameters. The optimization function minimized the error between the predicted and experimental $\Theta_{\text{combo}}$ values:

$$f(\vec{p}) = \sum_{i=1}^{n} \left[\Theta_b(\Theta_{\text{pac},i}, \vec{p}) - \Theta_{\text{combo}}(\Theta_{\text{pac},i})\right]^2 \quad \textbf{(94)}$$

where $\Theta_b$ is the model-predicted cell death probability for the combination treatment, $\vec{p}$ are the input model parameters $D_{\text{control}}, D_{\text{TPR}}, \beta_a, \ln(E/\overline{E})$, and $D_{\text{control}}$ and $D_{\text{TPR}}$ are the initial $D_n$ values for control and TPR-treated cells, respectively. We employed the minimization with bounds of (3, 20) for $\beta_a$ and ($e^{-3}$, $e^{3}$) for $\ln(E/\overline{E})$. This optimization yielded $\beta_a \approx 6$ and $\ln(E/\overline{E}) \approx -3$ for the A2780 cells, with an MSE of 0.031. The fit was primarily dependent on $\beta_a$, with very little impact of $\ln(E/\overline{E})$. These optimized parameters were then used to generate both exact and approximate $\Theta_b$ curves for comparison with experimental data.

**Patient-Derived Xenograft (PDX) Tumor Models.** The following research protocol was approved by Northwestern University's institutional review board. High Grade Serous Ovarian Cancer (HGSOC) patient-derived tissue samples were obtained from chemotherapy-naive ovarian cancer patients following surgical resection at Prentice Women's Hospital of Northwestern University, from September 2013 to June 2014, with patient consent for tissue acquisition. For this study, we utilized a cryopreserved patient-derived tissue sample (OVCA10) at its fourth generation (passage 4). This PDX model was selected for its ability to closely mimic the heterogeneity and drug response characteristics of human ovarian cancer, offering a more clinically relevant system compared to traditional cell line-derived xenografts.

Tumor fragments measuring 2 × 2 mm were subcutaneously engrafted into the right flank of non-obese diabetic/severe combined immunodeficient (NOD/SCID) gamma (NSG) mice (Jackson Laboratory). Once the engrafted PDX tumors reached a volume of 150–200 $mm^3$, the mice were randomized into five experimental groups: celecoxib vehicle control, paclitaxel vehicle control, 25 mg/kg celecoxib, 1.7 mg/kg paclitaxel, and a combination group (25 mg/kg celecoxib and 1.7 mg/kg paclitaxel).

Celecoxib (25 mg/kg) or vehicle was administered daily via oral gavage, while paclitaxel (1.7 mg/kg) or vehicle was administered twice weekly (Mondays and Thursdays) via intraperitoneal (IP) injection. The experiment spanned 4 weeks, with tumor size and body weight monitored biweekly. Tumor dimensions were measured using digital calipers, recording the longest diameter (length, $l$) and the diameter perpendicular to it (width, $w$). Tumor volume ($V$) was calculated using the formula $V = l \cdot w^2/2$. At the end of the 4-week treatment period, the mice were euthanized, and PDX tumors were collected and preserved in 10% formalin.

For fitting the adaptive inhibition model to experimental data, non-linear least squares optimization was performed using `scipy.optimize.curve_fit`.

**Flow Cytometry and Cell Counting.** Flow cytometry was performed at the Northwestern University Flow Cytometry Core using a BD LSRII instrument, and data were analyzed using FlowCytometryTools 0.4.5, an open-source Python software package. Apoptotic induction was measured using CellEvent Caspase-3/7 Green Detection Reagent and Hoechst 33342 (#C10423 and #H3570; ThermoFisher Scientific, Waltham, MA). Cells were trypsinized, stained with 2 µM Caspase-3/7 and 4 µM

Hoechst 33342 for 30 minutes, then processed for flow cytometry. Following staining, cells were centrifuged for 5 minutes at 500×g, washed with PBS, and resuspended in 1 mL of fresh media. Mock-stained cells were collected under the same preparation conditions.

We collected data from 20,000 cells by forward and side scattering channels for each group, setting illumination intensities to minimize autofluorescence from unstained cells. Flow cytometry was conducted on the following groups of A2780 cells: unstained control cells, stained control cells, stained 48-hour paclitaxel mono-treated cells, stained 48-hour putative TPR-treated cells (celecoxib, and digoxin), and stained 48-hour co-treated paclitaxel + putative TPR cells. Gates were established for Caspase-3/7 staining and Hoechst 33342 to minimize false positives from unstained cells (less than 0.1% of total). The percentage of apoptotic cells was assessed as the ratio of Caspase-3/7 positive cells to the population of Hoechst 33342 positive cells. Error bars represent uncertainty based on a ±10% change in gating thresholds.

Cell counts were measured using an automatic cell counter (Countess II FL Automated Cell Counter). Cells were trypsinized, combined with floating cells from the same population, and stained with either 1 µM Image-iT DEAD Green viability stain, 4 µM Hoechst 33342, or 2 µM YO-PRO-1 Iodide (#I30201, #H3570, and #Y3603; ThermoFisher Scientific, Waltham, MA). After staining, cells were centrifuged for 5 minutes at 1000 rpm, washed with PBS, and resuspended in 1 mL of fresh media. Automated cell counting was performed on various groups of A2780 cells under different treatment conditions: control cells, 48-hour celecoxib-treated cells, 48-hour paclitaxel-treated cells, and 48-hour celecoxib and paclitaxel co-treated cells.

**Chromatin Scanning Transmission Electron Microscopy (ChromSTEM).** Chromatin Scanning Transmission Electron Microscopy (ChromSTEM) was employed to assess the characteristics of packing domains in HCT116 cells, adhering to established protocols (2). The key parameters evaluated included the packing domain scaling exponent ($D_{\text{PD}}$), the average packing domain chromatin volume concentration ($\text{CVC}_{\text{PD}}$), the average packing domain volume packing efficiency ($A_v$), and the average genomic size of a packing domain ($N_{\text{PD}}$).

For electron microscopy preparation, HCT116 cells underwent standard fixation and staining procedures. Briefly, cells were fixed in 2.5% glutaraldehyde in 0.1 M sodium cacodylate buffer, post-fixed with 1% osmium tetroxide, dehydrated through a graded ethanol series, and embedded in Epon resin. Ultrathin sections (70–90 nm) were cut using a diamond knife on an ultramicrotome and subsequently mounted on copper grids. The sections were stained with uranyl acetate and lead citrate prior to imaging. ChromSTEM imaging was performed using a Thermo Fisher Talos F200X G2 transmission electron microscope operating at an accelerating voltage of 200 kV in STEM mode. Images were acquired with a pixel size of 1 nm and a dwell time of 20 µs. The chromatin packing domain parameters, including $D_{\text{PD}}$, $r_{\text{PD}}$, $r_{\text{fiber}}$, $\text{CVC}_{\text{PD}}$, and $A_v$, were quantified using custom MATLAB scripts (2).

The genomic size of a packing domain, $N_{\text{PD}}$, was calculated through a computational approach implemented in Python. This process involved determining the number of voxels within the packing domain and multiplying by the amount of chromatin contained within a single voxel, expressed in base pairs (bp). The genomic size of chromatin within individual ChromSTEM voxels was estimated based on an assumed DNA density and voxel volume. The DNA density ($\rho_{\text{DNA}}$) was assumed to be 2 g/cm$^3$, corresponding to unhydrated DNA, which reflects the condition that the highest intensity observed in tomograms represents 100% unhydrated DNA.

The voxel mass, measured in base pairs ($N_{\text{voxel}}$), was computed using the following equation (1):

$$N_{\text{voxel}} = \frac{V_{\text{voxel}} \cdot \rho_{\text{DNA}}}{2 \cdot M_{\text{nucleotide}} \cdot (1.660 \times 10^{-24})} \tag{95}$$

Here, the voxel volume ($V_{\text{voxel}}$) was based on an assumed ChromSTEM voxel size ($l_{\text{voxel}}$) of 2 nm, converted to cm$^3$. The molecular weight of a nucleotide ($M_{\text{nucleotide}}$) was assumed to be 325 Daltons, and the atomic mass unit conversion factor was $1.660 \times 10^{-24}$ g/AMU.

To calculate $N_{\text{PD}}$, the genomic size of the packing domain, we applied the fractal scaling relationship (2):

$$N_{\text{PD}} = \left( A_v \cdot \frac{r_{\text{PD}}}{l_{\text{voxel}}} \right)^{D_{\text{PD}}} \cdot N_{\text{voxel}} \tag{96}$$

Sensitivity analyses were conducted using the 25$^{\text{th}}$, 50$^{\text{th}}$, and 75$^{\text{th}}$ percentiles for $D_{\text{PD}}$, $N_{\text{PD}}$, and $A_v$ to evaluate parameter variability. For additional analyses, mean values from the HCT116 cell population were employed: $D_{\text{PD}} = 2.62$, $\text{CVC}_{\text{PD}} = 0.275$, $A_{\text{v,PD}} = 0.61$, and $N_{\text{PD}} = 381$ kbp. Median values were used for $r_{\text{PD}} = 110$ nm and $r_{\text{fiber}} = 10$ nm.

**Statistical Analysis.** All statistical analyses were conducted using Python, with SciPy and pandas as the primary libraries for data processing. To assess statistical significance across multiple conditions, a custom function was developed to perform pairwise comparisons between a reference condition and all other conditions within each group. Welch's t-test was used to calculate P-values for these comparisons. This approach was applied in several contexts, including comparisons of nuclear $D_n$ values between treated and control cell populations, assessments of cell viability between different treatment conditions,

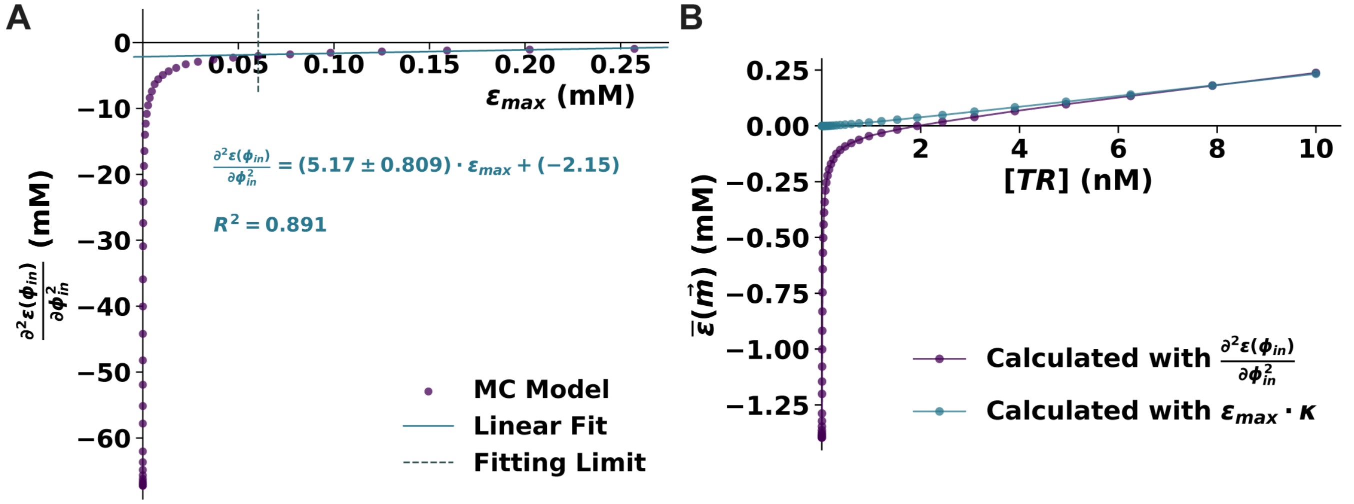


**Fig. 1.** Determination and validation of $\kappa$ for approximating the second derivative in the Taylor series expansion. (A) Plot of the second derivative $\left(\frac{\partial^2 \varepsilon(\phi_{\text{in}})}{\partial \phi_{\text{in}}^2}\right)$ as a function of the maximum expression ($\varepsilon_{\text{max}}$) for the input vector of molecular factors ($\vec{m}$). The dashed gray line indicates where the region for the linear fitting begins. (B) Comparison of $\bar{\varepsilon}$ calculated directly with the second derivative vs. the $\kappa$ approximation. For low concentrations of TR, the $\kappa$ approximation diverges from the direct calculation with the second derivative.

and analyses of tumor volumes in PDX models across treatment groups. We considered P-values less than 0.05 as statistically significant, and significance levels were denoted as follows: $^{*}P < 0.05$, $^{**}P < 0.01$, and $^{***}P < 0.001$.
In visualizing $D_n$ distributions, violin plots were used, with dashed lines indicating the $25^{\text{th}}$, $50^{\text{th}}$ (median), and $75^{\text{th}}$ percentiles. The width of each violin reflects the frequency distribution of $D_n$ values, providing a detailed representation of the data. Unless otherwise specified, error bars in figures represent the standard error of the mean (SEM). For error propagation in calculations, the uncertainties package was used to automatically propagate uncertainties through all relevant steps.

**Computational Resources and Code Availability.** All computational analyses were conducted using Python 3.7.1, with NumPy 1.15.4, SciPy 1.1.0, and pandas 0.23.4. Data visualization was performed using Matplotlib 3.3.2 and Seaborn 0.11.0.

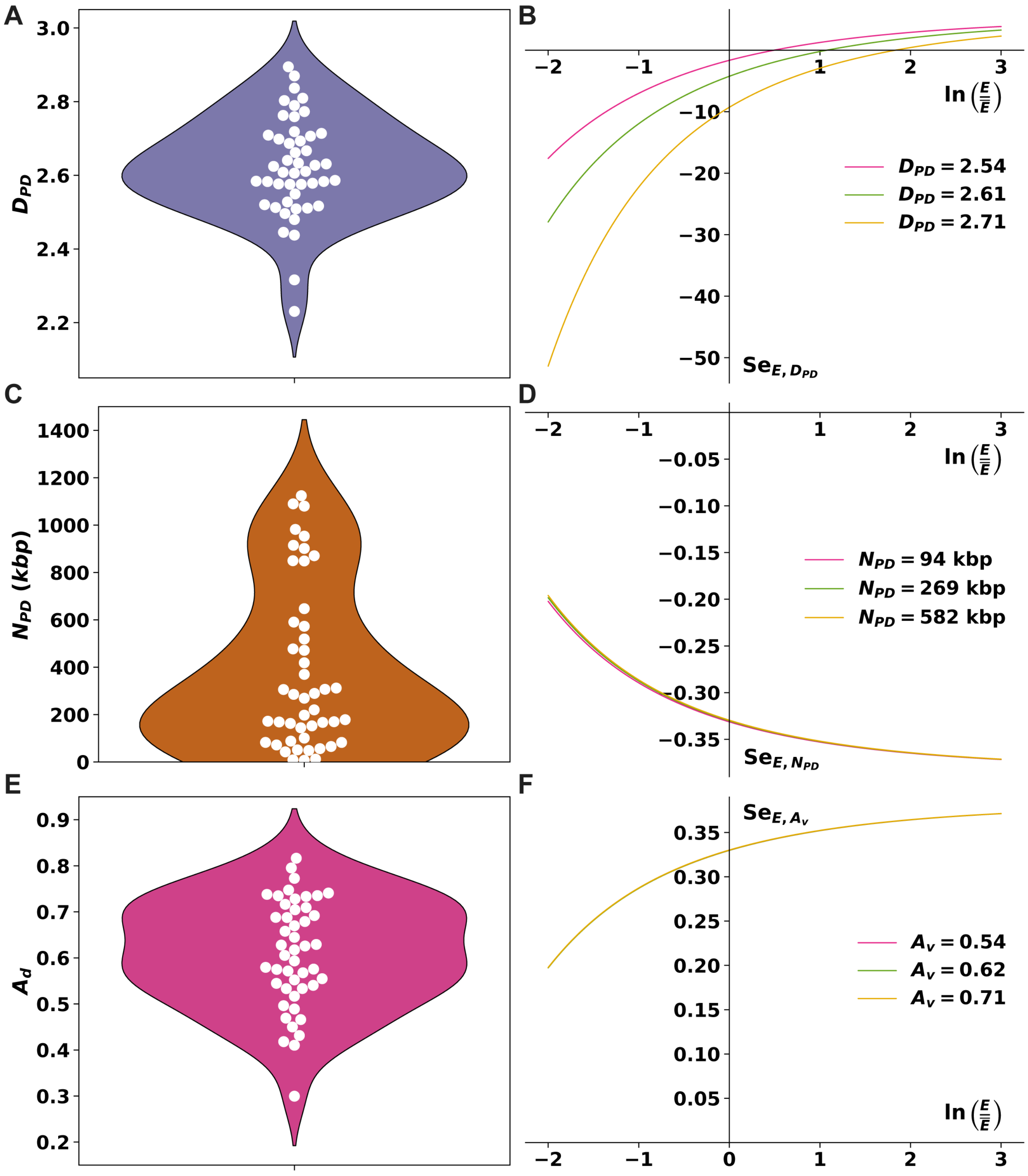


**Fig. 2.** CPMC parameters and their effects on gene expression sensitivity. (A) Distribution of scaling within packing domains ($D_{\text{PD}}$) in HCT116 cells. (B) Sensitivity of gene expression to changes in packing domain scaling ($Se_{E,D_{\text{PD}}}$) as a function of relative gene expression ($\ln(E/\overline{E})$) for different $D_{\text{PD}}$ values. (C) Distribution of packing domain sizes ($N_{\text{PD}}$) in HCT116 cells. (D) Sensitivity of gene expression to changes in packing domain size ($SeE, N_{\text{PD}}$) as a function of relative gene expression for different $N_{\text{PD}}$ values. (E) Distribution of packing domain packing efficiency factors ($A_v$) in HCT116 cells. (F) Sensitivity of gene expression to changes in average nuclear crowding ($Se_{E,A_v}$) as a function of relative gene expression for different $A_v$ values. Sensitivities were calculated using the CPMC model with parameters derived from ChromSTEM analysis of HCT116 cells. Note the different y-axis scales, indicating that $Se_{E,D_{\text{PD}}}$ has the largest magnitude among the three parameters.

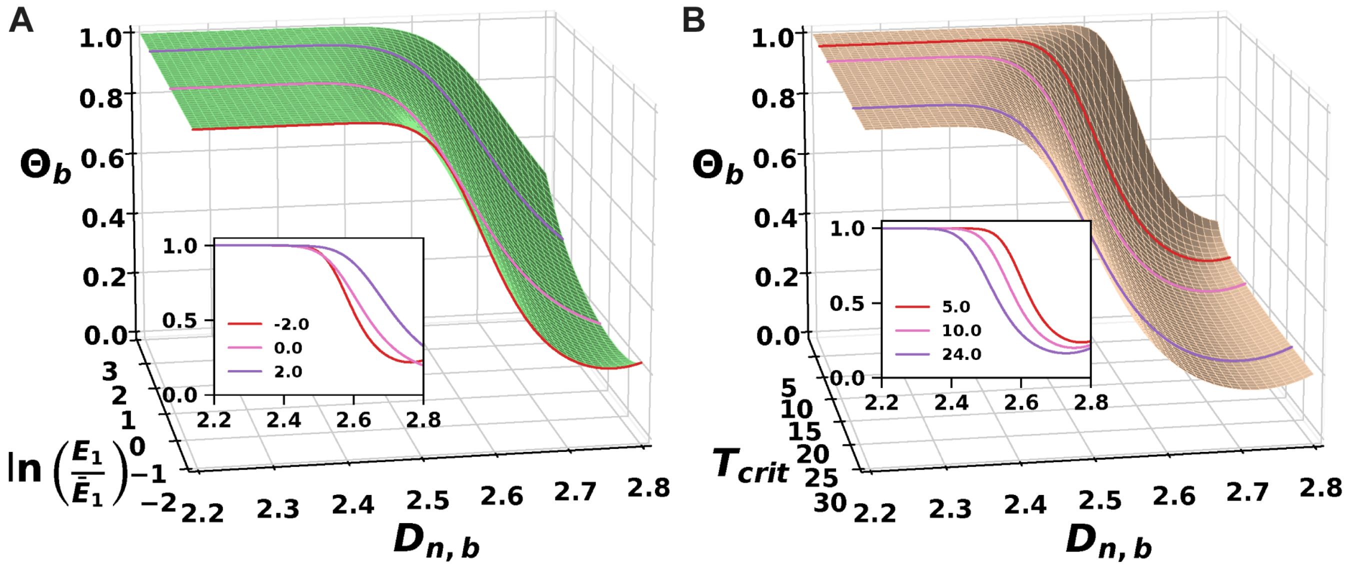


**Fig. 3.** Parameter scans for selected free parameters in the CDA model. (A) Cell death probability ($\Theta$) as a function of chromatin packing scaling ($D_{n,b}$) and relative initial gene expression ($\ln(E_1/\overline{E}_1)$). Inset shows individual curves for select $\ln(E_1/\overline{E}_1)$ values. Fixed parameters: $\beta_a = 10$, $x_{\text{crit}} = 5$, $T_{\text{crit}} = 7$ hours. (B) $\Theta$ as a function of $D_{n,b}$ and critical decision time ($T_{\text{crit}}$). Inset displays curves for select $T_{\text{crit}}$ values. Fixed parameters: $\ln(E_1/\overline{E}_1) = -2$, $\beta_a = 10$, $x_{\text{crit}} = 5$. All plots were generated with $D_{n,a} = 2.6$ and $D_{n,b}$ ranging between 2.2 and 2.8.

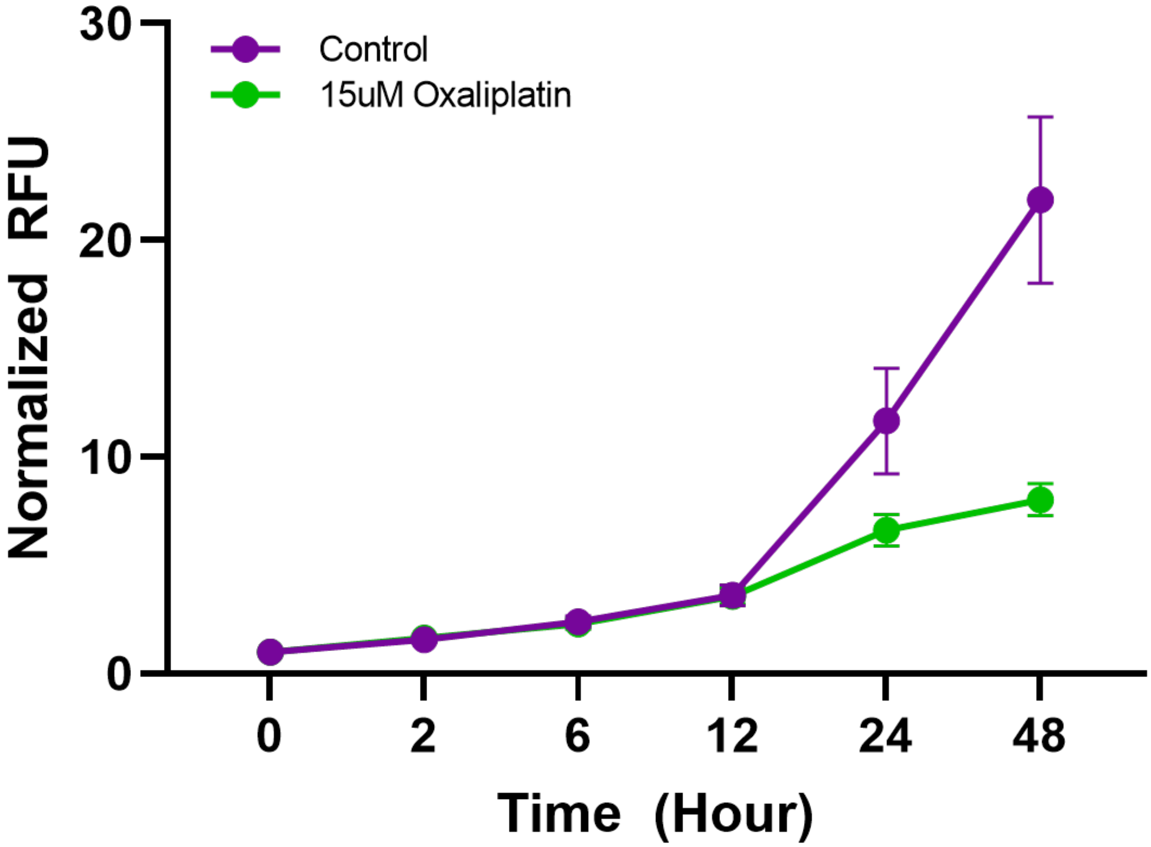


**Fig. 4.** Cell viability analysis of HCT116 cells treated with oxaliplatin. The line plot shows the time course of normalized relative fluorescence units (RFU) measuring cell viability in control (purple) and 15 µM oxaliplatin-treated (green) HCT116 cells over 48 hours. Fluorescence intensity was measured using a viability assay at 0, 2, 6, 12, 24, and 48 hours post-treatment. Error bars represent standard error of the mean from three independent experiments.

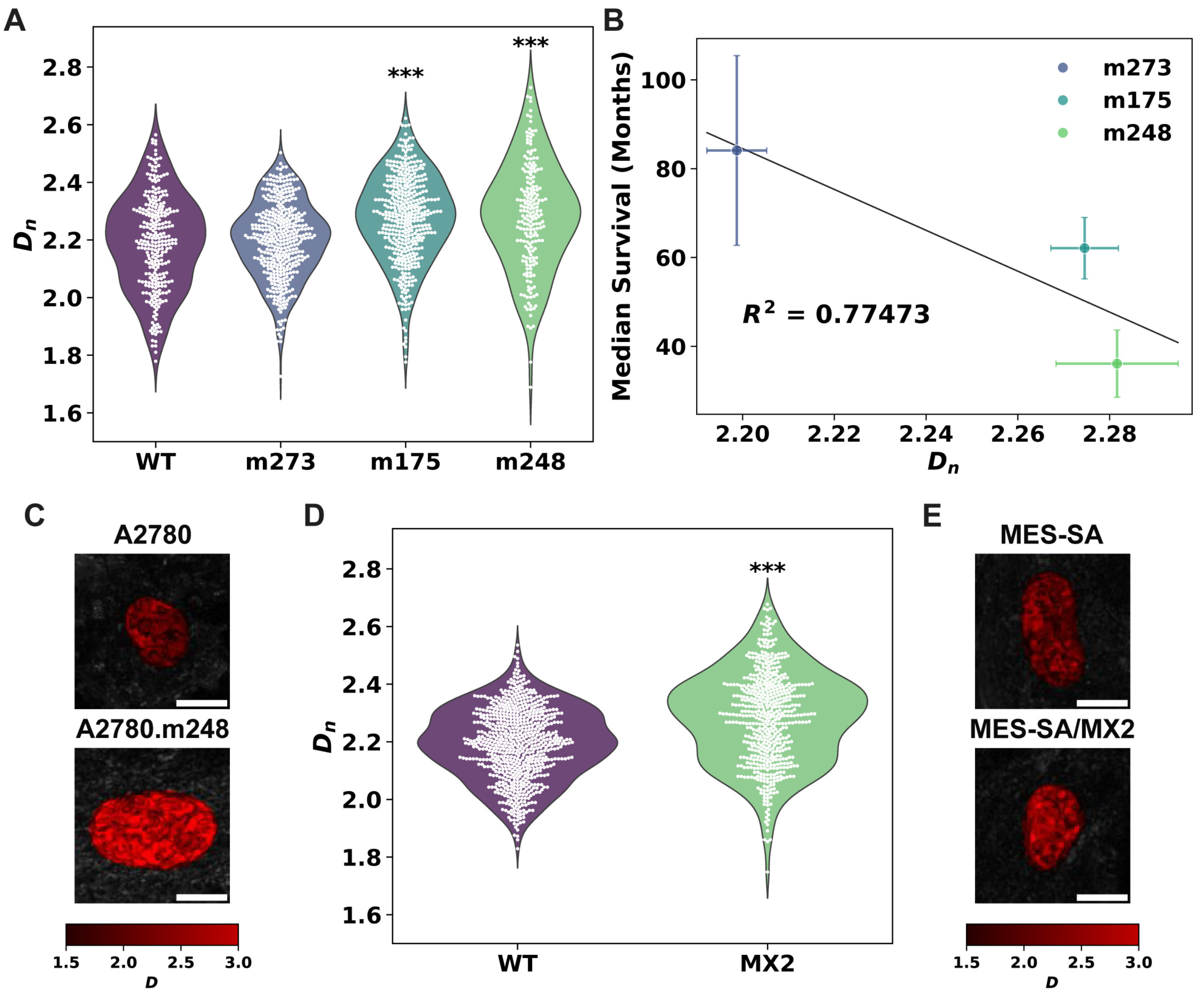


**Fig. 5.** Cancer cells with selective resistance to chemotherapy have higher $D_n$. (A) Violin plots showing $D_n$ distributions for A2780 wild-type (WT) and TP53 mutant subclones (m273, m175, m248) under normal growth conditions. Significant increases in $D_n$ are observed in m175 and m248 compared to WT ($P < 0.001$). (B) Correlation between median survival of high-grade serous epithelial ovarian carcinoma patients (TCGA data) and $D_n$ for cells with matching TP53 mutations. A strong negative correlation is observed ($R^2 = 0.77473$). (C) Representative PWS microscopy images of A2780 WT and TP53 mutant A2780.m248 cells. (D) Violin plots showing increased $D_n$ in chemoresistant MES-SA/MX2 subclone compared to chemosensitive MES-SA WT cells ($P < 0.001$). (E) Representative PWS images of MES-SA WT and MES-SA/MX2 cells. For A and D, dashed lines in violins represent $75^{\text{th}}$ percentile, median, and $25^{\text{th}}$ percentile. For C and E, pseudocolor represents $D_n$ values, with brighter red indicating higher $D_n$. Scale bars: 15 µm.

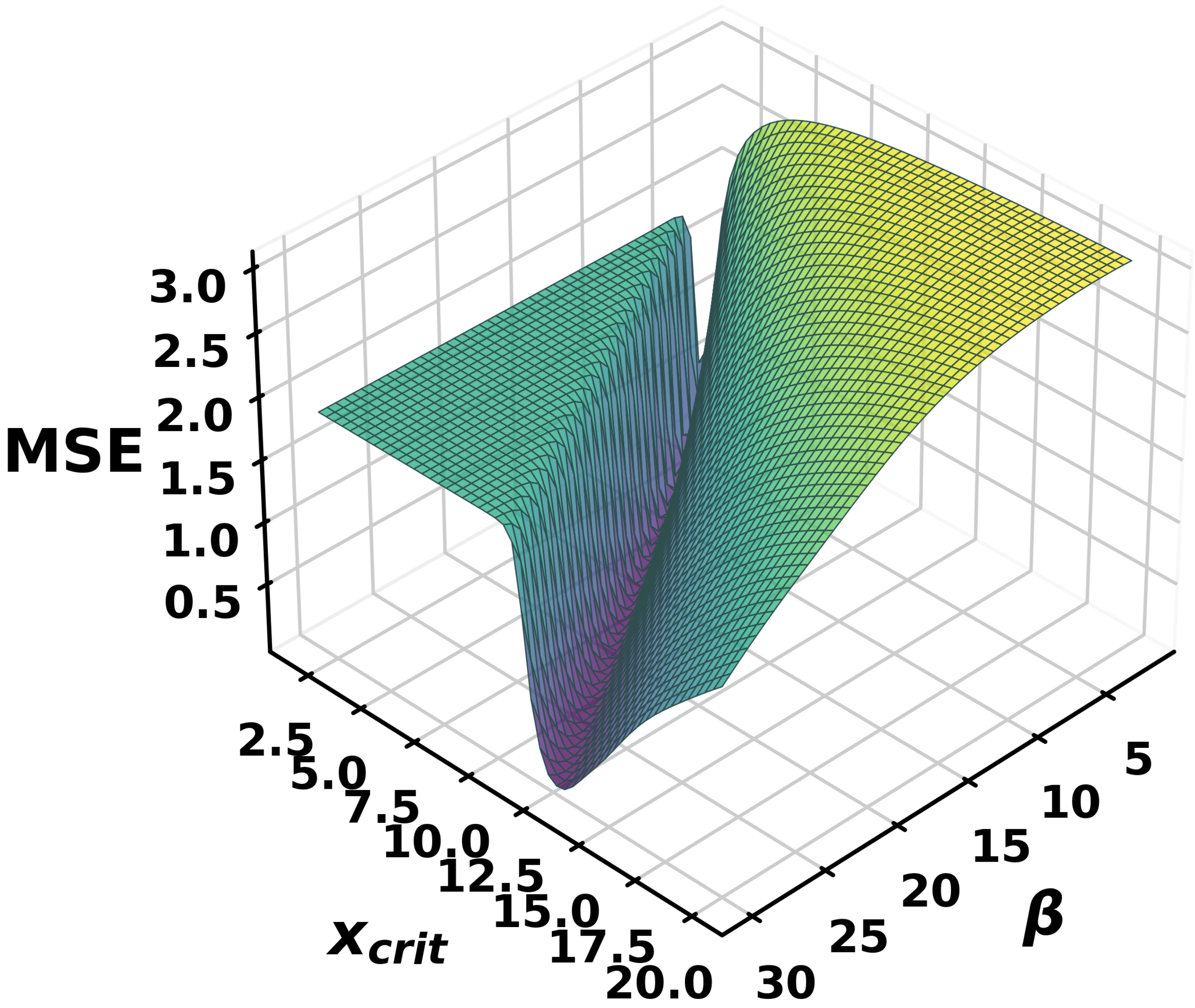


**Fig. 6.** Optimization of CDA model parameters for $\Theta(D_n)$. 3D plot showing mean squared error (MSE) of the CDA model fit as a function of $\beta_a$ and $x_{\text{crit}}$. The optimization was performed using experimental data from HCT116 cells treated with oxaliplatin. $\beta_a$ represents the gene upregulation factor, while $x_{\text{crit}}$ is the critical threshold for upregulation. The color gradient represents MSE values, with darker colors indicating lower MSE and better model fit. The plot demonstrates that the ratio between $\beta_a$ and $x_{\text{crit}}$ primarily determines the quality of fit, with optimal values consistent with key survival genes having significant upregulation ($\beta_a \approx 19$) above a critical threshold ($x_{\text{crit}} \approx 8$).

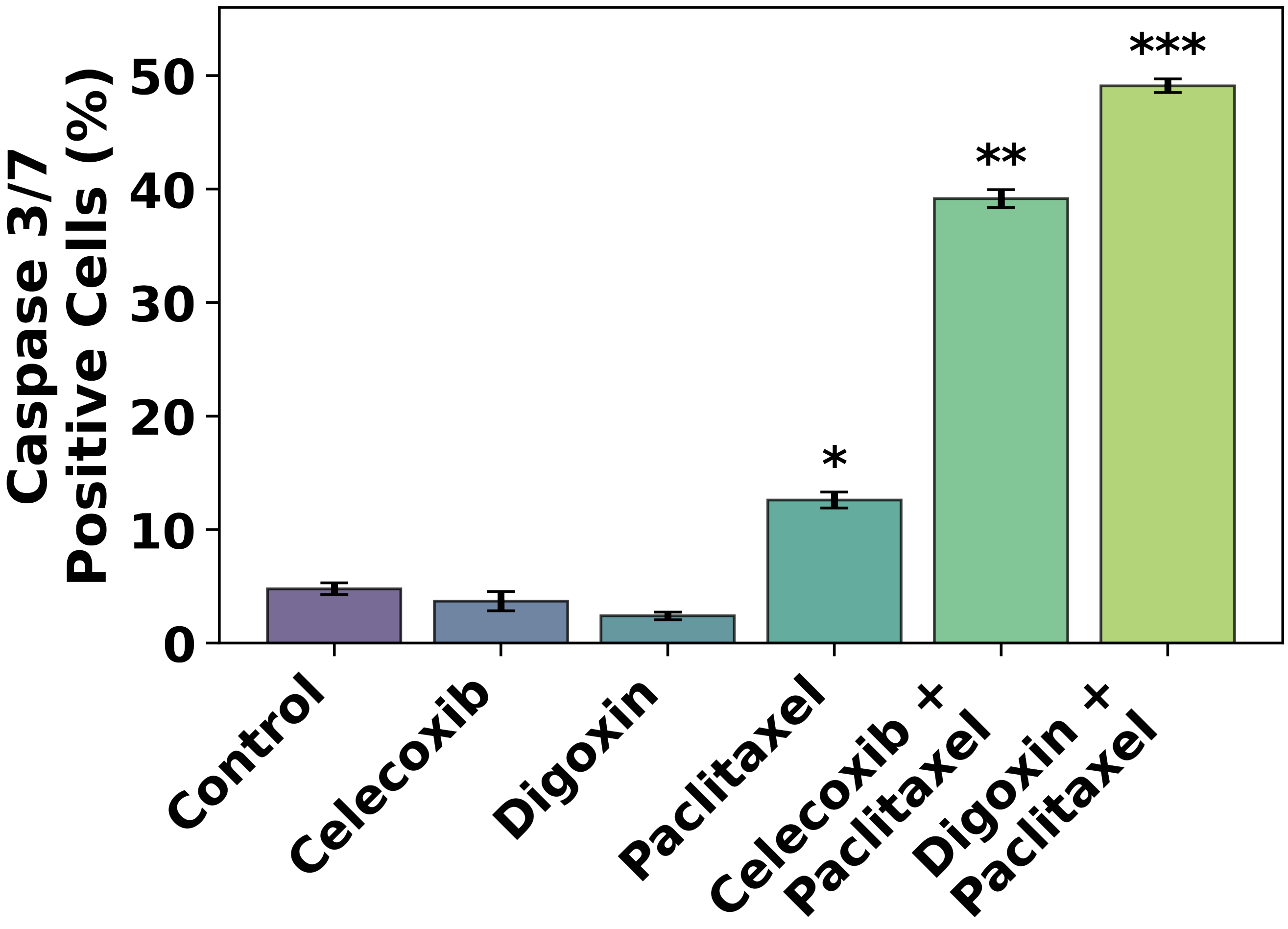


**Fig. 7.** TPRs enhance chemotherapy-induced apoptosis without inducing significant cell death on their own. Bar graph shows the percentage of Caspase 3/7 positive A2780 cells after 48 hours of treatment with control, celecoxib alone, digoxin alone, paclitaxel alone, and combinations of celecoxib or digoxin with paclitaxel. Error bars represent standard error of the mean from three independent experiments. Significance levels compared to paclitaxel alone: $^{*}P < 0.05$, $^{**}P < 0.01$, $^{***}P < 0.001$ (determined using unpaired t-test with unequal variance).

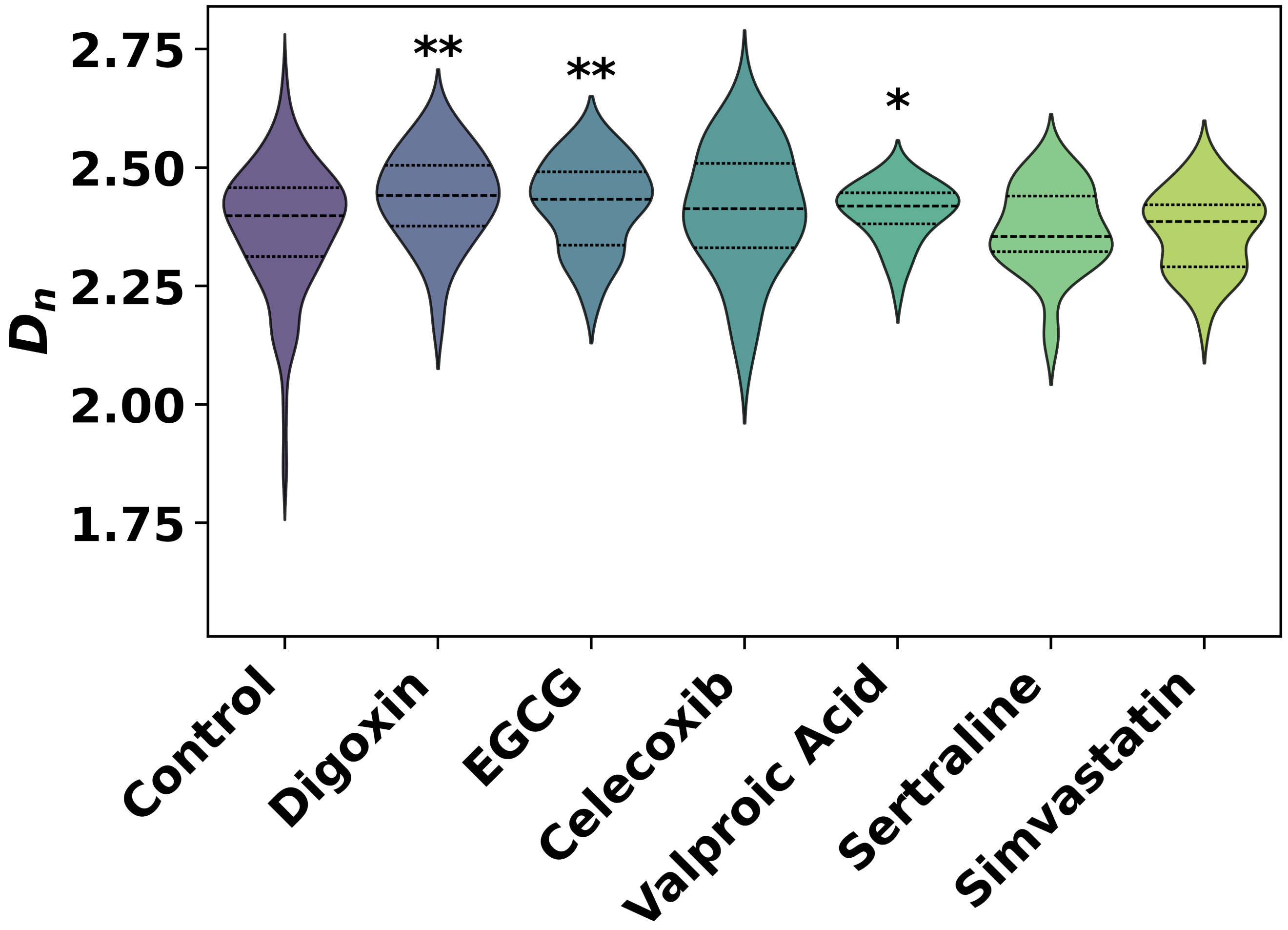


**Fig. 8.** TPR candidate drug screen results in live osteoblast cells. Violin plots show $D_n$ distributions for various treatments, including control and potential TPRs (digoxin, EGCG, celecoxib, valproic acid, sertraline, and simvastatin). Dashed lines within the violins represent the interquartile range. Significance levels compared to control: $^{*}P < 0.05$, $^{**}P < 0.01$, $^{***}P < 0.001$ (determined using unpaired t-test with unequal variance).

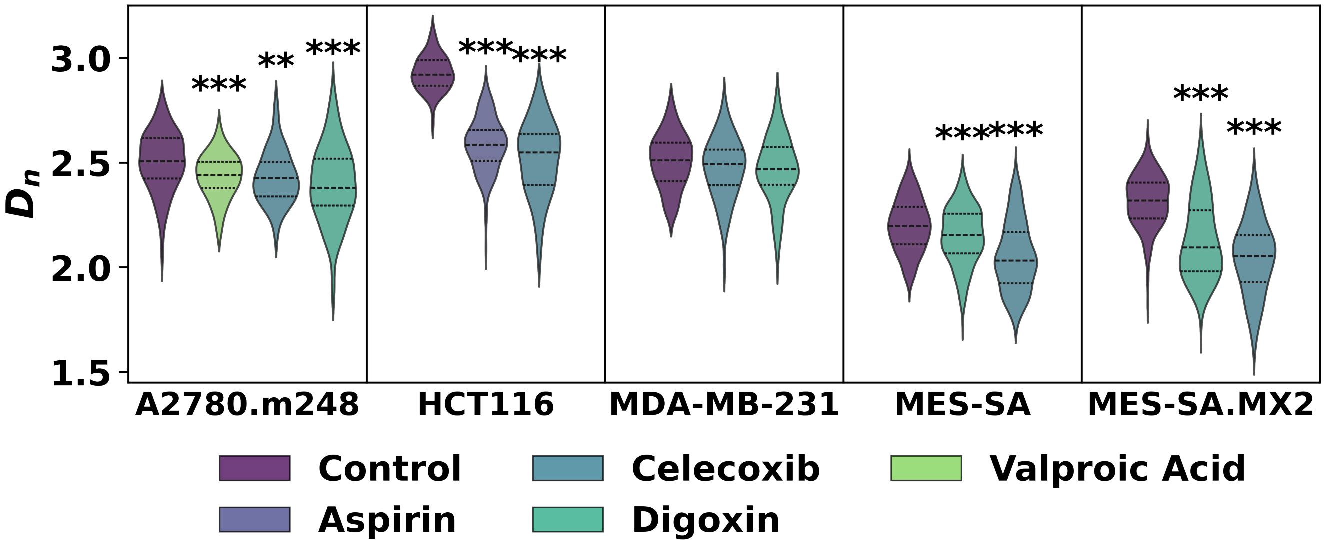


**Fig. 9.** Effects of Transcriptional Plasticity Regulators (TPRs) on $D_n$ across multiple cancer cell lines. Violin plots show the distribution of $D_n$ values for different cancer cell lines (A2780.m248, HCT116, MDA-MB-231, MES-SA, and MES-SA.MX2) treated with various TPRs (celecoxib, valproic acid, aspirin, and digoxin) compared to untreated controls. Cells were treated with TPRs for 30 minutes prior to PWS imaging. The width of each violin represents the frequency of $D_n$ values, while the internal lines indicate the quartiles. Asterisks denote statistical significance compared to the control group (*$P < 0.05$, **$P < 0.01$, ***$P < 0.001$; unpaired t-test with unequal variance).

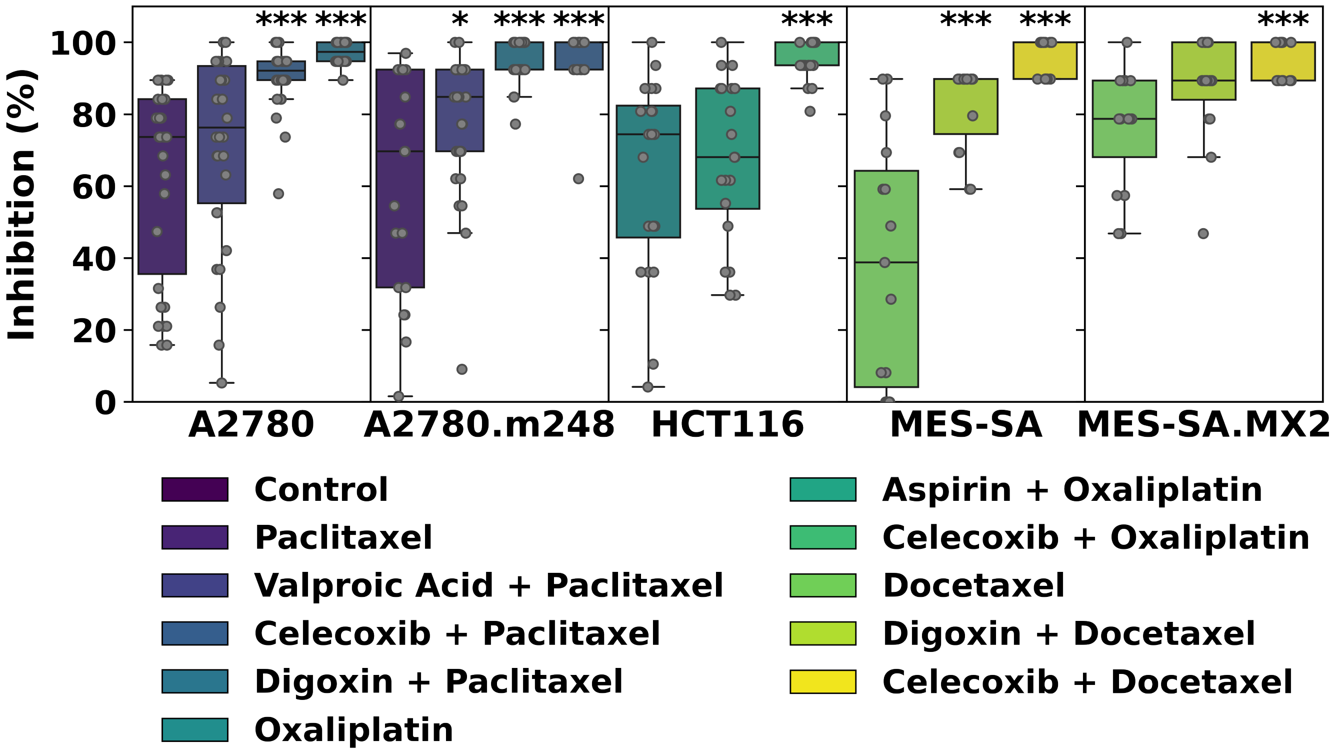


**Fig. 10.** Enhanced chemotherapeutic efficacy with TPR co-treatment across multiple cancer cell lines. Box plots show percent inhibition of cell growth for A2780, A2780.m248, HCT116, MES-SA, and MES-SA.MX2 cells treated with chemotherapy alone or in combination with TPRs. Treatments include paclitaxel, oxaliplatin, or docetaxel alone, and combinations with valproic acid, celecoxib, digoxin, or aspirin. Each data point represents an independent experiment. The box extends from the $25^{\text{th}}$ to $75^{\text{th}}$ percentiles, with the line in the middle representing the median. Whiskers show the minimum and maximum values. Significance levels compared to chemotherapy alone: $^{*}P < 0.05$, $^{**}P < 0.01$, $^{***}P < 0.001$ (determined using unpaired t-test with unequal variance).

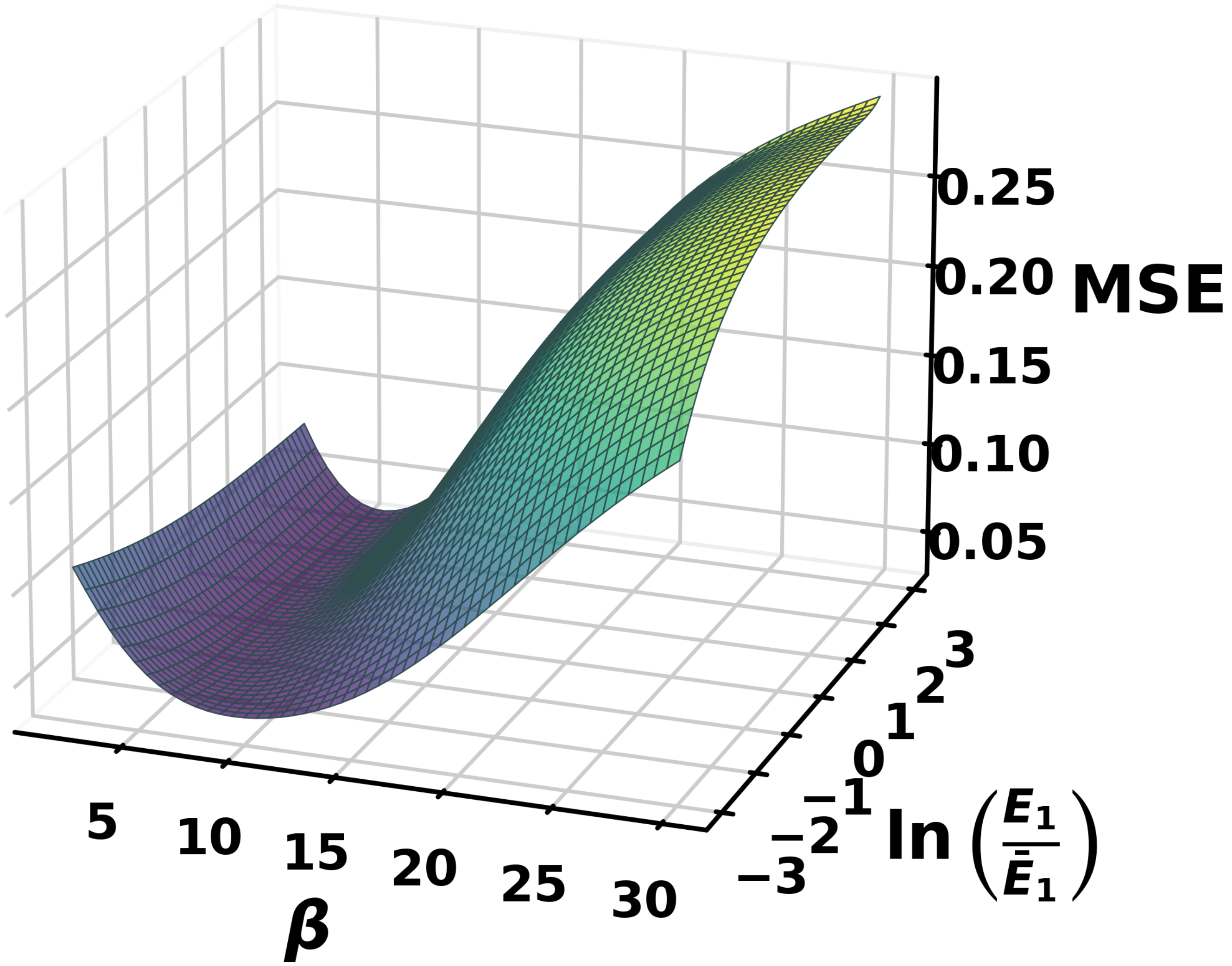


**Fig. 11.** Optimization of CDA model parameters for $\Theta_b(\Theta_a)$. Heat map showing the mean squared error (MSE) of the CDA model fit as a function of gene upregulation $\beta_a$ and initial relative expression rate $\ln(E_1/\overline{E}_1)$. The optimization was performed using experimental data from A2780 cells treated with paclitaxel and celecoxib. The color gradient represents MSE values, with darker colors indicating lower MSE and better model fit. The plot demonstrates that changes in $\beta_a$ most strongly influence the quality of fit to experimental data, with negligible effects of the initial relative expression rates.

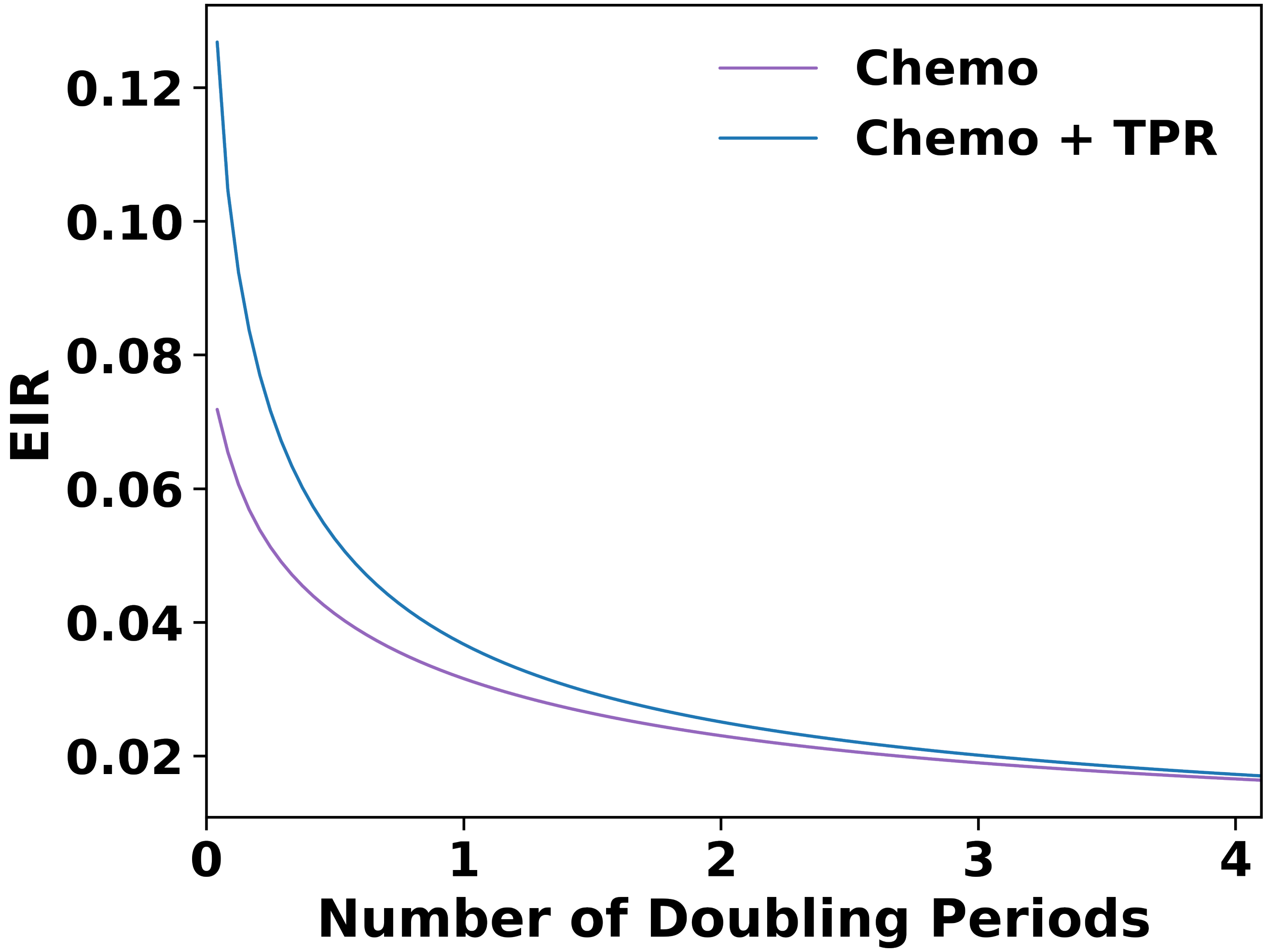


**Fig. 12.** CDA model predictions of Effective Inhibition Rate (EIR) using *in vitro* experiment results. The graph shows the predicted EIR as a function of the number of cell doubling periods for chemotherapy treatment alone (purple line) and chemotherapy combined with TPR (blue line). EIR represents the cumulative cancer cell death at a given time point.

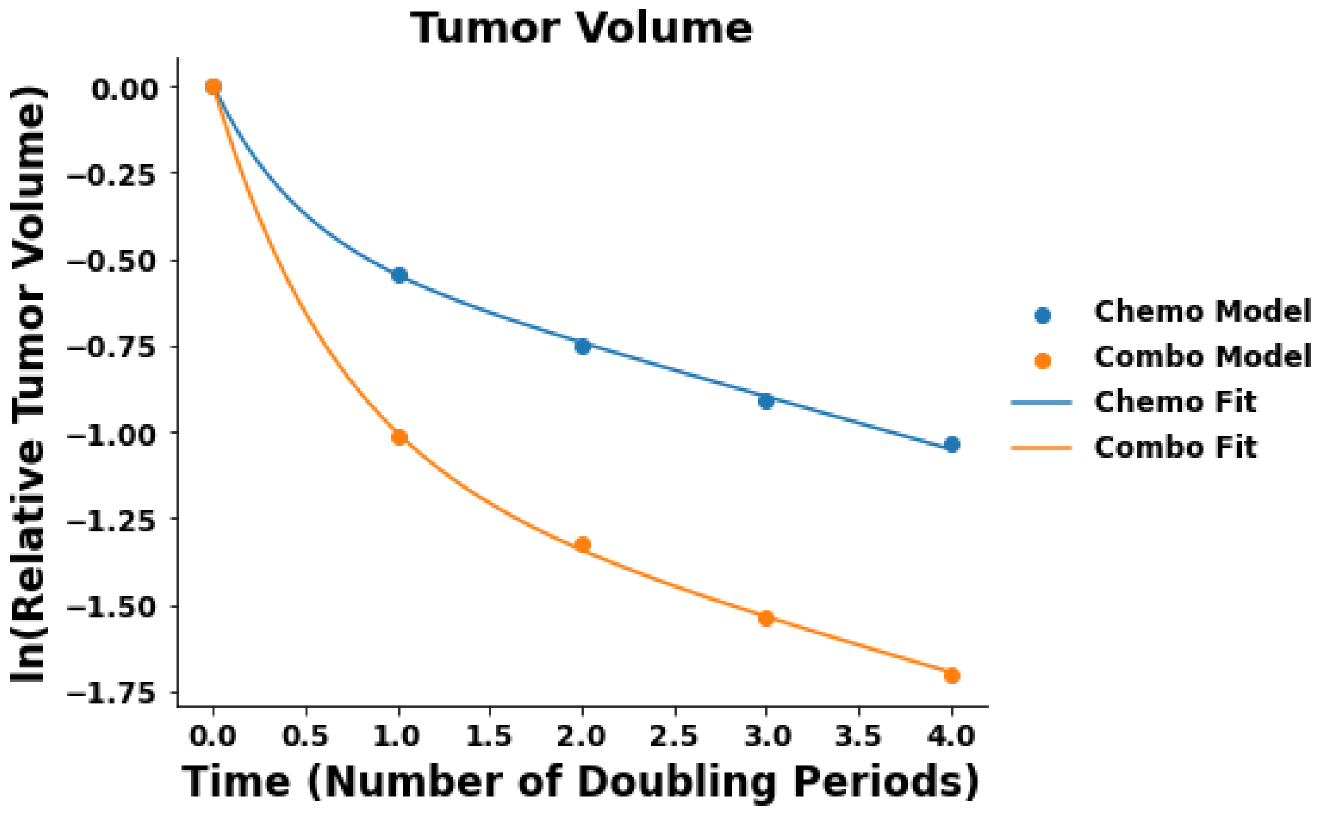


**Fig. 13.** CDA predictions for change in tumor volume using *in vitro* experiment results. The relative tumor volume was predicted based on a changed in $D_n$ for cells treated with chemotherapy alone (blue) vs. combination treatment of chemotherapy with a TPR (orange). CDA model predictions (dots) were fit with the adaptive inhibition model (Eq. 73; lines).

**Table 1.** Numerical values of Macromolecular Crowding (MC) model parameters

| Parameter | Description | Value for $\phi = 0$ |
|---|---|---|
| $V_{cell}$ | Volume of a typical HeLa cell | 500 $\mu m^3$ |
| $L_{DNA}$ | Number of DNA base pairs in a diploid human cell | $6 \times 10^9$ bp |
| $L_{DNA,half}$ | One-half of the total length of genomic DNA | 1 m |
| $l_{bp}$ | Length of one base pair | 0.34 nm |
| $N_{bp/turn}$ | Number of base pairs per turn of DNA | 10 |
| $r_{DNA}$ | Radius of the DNA molecule | 1 nm |
| $r_{TF}$ | Radius of TF (spherical approximation) | 4.0 nm |
| $r_{Pol\ II}$ | Radius of Pol II (spherical approximation) | 5.4 nm |
| $r_{crowder}$ | Radius of nuclear crowding agents | 3.0 nm |
| $D_{TF}$ | Diffusion coefficient of TF | 3 $\mu m^2/s$ |
| $D_{Pol\ II}$ | Diffusion coefficient of Pol II | 2 $\mu m^2/s$ |
| $D_{1,TF}$ | One-dimensional diffusion coefficient of TF on DNA | 0.046 $\mu m^2/s$ |
| $D_{1,Pol\ II}$ | One-dimensional diffusion coefficient of Pol II on DNA | 0.03 $\mu m^2/s$ |
| $k_t^{ns}$ | Association rate constant for nonspecific TF-DNA binding | $4.9 \times 10^4\ \mu M^{-1}s^{-1}$ |
| $k_f^{ns}$ | Association rate constant for nonspecific Pol II-DNA binding | $3.6 \times 10^4\ \mu M^{-1}s^{-1}$ |
| $k_o^{ns}$ | TF-DNA nonspecific dissociation rate | $4.9 \times 10^4\ s^{-1}$ |
| $k_b^{ns}$ | Pol II-DNA nonspecific dissociation rate | $3.6 \times 10^4\ s^{-1}$ |
| $K_{D,TF}^{ns}$ | Dissociation constant for nonspecific TF-DNA binding | 1 $\mu M$ |
| $K_{D,Pol\ II}^{ns}$ | Dissociation constant for nonspecific Pol II-DNA binding | 1 $\mu M$ |
| $k_t$ | Association rate constant for TF-promoter (O) binding | 0.05 $nM^{-1}s^{-1}$ |
| $k_f$ | Association rate constant for Pol II-Complex I binding | 0.03 $nM^{-1}s^{-1}$ |
| $k_o$ | TF-promoter (O) dissociation rate | 1.0 $s^{-1}$ |
| $k_b$ | Pol II-Complex I dissociation rate | 0.6 $s^{-1}$ |
| $K_{D,TF}$ | Dissociation constant for TF-O (promoter) binding | 1 nM |
| $K_{D,Pol\ II}$ | Dissociation constant for Pol II-O (promoter) binding | 1 nM |
| $k_m$ | Rate of pre-mRNA production | 0.008 $s^{-1}$ |
| $\gamma$ | Nuclear export rate of mRNA | $8 \times 10^{-4}\ s^{-1}$ |
| $\nu$ | mRNA degradation rate | $3 \times 10^{-4}\ s^{-1}$ |
| $[TF]_{tot}$ | Total concentration of TF | 30 nM |
| $[Pol\ II]_{tot}$ | Total concentration of Pol II | 30 nM |
| $[O]_{tot}$ | Total concentration of O (promoters) | 30 nM |
| $[D]_{tot}$ | Total concentration of DNA basepairs | 20 $\mu M$ |
| Coefficients for cubic fit to $f(\phi)$ from Brownian Dynamics simulations (6) | | |
| $\alpha_{TF}, \beta_{TF}, \gamma_{TF}$ | Coefficients for TF | -2.83, 3.87, -4.11 |
| $\alpha_{Pol\ II}, \beta_{Pol\ II}, \gamma_{Pol\ II}$ | Coefficients for Pol II | -3.89, 7.72, -7.72 |
| Coefficients for $\phi_{in}$-influenced free energies from Monte Carlo simulations (6) | | |
| $f_{cro,TF}(\phi)$ | TF crowding free energy | $-3.2\phi - 2.0\phi^2$ |
| $f_{cro,Pol\ II}(\phi)$ | Pol II crowding free energy | $-3.7\phi - 2.7\phi^2$ |
| $f_{cro,Pol\ II,s}(\phi)$ | Pol II sliding crowding free energy | $-2.6\phi - 4.6\phi^2$ |
| $f_{ba,TF}(\phi)$ | TF barrier free energy | $2.5\phi^2$ |
| $f_{ba,Pol\ II}(\phi)$ | Pol II barrier free energy | $3.1\phi^2$ |
| $f_{ba,Pol\ II,s}(\phi)$ | Pol II sliding barrier free energy | $0.1\phi^2 + 9.2\phi^3$ |

**Table 2.** Parameters used in the Chromatin-Dependent Adaptability (CDA) model

| Parameter | Description | Value |
|---|---|---|
| $l$ | Length along DNA of one basepair | 0.34 nm |
| $b$ | Radius of DNA molecule | 1 nm |
| $\xi$ | Characteristic distance between DNA strands | 35 nm |
| $L$ | One-half of total length of DNA in nucleus | 1 m |
| $T_{crit}$ | Critical time point for cell death decision | 7 hours |
| $\tau_{1/2}$ | Half-life of mRNA | 10 hours |
| $\phi_{in}$ | Volume fraction of crowders in nucleus | 0 – 0.5 |

**Table 3.** Mechanisms of action and effects on chromatin for chemotherapeutics

| Compound | Dose | Cell Line | Number of nuclei | Change in $D_n$ (%) | $P$ value | Mechanism of action |
|---|---|---|---|---|---|---|
| Control | – | A2780 | 332 | – | – | – |
| | – | A2780.m248 | 259 | – | – | – |
| | – | HCT116 | 262 | – | – | – |
| | – | MDA-MB-231 | 128 | – | – | – |
| | – | MES-SA | 265 | – | – | – |
| | – | MES-SA.MX2 | 203 | – | – | – |
| 5-fluorouracil | 0.5 µM | A2780 | 147 | +1.10 | $1.4 \times 10^{-13}$ | Inhibits thymidylate synthase; disrupts DNA |
| | 0.5 µM | A2780.m248 | 100 | +2.68 | $1.35 \times 10^{-5}$ | synthesis and repair |
| | 0.5 µM | MDA-MB-231 | 81 | +1.11 | $9.33 \times 10^{-2}$ | |
| Docetaxel | 5 nM | MES-SA | 194 | +1.19 | $4.19 \times 10^{-2}$ | Binds to $\beta$-tubulin; stabilizes microtubules; in- |
| | 5 nM | MES-SA.MX2 | 82 | +6.36 | $8.01 \times 10^{-17}$ | duces mitotic arrest |
| Gemcitabine | 50 nM | MES-SA | 101 | +4.82 | $9.68 \times 10^{-15}$ | Inhibits DNA polymerase; terminates DNA |
| | 50 nM | MES-SA.MX2 | 69 | +1.07 | $1.03 \times 10^{-1}$ | chain elongation |
| Oxaliplatin | 5 µM | A2780 | 101 | +8.04 | $8.21 \times 10^{-36}$ | Forms DNA adducts; induces DNA damage |
| | 5 µM | A2780.m248 | 85 | +5.36 | $5.33 \times 10^{-16}$ | and apoptosis |
| | 5 µM | MDA-MB-231 | 59 | +3.15 | $8.16 \times 10^{-5}$ | |
| | 15 µM | HCT116 | 289 | +8.29 | $8.93 \times 10^{-57}$ | |
| Paclitaxel | 5 nM | A2780 | 99 | +4.50 | $4.06 \times 10^{-10}$ | Binds to $\beta$-tubulin; stabilizes microtubules; |
| | 5 nM | A2780.m248 | 45 | +5.86 | $6.58 \times 10^{-9}$ | blocks cell cycle progression |
| | 5 nM | MDA-MB-231 | 36 | +4.35 | $7.99 \times 10^{-5}$ | |

**Table 4.** Mechanisms of action and effects on chromatin for candidate TPRs in A2780 cells

| Compound | Dose | Number of nuclei | Change in $D_n$ (%) | $P$ value | Mechanism of action |
|---|---|---|---|---|---|
| UNC1999 | 100 nM | 158 | -1.96 | $5.6 \times 10^{-4}$ | Inhibits H3K27 methyltransferases EZH1/2 |
| Metoprolol | 200 µM | 111 | -1.84 | $6.3 \times 10^{-3}$ | $\beta$-1-adrenergic receptor antagonist |
| EGCG | 25 nM | 276 | -1.77 | $8.8 \times 10^{-5}$ | Inhibits DNMT, HDAC1, and HDAC3 |
| UNC0638 | 100 nM | 158 | -1.96 | $5.6 \times 10^{-4}$ | Inhibits H3K9 methyltransferases G9a and GLP |
| Propranolol | 200 µM | 111 | -1.84 | $6.3 \times 10^{-3}$ | Nonselective $\beta$-adrenergic receptor antagonist; dephosphorylates histone H3 |
| Simvastatin | 10 µM | 261 | -2.05 | $7.1 \times 10^{-6}$ | Inhibits HMG-CoA reductase and HDAC1/2 |
| Resveratrol | 35 µM | 271 | -2.05 | $1.3 \times 10^{-5}$ | Inhibits COX-1/2 and HDAC1-11 |
| Valproic acid | 100 µM | 117 | -3.68 | $3.1 \times 10^{-10}$ | Blocks voltage-gated ion channels; inhibits HDACs |
| Sertraline | 10 µM | 157 | -3.98 | $2.2 \times 10^{-11}$ | Inhibits serotonin reuptake |
| Digoxin | 150 nM | 130 | -5.79 | $1.7 \times 10^{-24}$ | Cardiac glycoside; inhibits $Na^+$/$K^+$-ATPase, increasing intracellular $Ca^{2+}$ |
| Celecoxib | 75 µM | 132 | -7.38 | $3.2 \times 10^{-25}$ | Antiinflammatory; inhibits voltage-gated $Na^+$, $Ca^{2+}$, and $K^+$ channels; COX-2 inhibitor |
